# Self-Healable Spider Dragline Silk Materials

**DOI:** 10.1101/2023.04.01.535237

**Authors:** Wen-Chia Chen, Ruei-Ci Wang, Sheng-Kai Yu, Jheng-Liang Chen, Yu-Han Kao, Tzi-Yuan Wang, Po-Ya Chang, Hwo-Shuenn Sheu, Ssu Ching Chen, Wei-Ren Liu, Ta-I Yang, Hsuan-Chen Wu

**Affiliations:** Department of Biochemical Science and Technology, National Taiwan University, Taipei City, 10617, Taiwan; Genomics Research Center, Academia Sinica, Taipei City, 11529, Taiwan; National Synchrotron Radiation Research Center, Hsinchu City, 30076, Taiwan; Department of Chemical Engineering, Chung Yuan Christian University, Taoyuan City, 32023, Taiwan

**Keywords:** self-healing, spider silk, *Nephila pilipes*, graphene, wearable electronics

## Abstract

Developing materials with structural flexibility that permits self-repair in response to external disturbances remains challenging. Spider silk, which combines an exceptional blend of strength and pliability in nature, serves as an ideal dynamic model for adaptive performance design. In this work, a novel self-healing material is generated using spider silk. Dragline silk from spider *Nephila pilipes* is demonstrated with extraordinary *in situ* self-repair property through a constructed thin film format, surpassing that of two other silks from spider *Cyrtophora moluccensis* and silkworm *Bombyx mori*. Subsequently, R2, a key spidroin associated with self-healing, is biosynthesized, with validated cohesiveness. R2 is further programmed with tunable healability (permanent and reversible) and conductivity (graphene doping; R2G) for electronics applications. In the first demonstration, film strips from R2 and R2G are woven manually into multidimensional (1D-3D) conductive fabrics for creating repairable logic gate circuits. In the second example, a reversibly-healable R2/R2G strip is fabricated as a re-configurable wearable ring probe to fit fingertips of varying widths while retaining its detecting capabilities. Such prototype displays a unique conformable wearable technology. Last, the remarkable finding of self-healing in spider silk could offer a new material paradigm for developing future adaptive biomaterials with tailored performance and environmental sustainability.

## Introduction

The technical advancements in the electronics sector during the last half-century have had profound impacts on global economies and societies [1]. The extensive spectrum of applications includes computing, communication, robotics, manufacturing, energy, and even biomedicine [2-5]. In addition, one growing need for high-performance electronics with distinctive features for mobility and adaptability has expedited the development of next-generation technologies capable of converting rigid silicon-based devices into conformable electronics [6, 7]. To enable the so-called “soft” electronics technology, mechanically malleable and stretchable materials are combined with electrical components or conductive fillers [8-10]. Several petroleum-based chemically-synthesized elastomers (PVA, PET, PDMS, PI, and PU) have been intensively explored as elastic matrices in the production of soft electronics with remarkable success [11].

In addition, a newly engineered attribute of elastomer materials, self-healing, has been granted as a unique and favorable function for soft electronics and mechanics [12-14]. Self-healing of materials refers to the inbuilt capacity to undertake structural renewal after impairments [15]. It can significantly reinforce the environmental adaptability and robustness of materials. Numerous repairable material strategies have been innovated, including *in situ* polymerization [16, 17], irreversible chemical reactions[18], and reversible dynamic interactions, such as hydrogen bonding [19], electrostatic interactions[20], disulfide bonding [21], and Diels–Alder reactions[22, 23]. The engineered self-healable electronics can not only actively restore the pre-failed structures but also proactively halt the possible damage propagation, enabling to retain their desirable performance and key functionality upon significant external disturbance [24, 25]. Therefore, self-repairable soft devices have long been envisioned for vast applications in future biosensors, wearable electronics, and medical equipment [26].

In Nature, biology offers enormous exciting examples of advanced self-healing soft materials technology and associated biomanufacturing systems that can be harnessed for fabricating future bioinspired devices. For instance, naturally derived biological materials, such as cationic polysaccharide chitosan [27], anionic polysaccharide alginate [28], and polypeptide gelatin [29] have been successfully demonstrated for specialized hydrogel based self-healable sensors/devices. Another unique polypeptide material, extracted from squid ring teeth (SRT), is shown to possess extraordinary intrinsic self-healing abilities [30]. Additionally, recombinant SRT inspired proteins have been biosynthesized and fabricated into configurable and self-restorative soft actuators. Many forms of bioinspired demonstrations provide ideal and innovative models into harnessing the power of biological design, synthesis, and fabrication for the manufacturing of next generation self-healable soft materials and devices in light of future sustainability and eco-friendliness [31-34].

Analogous to the structural hierarchy of SRT, another renowned biomaterial spider silk also possesses an interconnected molecular architecture comprising of both rigid and elastic domains [35, 36]. Unlike the polypeptide sequences of SRT, spider dragline silk possesses a unique molecular design of poly-alanine associated β-sheet domains and proline-associated β-spiral motifs that are accountable for rigid and flexible nanostructures, respectively [36-38]. These nanostructures, when perfectly orchestrated, would confer the ultimate mechanical performance of spider silk materials [39]. Amid the enormous material traits of spider silk, like low weight [40], strength [41], flexibility [42], toughness [43], supercontraction [44] and biocompatibility [45], yet its self-healing potential has not been fully explored. From the biomimicry perspective, it’s worthwhile deciphering whether the silk materials from spiders also carry the intrinsically-repairable characteristic. Furthermore, the model of spider silk material might offer a different dimension, structural and dynamic, in understanding and utilizing such complex molecular healing phenomena. This, in turn, can further advance future bioengineered materials or functional biodevices.

Herein, we used the film configuration to explore the self-healing potential of dragline silk generated by the main ampullate (MA) glands of the golden orb weaver *Nephila pilipes* (**Fig. 1**). With demonstrating the excellent *in situ* self-restoration capability of the composite MA silk collected form *N. pilipes*, we further investigated and recapacitated the healable silk systems via biosynthetic MA spidroins R1 and R2, resembling MaSp1 and MaSp2 respectively. Amazingly, the biomimetic silk variant R2 exhibited distinctly superior repairing characteristic than R1. Additionally, biosynthetic R2 was tuned with an adjustable healing capability. Next, we doped electrically-conductive graphene into the biosynthetic R2, rendering a multifunctional silk material R2G with conductivity and healability. We used R2G to fabricate flexible 3D logic gate devices, which were able to maintain signaling functionality even after being cut and repaired. Last, a flexible finger ring was fabricated with a reversibly-healable R2 and R2G, conferring a cyclic tunable ring configuration to accommodate fingers with varied sizes. By coupling the R2 ring with the designated electronic device, a smart healable finger recognition system was achieved.

**Figure 1.**
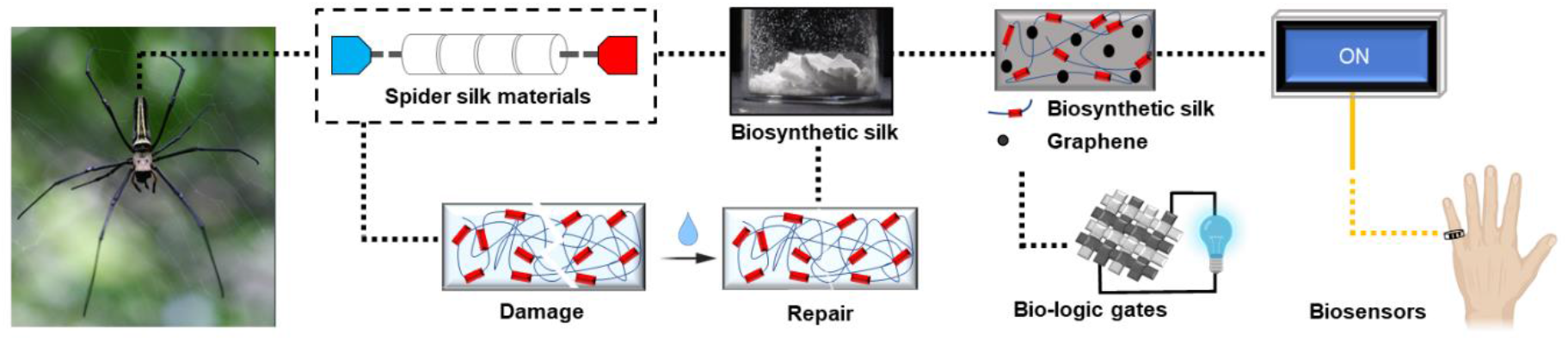
The schematic of self-healable spider silk materials. Dragline silk produced by *N. pilipes* was shown to have self-healable potential. Biosynthetic recombinant spidroin was used to further validate its self-healability. Coupling the self-healability of biosynthetic spidroin with electrical conductivity (graphene inclusion), the self-healing conductive biosynthetic silk material was developed with applications in logic gate circuits and wearable devices.

## Materials and Methods

### Native silk sample collection and processing

Silk collection process was adopted from Wu et al (2017) [46]. Briefly, female *Nephila pilipes* and *Cyrtophora moluccensis* were gently fixed on polystyrene form plates. MA dragline silks were pulled directly from the spinnerets of immobilized spiders and collected by a motorized rotor under a dissecting microscope, followed by the silk collection at the reeling speed of ∼0.5-1 m min^-1^. The obtained spider silk samples were stored for further experiments. Additionally, *Bombyx mori* cocoons were degummed by 0.02M Na_2_CO_3_ at 80 °C for 1h. After rinsing and drying, the degummed cocoons dissolved in 9.3M LiBr at 60 °C for 4h, dialyzed in DI water, and lyophilized to obtain silk fibroin power for subsequent experiments [47].

### Silk film preparation

Hexafluoroisopropanol (HFIP; Sigma-Aldrich) was used to dissolve silk samples at the concentration of 2.5 or 5%. After complete dissolution, we cast the clear solution in a Teflon tray and dried it overnight in the chemical hood (∼25 °C, 60% RH) to obtain the silk thin films.

### *In situ* crack and repair assessment of silk films

We used a razor blade to generate a minor crack at the thickness of 30 μm on the silk films. Subsequently, a drop of water (∼0.5-2 μL) was introduced to the scar area and dried out the films at ambient temperature. Afterward, the repaired films were then imaged under an optical microscope, followed by image analysis to evaluate the thickness of the remaining scar.

Additionally, we further employed a tensile test to briefly access the restoration of *N. pilipes* and *C. moluccensis* films. First, the silk films were secured to the gripper of the tensile tester. A metal bar with a round tip that was premounted to a syringe pump was positioned perpendicular to the films. The syringe pump propelled the metal bar toward the films at a velocity of 0.2 mm sec^-1^. Utilizing a digital video camera, the dynamics of film distortion and rod contact resistance were documented. Afterwards, the video footage was then processed to calculate the bar migration distance and angle of film deformation.

### MA gland RNA library construction and sequencing of *N. pilipes*

MA glands of female *N. pilipes* were isolated according to the procedures from Hwang et al (2005) [48] and Lewis et al (1992) [41]. Briefly, spiders were first depleted of MA silk by forced-silking to promote RNA expression inside the MA gland. The silked spiders were subsequently sacrificed, and MA glands were dissected under a dissecting microscope. The RNA extraction was immediately performed using RNeasy Mini Kit (Qiagen, USA), followed by mRNA library construction via TruSeq Stranded mRNA Library Prep Kit (Illumina, USA), and Illumina NovaSeq 6000 sequencing (Genomics BioSci & Tech Co., Taiwan). The raw reads were deposited to NCBI SRA as part of the BioProject PRJNA872937 and are available under accession number SRR21615954 (silked) and SRR21615955 (non-silked).

### Transcriptomic analysis and spidroin-like candidate identification

104,268,750 trimmed Illumina pair-end RNA reads were *de novo* assembled using Trinity v2.3.2 [49]. The assembled transcripts were blasted against 594 Araneoidea spidroin related gene candidates and 134 *Nephila clavipes* spidroin proteins for open reading frames (ORFs) annotation. In addition, 36 N-terminal domains (NTD) and 74 C-terminal domains (CTD) sequences were found to be corelated to NTD/CTD Sanger sequencing results. Transcripts with hitted length over 80 amino acids were selected as spidroin-like seeds for further annotation **(Supplementary Table_contig)**. Candidates with poly-alanine (A_n_, n=3-14) and GPGXX (MaSp2 motifs) or GGXGXXGGX (MaSp1 motifs) were further selected as represented sequences. The chosen contigs were displayed in **Supplementary Fig. S1**

### Biosynthetic spidroin gene construction

The *E. coli* codon-optimized monomeric r1 (MaSp1 associated) and r2 (MaSp2 associated) genes were constructed utilizing overlap PCR and cloned into pET28a derivative vectors. The amplification of repetitive units was generated by BioBrick restriction enzyme assembly using NheI and SpeI enzyme sets (New England BioLabs, USA). Subsequently, both NTD and CTD were amplified using overlap PCR and were cloned into the repetitive units containing vectors by NdeI/NheI and SpeI/HindIII restriction cloning, respectively. Following the cloning procedures, plasmids pENTH-NTD-(r1)_32_-CTD and pENTH-NTD-(r2)_32_-CTD, each containing 32 copies of r1 and r2, were constructed and designated as R1 and R2, accordingly. The plasmids were electroporated into *E. coli* BLR(DE3)Δ*endA* strain for further biosynthetic spidroin expression. Additional plasmid maps were listed in **Supplementary Fig. S3**.

### Biosynthetic spidroin protein production and purification

The transformant cells were inoculated in a 5 L benchtop fermenter (Major Science, Taiwan) for biosynthetic spidroin synthesis. Specifically, Terrific Broth (17mM K_2_HPO_4_, 72mM KH_2_PO_4_, 23.6g L^-1^ yeast extract, 11.8g L^-1^ Tryptone, 0.4% glycerol, 0.01% antifoam) with 50 μg mL^-1^ kanamycin and additional 50 μg/mL ampicillin (for R2 fermentation) was formulated as the growth medium. Additionally, a dissolved oxygen (DO) cascade control was set at 50% to sustain the cell growth at 30°C fermentation, followed by a supplementation of substrate feeding. Subsequently, cells at OD_600_ = 25 were induced by 0.5 mM IPTG and for a 24-hour spidroin production at 20 °C. After fermentation, cells were harvested and kept at −80 °C for storage.

For spidroin purification, the harvested cells were resuspended in lysis buffer (10 mM Tris-HCl pH8.0, 1.25 mg mL^-1^ Lysozyme, and 0.5% Triton X-100), followed by ultrasonication process for cell lysis on ice. After sonication, the pellet fraction was centrifuged, resuspended in washing buffer (5% SDS, 0.5% Triton X-100, and 10mM Tris-HCl pH8.0). Following a second centrifugation, the pellet fraction was recovered and washed four times with DI water and remaining impurities were eliminated. After cleaning-up, the silk product was treated with liquid nitrogen and was lyophilized by a vacuum freeze dryer.

### Nanostructural characterization of purified spidroin

The supramolecular interaction of purified biosynthetic silk protein was investigated through immunofluorescence staining and dynamic light scattering (DLS). First, the purified silk products of R1 and R2 (not lyophilized) were stained with our proprietary mouse anti-NTD antibody (**Supplementary Fig. S4**) developed in this study. The silk products were rinsed and subsequently stained with CF555-labeled goat anti-mouse IgG (H+L) (Biotium, USA). The resultant silk products were observed using a fluorescence microscope. Additionally, we also employed DLS to access the particle size and dispersion of the silk products (without lyophilization). NanoBrook 90Plus particle size analyzer (Brookhaven Instrument Corp., USA) was utilized to gather the data for both R1 and R2 for subsequent comparison.

### Protein analysis of purified spidroins

Purified R1 and R2 spidroins were analyzed by SDS-PAGE and Western blotting. Briefly, samples premixed with LDS sampling buffer were loaded to an 8% sodium dodecyl sulfate-polyacrylamide gel and run in MOPS buffer. Afterward, the gel was stained with Coomassie Blue R-250 and subsequently destained for further evaluation. In Western blotting analysis, the SDS-PAGE gel was transferred to a PVDF membrane at 400 mA current for 1.5h on ice. The transferred membrane was then blocked with Gelatin-NET solution, followed by mouse anti-NTD staining, rinsing, and HRP-labeled goat anti-mouse (PerkinElmer Life Science, USA) staining. The PVDF membrane was detected using chemiluminescence ECL kit (PerkinElmer Life Science, USA) and documented by BioSpectrum^®^ 810 Imaging Systems (Analytik Jena US LLC, USA).

### Post treatments of biosynthetic silk films

Two post-treatments were performed to treat as-cast biosynthetic silk films: alcohol immersion and water vapor annealing (WVA) [50-53]. For alcohol treatment, the films were immersed in various concentrations of ethanol solutions for 1 hour. After ethanol removal and water rinsing, the treated films were placed onto a petri dish for drying at the ambient environment.

For WVA treatment, the silk films were placed on a petri dish without covering and incubated in a prewarmed water bath at either 37 °C or 60 °C for 1-2h, without direct contact with bulk water. Subsequently, the petri dish was removed from the water bath and the silk films were dried at the ambient temperature.

### *In situ* cut-and-healing treatment of silk films

For biosynthetic film healing test, silk films were first cut into two half slices by a razor blade. Two halves of the silk films were surface-moisturized with DI water (∼0.5-2 μL) and joined with a minimum processable overlap region of 2.5 mm x 1.5 mm. The healed silk films were dried at ambient condition for subsequent experiments.

### Tensile measurements of silk films

First, the film thickness was evaluated by a digital micrometer. Next, the tensile properties of silk films were measured by Shimadzu EZ-SX (Shimadzu Co., Japan). The silk films at the dimension of 30 mm x 1.5 mm were clamped by the sample grips on a tensile tester, with the initial gauge length of 10 mm. Upon the tensile measurement, the film samples were extended at the speed of 5 mm min^-1^ and the stress-strain characteristics were obtained.

### X-ray diffraction analysis of silk films

The crystalline structure of silk sample was studied using X-ray diffraction performed on the TLS 01C2 beamline at National Synchrotron Radiation Research Center (NSRRC), Taiwan. The wavelength of the X-ray beamline was 0.1 nm, with a fixed energy of 12 keV. After measurements, one-dimensional diffraction profiles were generated from the raw XRD data using Fit2D software, with 2θ spanning from 6° to 20°. The d-spacings of 0.84, 0.52, 0.43, 0.36 nm [54-56] were associated with the β-sheet nano-crystallines in the silk samples. The degree of crystallinity was defined as the integrated intensity ratio of crystal peaks to the sum of crystal peaks and the amorphous halo by Gaussian deconvolution using Origin 9.0 (OriginLab, USA).

### Fourier transform infrared study of silk films

Attenuated total reflection-Fourier transform infrared (ATR-FTIR) was utilized to profile the silk films. The FTIR spectrum of silk samples was acquired via FTIR-4100 spectroscopy with ATR PRO 410-A accessory (JASCO, USA). The silk samples were identified and the resulting Amide I region (wavenumber between 1600 and 1700 cm^-1^) from the FTIR data was selected for Gaussian deconvolution and further secondary structures determination.

### Resistance of silk films to water

To gauge the silk films durability against water, the mass loss of silk samples was measured after incubation in water. Briefly, silk films with equal mass and post-treatment history were created. One silk film was dried overnight at 60 °C (control), while another was treated with three cycles of 5-minute incubation in DI water before being dried overnight at 60 °C (test). Accordingly, the collected weights obtained from each film were recorded. The weight retention (or resistance) of the silk film was assessed by comparing the residual dry mass of the test silk film to the control.

### Graphene-silk composite films and logic-gated electrical circuit

The biosynthetic spidroin R2 was blended with graphene, provided by Dr. Wei-Ren Liu [57], yielding a graphene-silk composite, R2G. Specifically, the weight ratio of graphene to R2 was 0.15 for a total graphene-silk concentration of 5% in HFIP. After The graphene-silk solution was cast overnight at room temperature onto a Teflon tray to produce R2G thin films. To fabricate the biosynthetic silk-based logic gates, R2G and R2 strips (30 mm x 1.5 mm) were used as the building blocks for manually weaving into 2D fabrics or 3D-tublar fabrics. The interwoven fabric structures were then treated with WVA at 37 °C for 2 hours to firmly connect the R2 and R2G areas. In addition, 3D-tubular constructions were generated by bending 2D-fabrics and self-healing the head-to-tail section. Afterwards, all 1D to 3D silk-fabrics were cut in half and self-healed to create patched fabric structures. These self-patched silk fabrics were subsequently connected to a pre-mounted, incomplete electrical circuit consisting of a battery, a blue LED, a power switch, and an open break that rendered the circuit unfinished. After inserting the self-patched silk fabrics into the circuit by bridging the open break, the resulting change in circuit completeness and LED lighting was captured by a digital camera.

### Reversible healing of silk films and reconfigurable silk rings

Biosynthetic R2 films (30 mm x 1.5 mm) were treated with 60 °C WVA for 1 hour and dried at room temperature. The treated R2 films were then cut in half, moistened with DI water (∼0.5-2 μL), adhered, and dried at room temperature for 30 minutes. Then, we introduced a greater volume of water (>10 μL) to the healed regions of the patched silk films in and then gently withdrew them by hand, resulting in a complete separation of the film parts. In addition, we evaluated the possibility of re-healing the halved silk films at the same healing regions. Multiple cycles of reversible healing-separation were performed, and the tensile properties of the re-healed films were compared. To create an adjustable ring structure, a long silk strip (60 mm x 3 mm) was treated with 60 °C WVA for 1 hour, and then one end was attached to the strip’s body to form a circular ring with the aid of surface-moisturization of water. The pre-formed ring was easily reopened by applying a higher amount of water (>20 μL) to the patched portion of the strip. With repetitive opening and reforming, a variety of ring sizes were created on the same silk strip.

### Graphene-silk ring-based resistance measurement and finger recognition device

The silk ring was fashioned as a sensing probe for the purpose of gauging the electrical resistance of human fingertips. Briefly, the ring probe was composed of a self-healing R2 strip with R2G components coupled to it. Two enameled wires were attached to a Keithley 2400 source meter (Tektronix, USA) and plugged into the R2G electrodes of the R2 ring strip. The system’s detection range was determined by testing several conductive and non-conductive reference materials. Subsequently, 7 participants with 70 finger resistance data points were measured for further analysis.

The prototype finger identification device was constructed using a silk-ring finger detecting probe, an Arduino Uno microcontroller board, and an LCD display. Additional Arduino sketch code was shown in **Supplementary Fig. S13**, and the detailed assembly configuration was presented in **Supplementary Fig. S14.** The effectiveness of the identification device was evaluated using the adjustable silk ring probe on test participants.

### Results and Discussion

Whereas research into chemically synthesized self-repairing materials is extensive, the development of natural biomaterials that procure unique healability is still in its infancy. Recently, two forms of biopolymers derived from SRT and silkworm cocoons [47, 58] have been found to perform solvent-assisted repairability upon damage. From the biomimicry perspective, it’s worthwhile deciphering whether the silk materials from spiders also carry the intrinsically-repairable characteristic. Furthermore, the model of spider silk material might offer a different dimension, structural and dynamic, in understanding and utilizing such complex molecular healing phenomena.

### Repairability of natural silk materials

In this research, we aimed to unravel the material healing capability of spider silk. Particularly, MA dragline silks collected from two eco-types of spiders, *Nephila pilipes* and *Cyrtophora moluccensis*, were both explored for their potential in material healability. In comparison, *N. pilipes*, a golden orb weaver, builds an elastic 2D web, and *C. moluccensis*, a tent-web builder, constructs a rigid 3D dome web for foraging and protection (**Fig. 2A**). It’s therefore noteworthy to study the molecular healability as a material indicator for the diverse forms of spider silks.

**Figure 2.**
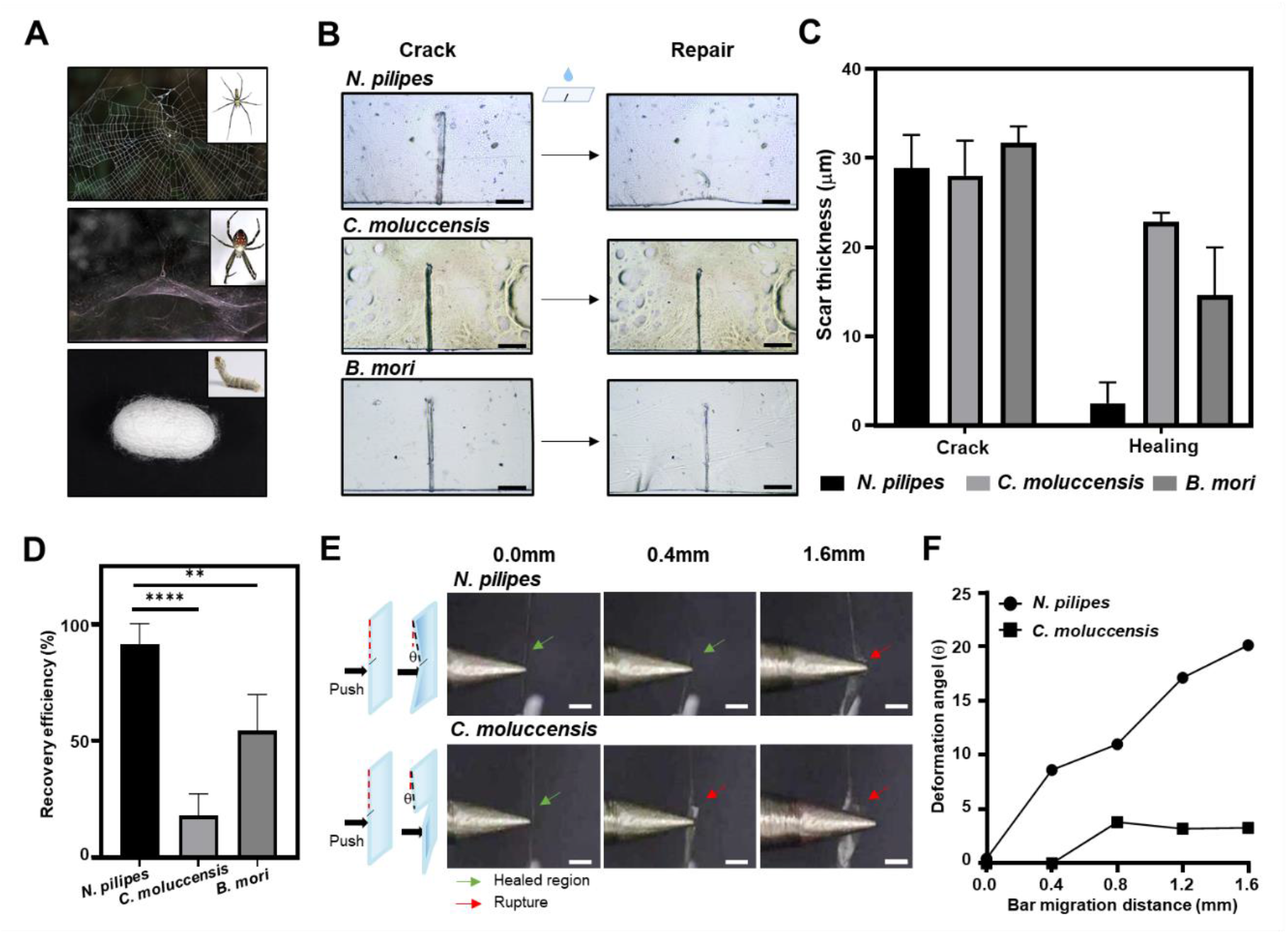
Characterization of self-healability of natural silk materials. (A) Silk samples collection from various species: *N. pilipes, C. moluccensis*, and *B. mori*. (B) Crack-and-repair examinations of the film-formatted natural silks. A crack in the films was gently introduced by a razor blade and moistened with water for repair observation. Scale bar: 200μm. (C) Quantitative measurement of the repair process based on the scar thickness on the films. Using imaging analysis, the pre- and post-repair scar thicknesses of silk films were measured and displayed. (D) Repair efficiency of the silk films. The degree of scar reduction in films was calculated by normalizing the scar thickness difference to the original scar thickness. (n=5; ** p<0.01, **** p<0.0001) (E) The tensile force-based resistance evaluation for the self-healing ability of silk films. The cut-and-healed films from both *N. pilipes* and *C. moluccensis* were challenged with a round-tipped metal bar recorded in a video. The film of *C. moluccensis* was resistant to bar migration to 0.4 mm until the healed region fractured, whereas the film of *N. pilipes* was more resistant up to 1.6 mm. Scale bar: 1mm. (F) The deformation angle of the films upon bar migration. The film distortion angle was determined from the video footage and represented on the plot.

Briefly, female spiders from both species were gently-immobilized, followed by the MA dragline fibers collection by a motorized rotor. Subsequently, the MA silk samples were dissolved in HFIP and cast in a Teflon tray to fabricate thin films. Similarly, the silk cocoons harvested from silkworm *Bombyx mori* (**Fig. 2A bottom**) were also processed into thin films, serving as a reference control for further healing tests.

The as-cast films, two from the spiders and one from the silkworm, were challenged for film healing capacity. First, we utilized a razor blade to generate a minor scratch with ∼30 μm width to all of the film samples (**Fig. 2B left**). Subsequently, a drop of DI water (∼0.5-2 μL) was introduced to the crack region of the films. After incubating for 20 minutes at ambient conditions (∼25°C, 60% RH), the water-treated silk films were monitored again under the microscope to evaluate the progression of film recovery. As indicated in **Fig. 2B Right**, differential recovery of the crack mark was observed among the film samples. The scar on the silk films of *N. pilipes* diminished significantly after healing. In contrast, the residual scars on the treated films of *B. mori* and *C. moluccensis* remained clearly discernible.

For a more quantitative comparison regarding the healing capacity of the silk films, we measured the width of the scar marks, before and after healing, on the silk films under the microscope. As the results depicted in **Figs. 2C & D**, the initial cracks generated among the silk films was at 28∼31 μm (no statistical difference). All of the silk films were repairable after the supplementation of water, with scar reduction to 2.4 μm for *N. pilipes*, 22.8 μm for *C. molecensus*, and 14.6 μm for *B. mori*, respectively in **Fig. 2C**. Last, the overall healing capacity of each silk film was obtained by calculating the ratio of scar reduction to the initial cut (**Fig. 2D**). The statistical results indicated distinct healability among the silk films. That is the silk film from *N. pilipes* recovered the most (∼ 91%); *B. mori*’s silk film exhibited moderate recovery capacity (∼54%); nevertheless, the silk film of *C. molecensis* displayed the least repairing performance (∼18%). Compared to *C. molecensis* silk film and *B. mori*’s silk film, the silk film from *N. pilipes* demonstrated the best healability (****p<0.0001 and **p<0.01 respectively).

In addition to the morphological repair assay abovementioned, a force-based approach was alternatively utilized as a complementary repair index for the silk materials. Silk films from *N. pilipes* and *C. moluccensis* were processed by the same healing procedures (**Fig. 2B**), including cutting and repairing. Subsequently, a metal bar controlled by a syringe pump was gradually exerting a tensile force onto the healed silk films (at speed of 0.2 mm sec^-1^) and the film deformation was recorded by a digital camera. As shown in **Fig. 2E left**, the silk films were consequentially deformed upon the application of the tensile force by the motorized metal bar. Aligned with the axial direction of the bar, the film was pushed and deformed over time until it fractured at the healed region.

The strength of the silk film was greatly determined by the repair efficiency of its cut we introduced, as a fact that the weakness of the healed films is significantly governed by the presence of exterior structural defects. In **Fig. 2E right**, the healed film from *N. pilipes* was able to withstand the bar-induced tension further (1.6 mm until fracture at the healed site), as compared to the healed film from *C. moluccensis* (0.4 mm; fracture upon bar contact). This is indicative that a superior healing strength was observed in the silk material of *N. pilipes*, which was in accordance with the morphological evidence (**Fig. 2B-D**).

Last, the silk film’s deformation angle (θ), the degree of bending of the healed film that corresponds to the tensile force application, was also plotted along the pushing distance exerted from the metal bar. As depicted in **Fig. 2F**, the θ of *N. pilipes* film increased monotonically with the progression of the bar. It was endurable toward the external force up to θ ∼20° (1.6 mm for the bar migration) before rupturing again at the healed region. Conversely, *C. moluccensis* film exhibited minimal deformation, θ∼ 2.5°, along with the bar migration. In fact, the repaired *C. moluccensis* film was unresisting to the tensile force exerted from the bar, leading to an immediate rupture of the film. Therefore, no tension was built up toward the film to cause further film deformation, yielding a low degree in **Fig. 2F**.

Overall, the fabricated silk materials from our selection possess self-repair capability, particularly the MA silk of *N. pilipes* in comparison to the other forms of silks examined in this study. Such an original discovery offers a novel route in comprehending the nature of silk materials and engineering bioinspired adaptive materials.

### Construction of biosynthetic spidroins of *N. pilipes*

Inspired by the remarkable material healing competence of MA silk from *N. pilipes*, we sought to generate bioengineered materials with recapitulated healable functionality. To achieve this, two strategic approaches, next-generation sequencing (NGS) and synthetic biology (SynBio), were employed to decode the intricate silk library of *N. pilipes* and also synthesize biomimetic silk spidroins respectively.

First, the MA silk glands of female *N. pilipes* were isolated, followed by mRNA extraction, transcriptomic NGS analysis, and spidroin assembly and construction (**Fig. 3A**). For the candidate screening, three selection criteria were considered: expression level, transcript length, and spidroin signature motifs (poly-alanine and poly-glycine tandem repeats). Briefly, 6 variants of MaSp1 associated transcripts and 7 variants of MaSp2 associated transcripts of spidroins were identified (**Supplementary Table_contig**) and summarized in **Supplementary Fig. S1A**.

**Figure 3.**
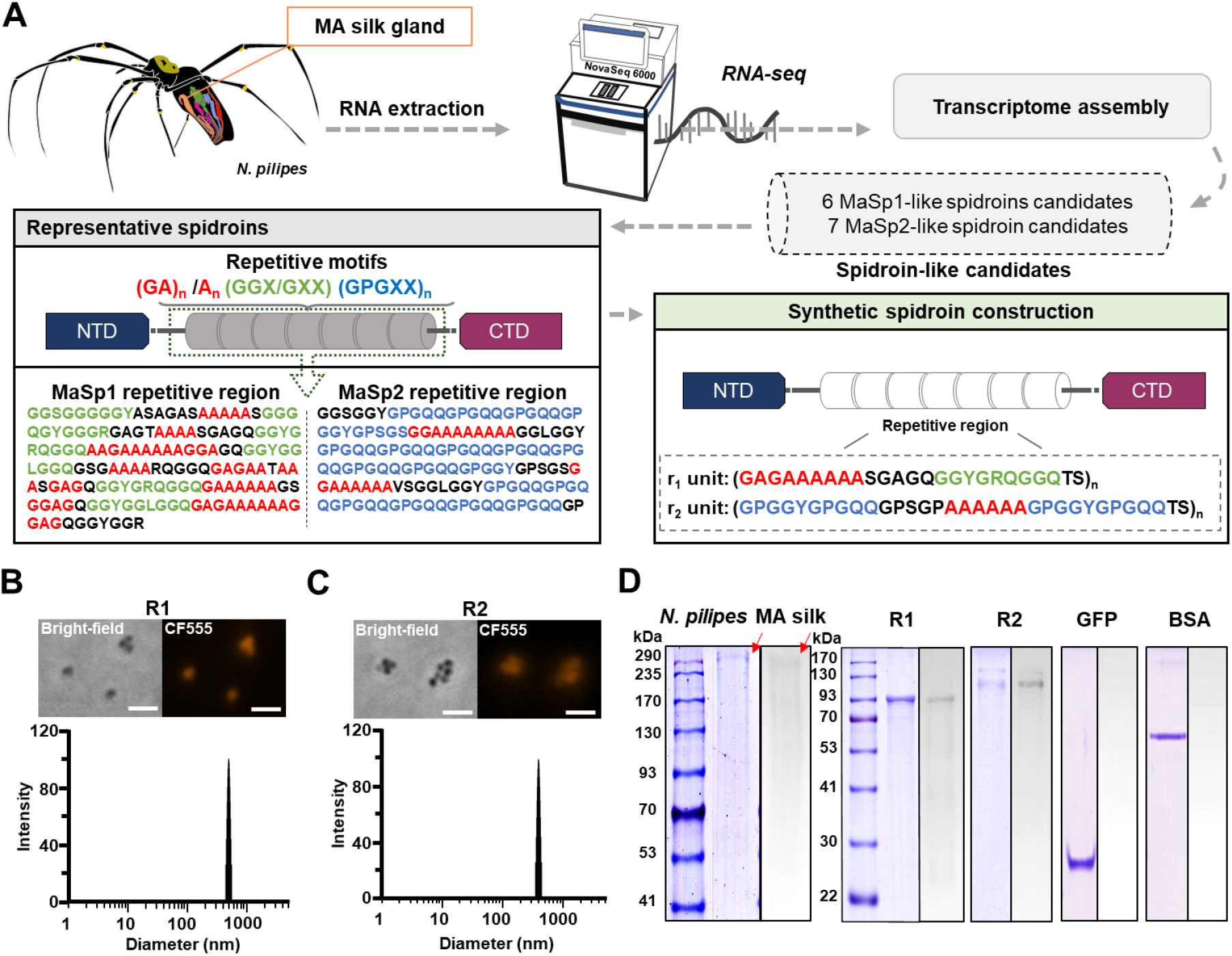
Construction of biosynthetic spidroins. (A) Transcriptomic study of MA glands from *N. pilipes*. The MA glands were isolated by *N. pilipes* dissection (top left) with RNA extraction and NGS analysis and assembly (top right Subsequently, the obtained candidate sequences were blasted for spidroin identification and annotation (bottom left) and the biosynthetic spidroins, R1 and R2, were designed and constructed (bottom right). (B) The nano-structural analyses of biosynthetic R1. The nanostructure was identified through immunofluorescence (top; scale bar: 2μm) and DLS (bottom). (C) The nano-structural analyses of biosynthetic R1. The nanostructure was identified through immunofluorescence (top; scale bar: 2μm) and DLS (bottom). (D) Protein analysis of R1 and R2. R1 and R2 were evaluated by SDS-PAGE and Western blot (via anti-NTD) for MW and purity identification. SDS-PAGE (left land) and Western blotting (right land) results were arranged for each sample. *N. pilipes* dragline silk was served as a positive control, and GFP & BSA as negative controls.

Subsequently, the consensus repetitive motifs within the transcript candidates were identified and shown in **Fig. 3A bottom left**. With the motif analysis, a series of iconic patterns were identified: (GA)_n_ /A_n_, (GGX/GXX), and (GPGXX)_n_. (GA)_n_ and A_n_, known as beta-sheet associated motifs, are the main building blocks for the crystallinity structures in both MaSp1 and MaSp2. (GGX/GXX), the helical motif, was only identified in MaSp1. (GPGXX)_n_, beta-spiral motif that is responsible for the elasticity of spider silk, is a signature in MaSp2. Representatively, the core repetitive regions of MaSp1 and MaSp2 were also shown (**Supplementary Fig. S1B**).

Building upon the sequence hierarchy of MaSp1 and MaSp2, two biomimetic motifs, namely r_1_ and r_2_, were designed accordingly. As indicated in **Fig. 3A bottom right**, the synthetic motifs of r_1_ and r_2_, monomeric building blocks, were repetitively elongated as the core of the silk polymers, flanked by the NTD and CTD domains (**Supplementary Fig. S1C**). With the BioBrick cloning strategies, we successfully constructed 32 duplications of r_1_ and r_2_ units, yielding synthetic silk constructs R1 (predicted MW: 91 kDa) and R2 (predicted MW:121 kDa). Further cloning strategies and plasmids were provided in **Supplementary Figs. S2 & S3**.

Consequently, both biosynthetic R1 and R2 spidroins were expressed in *E. coli* BLR(DE3) • Δ*endA* by fermentation and extracted using a novel purification methodology that is distinct from conventional affinity-based protein separation strategy. Inspired by the micelle-forming nature of native MA spidroins, we envisioned that the biosynthetic R1 and R2 could self-assemble into nanostructures, featuring a facile size-based purification strategy. Herein, we established a time- and cost-efficient purification protocol tailored for nanostructured spider silk materials by detergent washing and differential centrifugation. Our original spider silk purification scheme achieved the final spider silk products in a timely manner, without the extra procedures of chromatography and dialysis [59-61].

To characterize the nanostructures from the purified R1 and R2 samples, a primary antibody that targets the NTD domain, designated as mouse anti-NTD, was generated in this research to facilitate the silk evaluation **(Supplementary Fig. S4**). Both purified R1 and R2 samples were treated with mouse anti-NTD, followed by CF555 conjugated goat anti-mouse, and observed under a fluorescence microscope. As the results depicted in **Fig. 3B**, the purified R1 nanoparticles (∼400 nm) were observed in the bright-field images, in perfect accordance with the red signal by NTD immunostaining (red fluorescence image). Additionally, the DLS analysis of R1 products also revealed a monodispersed population (PDI ∼0.005) with a particle size ∼490 nm, which is remarkably consistent with the R1 imaging results. Similarly, R2 was confirmed by microscopy, immunofluorescence, and DLS analysis in **Fig. 3C**. The particle size of R2 by microscopic observation, ∼300nm, was comparable to the corresponding DLS analysis, at ∼362 nm (PDI ∼0.005).

In addition to probing the supra-nanostructures, SDS-PAGE and Western blotting were carried out to determine the molecular weight and purity of the purified silk products. Briefly, R1 and R2 were individually lyophilized, solubilized in HFIP, and examined by SDS-PAGE. Subsequently, Western blotting, transferred from the SDS-PAGE, was performed via the same immunostaining process (in **Figs. 3B & C**) with mouse anti-NTD. In **Fig. 3D**, natural MA silk from *Nephila pilipes* (positive control) revealed a significant signal at MW ∼300 kDa on both SDS-PAGE and Western blotting. Two negative controls, GFP (green fluorescence protein; predicted MW: 27 kDa) and BSA (bovine serum albumin; predicted MW: 60 kDa), showed corresponding protein bands in SDS-PAGE yet were absent in Western blots. With a high targeting specificity, anti-NTD was confirmed to be a reliable biomolecular probe for identifying NTD-carrying spidroins. Last, purified R1 and R2 were observed showing distinct signals at 90 kDa and 120 kDa respectively, in both SDS-PAGE and Western blots. Again, such results reinforce the validity of our novel spidroin purification process, yielding high purity of R1 and R2 at the biomolecular levels.

### Self-healing of biosynthetic spidroins

To profile the healability of the biosynthetic spidroins, R1 and R2, systematic material self-repairing examinations were performed. Briefly, the purified silk materials were lyophilized (**Fig. 4A top left**) and completely dissolved in HFIP, yielding a transparent silk solution (**Fig. 4A top right**). Subsequently, thin films of R1 and R2 were cast separately and exhibited significant transparency (**Fig. 4A bottom left**). R2 films were also further folded into an origami crane as a demonstration of their flexibility (**Fig. 4A bottom right**).

**Figure 4.**
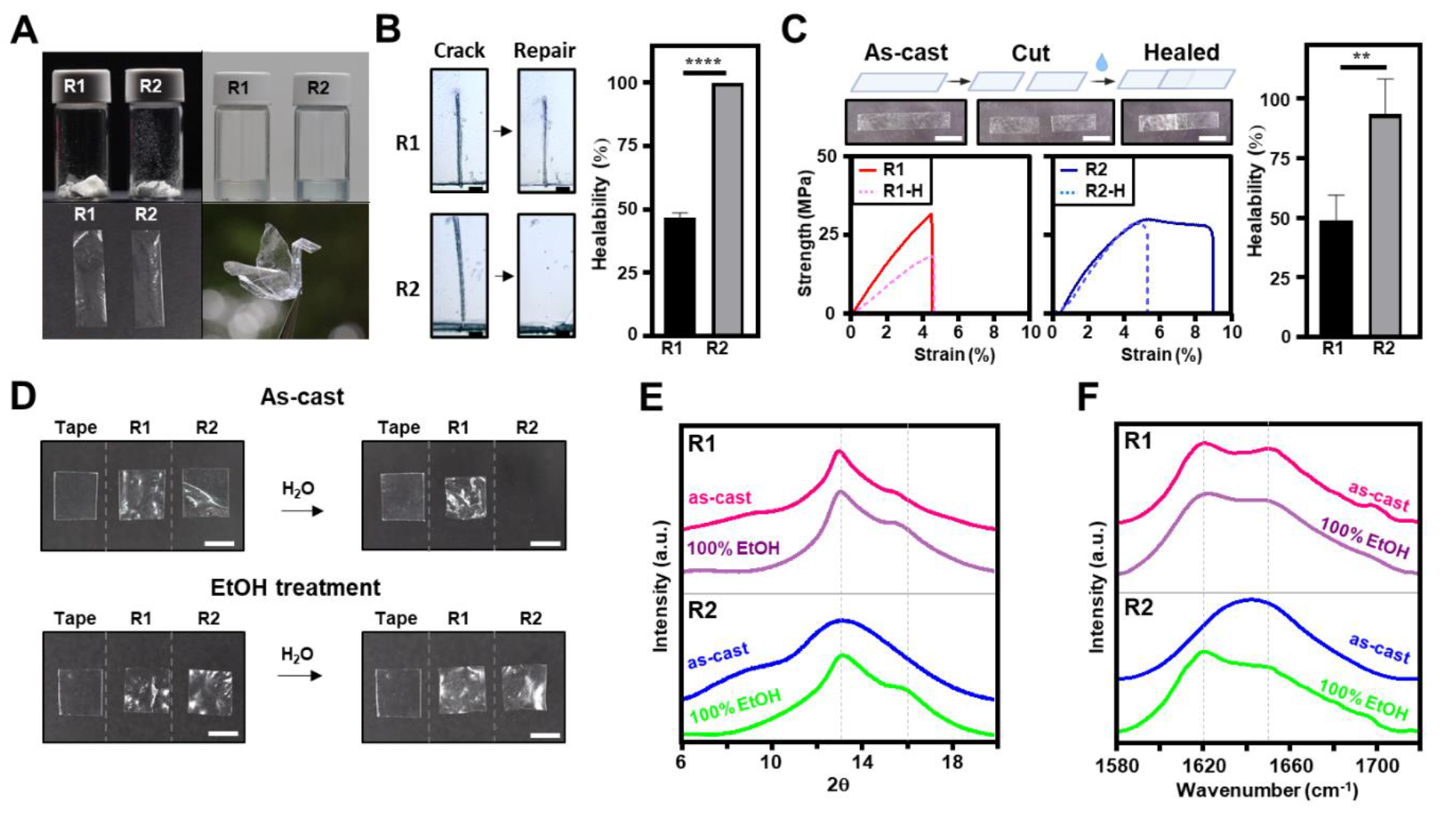
Characterization and self-healing of the biosynthetic spidroins films. (A) The purified biosynthetic spidroin powder, solution, and cast-films were presented, with the additional origami crane fabrication (by R2 films). (B) Crack-and-repair examination of biosynthetic spidroin films. Similar procedures to Fig. 2(B)-(D), the scar restoration efficiency for R1 and R2 was estimated (scale bar: 100μm) and the results were presented (n=5; **** p<0.0001). (C) Biosynthetic spidroin film tensile strength after self-healing. The tensile strength of the cut-and-healed silk films (designated R1-H & R2-H) was determined, in comparison to R1 & R2 as-cast (uncut) films. (n=4; **p<0.01) (D) Water stability of biosynthetic spidroin films. Both as-cast and ethanol-treated spidroin films were immersed in water, and the integrity of the resulting films was examined by imaging. (scale bar: 5mm) (E) The XRD patterns of biosynthetic R1 and R2 films before and after being treated with ethanol. (F) FTIR analysis of R1 and R2 films with ethanol treatment (control: as-cast samples).

Next, cast R1 and R2 thin films were challenged for the crack and repair test, similar to assays of **Fig. 2B**. The results in **Fig. 4B left** indicated that the scar of R1 films remained after healing; however, no scar was observed for the R2 films after healing. The statistical analysis revealed the superior healability of R2 (∼100%) than that of R1 (∼47%) (****p<0.0001) in **Fig. 4B right**.

Furthermore, we evaluated the silk film-to-film cohesiveness, the cut-and-healing process, by tensile testing, offering additional evidence of repairable capacities of biosynthetic spidroins. Briefly, a piece of cast film, either R1 or R2, was cut in half, moisturized by water, and bond together at the minimal overlapped region (**Fig. 4C top left**). Subsequently, the tensile performance of the mended films was evaluated as a measure of self-repairability. Illustrated in **Fig. 4C bottom left**, the cohered R1 film (R1-H) exhibited a lower strength (∼14 MPa) than the untreated R1 film at 29 MPa. In contrast, the cohered R2 film (R2-H) displayed the same degree of tensile strength (26 MPa) as the untreated R2 film. During tensile strength measurements, the cohered region of R1-H often detached easily, but R2-H remained firmly bond throughout the test. Significantly, R2 films possessed stronger cohesive healability than R1 films (**p<0.01), as evidenced by the normalized tensile strength in **Fig. 4C right**.

Additionally, it’s noteworthy that R2 and R1 exhibited distinctive material responses in bulk water settings. The as-cast R1 film was impervious to water and retained its integrity following water immersion. Nevertheless, the as-cast R2 film was highly vulnerable to water disintegration and completely irretrievable following a water bath (**Fig. 4D top)**. As the control in the test, a clear slice of plastic tape that remained stable after water incubation was presented. Another finding of R2-assisted film propulsion (**Supplementary Figs. S5 & S6**) also provided evidence for R2 disintegration upon contact with water. Subsequently, an ethanol post-treatment strategy was employed to enhanced the R2 films integrity against water[50]. As the results shown in **Fig. 4D bottom**, the treated R2 films (pretreatment with 100% ethanol for 1hr) became water-repellent throughout incubation (complementary to **Fig. 4D top**). Whereas, the ethanol-treated R1 films remained unaffected after the water incubation challenge.

In this context, we further accessed the structural evolution of both R1 and R2 in respond to ethanol treatments via XRD and FTIR. **Fig. 4E** illustrates silk film crystallinity peaks at 2θ of 13.5° and 16° corresponded to β-sheet nanostructures, as a result of synchrotron radiation XRD analysis. The XRD profile of R1 films were comparably unaltered before and after ethanol treatments, indicating that the highest degree of crystalline structures was already present in the as-cast R1 films (**Supplementary Fig. S7**). Yet, R2 films showed substantial enhancement in crystallinity after ethanol treatment.

In accordance with XRD findings, ATR-FTIR analyses revealed similar nano-structural tendencies in R1 and R2. Particularly in **Fig. 4F**, the amide I region (1600-1700 cm^-1^) that reveals the secondary structures of silk samples was analyzed. As indicated for R1, the major peak at 1622 cm^-1^ signifies the β−sheet structures (crystalline) while another peak at 1650 cm^-1^ represents non-crystalline random coil structures. R1’s FITR spectra was unaffected by ethanol treatment, but R2 underwent a substantial conformational shift toward β−sheet crystalline structure.

In sum, biosynthetic spidroin R2 had more flexible and adaptable material characteristics than R1, and more importantly, R2 was highly repairable. In fact, R1, rich in β-sheet crystallinity, was and more rigid and less cohesive than R2 (crystallinity data shown in **Supplementary Fig. S8**). We believe that is due to the distinctive molecular architecture and hierarchical structure of R2. Given the considerable promise in material healing and flexibility, R2 was chosen as the model throughout the rest of the investigation into healability and applications.

### Programed self-healing of biosynthetic spidroin R2

Next, we sought to further systematically enhance the durability of biosynthetic spidroin R2 and access its healing effectiveness alongside post-fabrication processes. A series of treatments, including ethanol immersion (40%, 70%, and 100%) and WVA (at 37°C or 60 °C), were employed to process the as-cast R2 films (**Fig. 5A top**).

**Figure 5.**
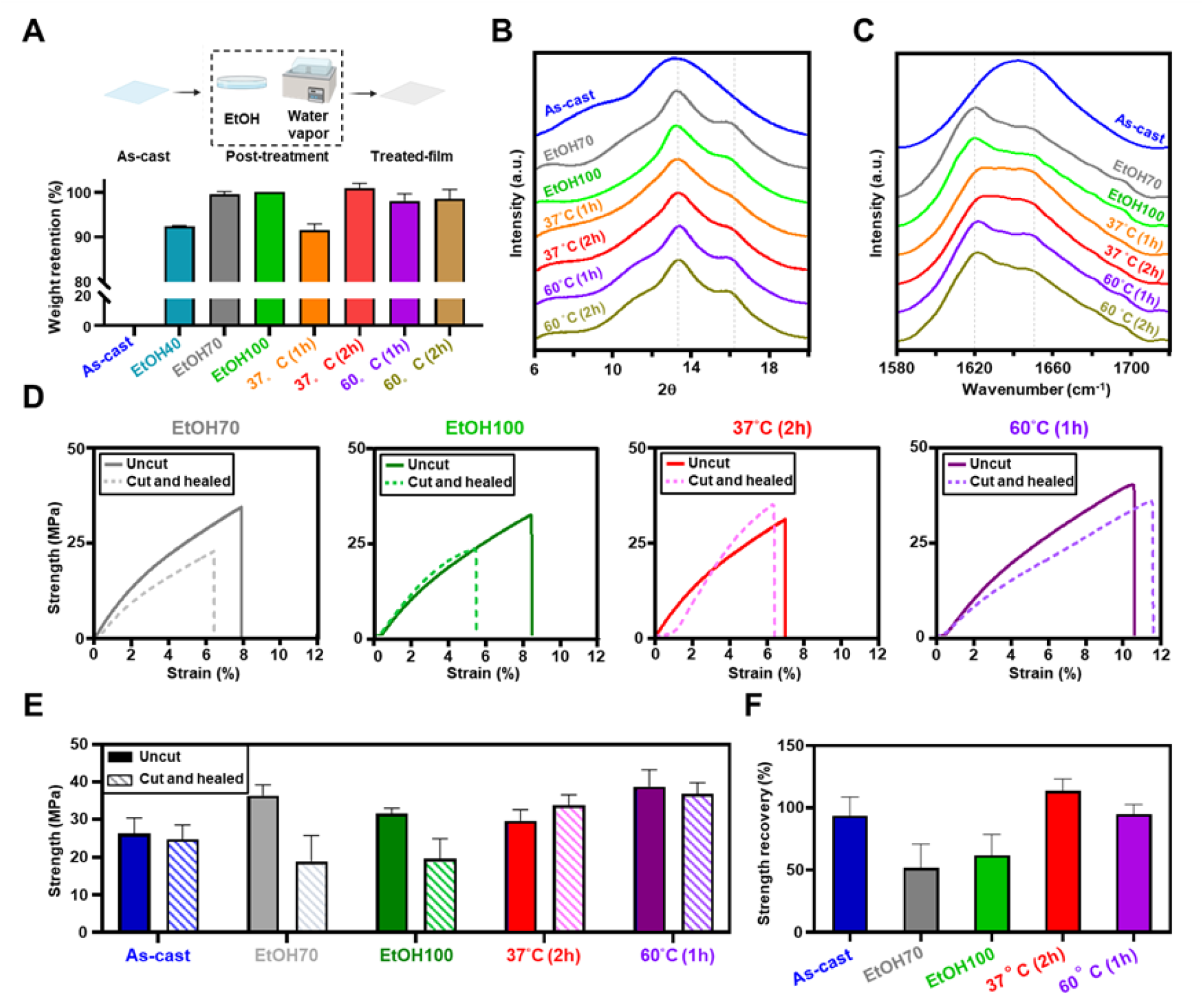
Post-treatment of biosynthetic spidroin films on structural stability and self-healability. (A) The water resistance of post-treated R2 films. A series of treatments, including ethanol immersion and WVA (water vapor annealing at 37°C or 60 °C), were introduced to R2 films. After post-treatments, the films were soaked in water to determine their weight retention ratio. ETOH40, ETOH70, ETOH100 denoted 40%, 70%, and 100% ethanol treatment, respectively. (B) XRD analysis of the post-treated R2 films. (C) The FTIR spectra of post-treated R2 films. (D) Tensile examinations of pre-treated R2 films to determine their capacity for self-healing. R2 films treated with ETOH70, ETOH100, WVA 37°C (2h), and WVA 60 °C (1h), were subjected to cut-and-healed procedures, and the tensile profiles were shown. Uncut: the control R2 films with the same treatment. The maximum strength of the tensile profiles among the treated R2 films was summarized in (E) and the normalized strength recovery efficiency of the cut-and-healed films was shown in (F).

Subsequently, the durability of treated R2 samples was probed by evaluating the weight retention of R2 films under a water-immersion challenge. In **Fig. 5A bottom**, the weight retention ratio of R2 films after water incubation was shown. Herein, 40%, 70%, and 100% ethanol treatments on R2 films were designated as EtOH40, EtOH70, and EtOH100 respectively. In WVA treatments, 37°C for 1hr, 37°C for 2hr, 60°C for 1hr, and 60°C for 2hr were denoted as 37°C (1h), 37°C (2h), 60°C (1h), and 60°C (2h), respectively. As illustrated in the plot, most of the treatments, EtOH70, EtOH100, 37°C (2h), 60°C (1h), and 60°C (2h) yielded strong weight retention of R2 (>98%). However, both EtOH40 and 37°C (1h) treatments yielded moderate weight loss (>8%), while the untreated R2 films rendered a complete weight loss, irretrievable after water incubation.

To follow up, we further tracked the nano-structural evolution of treated R2 using XRD and FTIR, in line with the water endurance tests abovementioned. As displayed in **Fig. 5B**, the progressively augmented peaks (13.5° and 16°) in XRD signified the advancement of enhanced crystallinity along the treatments, ranging from 37°C (1h), 37°C (2h), 60°C (1h), and 60°C (2h), EtOH70, and EtOH100. Showing a similar tendency in the ATR-FTIR analysis (**Fig. 5C**), we observed an elevated spike of β-sheet signature (1620 cm^-1^) over the random coil signal (1650 cm^-1^) of R2 along with the post treatments.

Additionally, both XRD and FTIR spectra of R2 under various treatments were subsequently deconvoluted by Gaussian decomposition for crystallinity estimation and secondary structural analysis, respectively. As summarized in **Supplementary Figs. S7 & S8**, our structural analyses reported a strong correlation between XRD and FTIR for R2 materials. For as-cast R2 sample (untreated), the deconvoluted results in XRD and FTIR both revealed the lowest and similar ratios, 19.9% in XRD crystallinity and 21.7% in β-sheet structures. The R2 sample under EtOH100 treatment yielded the most ordered structural ratios, 32.8% in XRD crystallinity and 35.3% in β-sheet structure. Besides, the sequence of 60°C WVA treatments resulted in stronger ordered structural ratios than 37°C WVA treatments.

After gaining knowledge in tuning R2’s material stability and structural organization, we proceeded to characterize the cohesive healability of treated R2. Explicitly, we chose EtOH70, EtOH100, 37°C (2h), and 60°C (1h) as our experimental sets, due to enhanced film durability against water. Subsequently, tensile tests were used to evaluate the repairability of the pretreated R2 films by gauging their degree of regained strength after being repaired (similar to **Fig. 4C**). In **Fig. 5D**, representative tensile strength profiles of each pretreated R2 are shown, and in **Fig. 5E**, the highest strengths are summarized and plotted. Interestingly, different post-treatments posed varying impacts on the tensile strength of uncut films. Compared to as-cast R2 (untreated), R2 films treated with ethanol and WVA were considerably strengthened. Furthermore, after being cut and healed, the restored strengths of the WVA-treated R2 films were equivalent to their counterparts (uncut samples), indicating a completely healable capacity. Yet in the ethanol treated groups, the cut-and-healed samples were unable to match the native strength of the uncut counterparts.

A further relative strength recovery among the treatments was calculated by normalizing the strength of the cut-and-healed samples to the strength of their corresponding uncut counterparts. This, in turn, could serves as a cohesiveness index to determine the repair efficiency, in terms of mechanical performance. Shown in **Fig. 5F**, both EtOH100 and EtOH70 treated R2 samples exhibited inferior strength recovery, whereas R2 samples treated with WVA at 37°C (2h) and 60°C (1h) restored full strength throughput the repairing process.

In light of the fact that WVA treatments (at 37°C and 60°C) produced R2 films with the most robust healability and stability, we also distinguished between these two procedures. During bulk water incubation, patched R2 films pre-treated at WVA 60°C separated, while patched R2 films pre-treated at WVA 37°C remained securely adhered (data not shown). The distinctions in the healing mechanisms of R2 materials, including either permanent or temporary bonding, may provide us with additional ideas for the development of diverse material repair structures.

### Self-healing mechanisms of biosynthetic spidroins

Learning from our biosynthetic spidroin models, we hypothesize that the secrets of the healing capacity lies within in the dynamic interplay between their noncrystalline regions. The amorphous components of biosynthetic silk materials are principally responsible for the adaptive healing properties. Silk materials are capable of fast repair owing to the supramolecular coupling of freely accessible chains driven by non-covalent hydrogen bonds. Additionally, the plasticizing impact of water was employed in biosynthetic silk materials to actively soften the freely unstructured polymeric chains upon physical contact, hence accelerating the cohesive bonding efficiency.

Additionally, this work has demonstrated, for the first time, the tunable healability of the biosynthetic silk materials. There distinct degrees of healing are accomplished by fabrication processes: irreversible, reversible, and minimal cohesiveness. In line with the hypothesized amorphousness-driven healing paradigm, the differential healing capacities of treated silk materials correspond to their amorphous composition. Milder treated biosynthetic spidroins with the highest number of mobile chains (37°C WVA) result in permanent cohesiveness upon healing, while overtreated spidroin samples with less mobile chains result in worse healing (ethanol). In the context of creating future repairable biomaterials or biointerfaces, programmable biosynthetic spidroin materials with durability and reversibility in cohesion (60°C WVA) might provide an intriguing viewpoint.

In contrast to our amorphous domain-driven healing hypothesis, the active participation of the crystalline fractions in self-healing of biosynthetic spidroins is yet to be validated further. Higashi et al. (2021) [62] has recently proposed a material healing model based on the β-sheet crystallinity of alanine-rich synthetic peptides. Conversely, Pena-Francesch and Demirel (2020) [30] on the bioengineered SRT model, primarily substantiated the key function of amorphous regions in SRT material damage repairing. In fact, protein-based intrinsic self-healing mechanisms are less extensively explored than those based on chemically-synthesized materials. The intricate healing dynamics of protein-based materials would be considerably influenced by a large number of connected aspects, such as biochemical designs, molecular weights, and manufacturing processes. Further research efforts are essential to elucidate the underlying molecular mechanisms and develop future healable materials.

### Biosynthetic R2-based flexible logic gate devices

In addition to the adaptive cohesive performance, we sought to endow the biosynthetic spidroin with additional functionalities, such as electrical competence. Specifically, graphene, an inorganic engineering material extensively utilized in electronics and medical applications, was blended with the biosynthetic spidroin to form a novel hybrid composite[57, 63]. As a proof-of-concept demonstration, the potentials of fabricating electronic biodevices using the healable spider silk materials were explored further. To achieve such goal, we investigated the effect of graphene doping on the overall cohesiveness and mechanical performance of the hybrid biosynthetic spidroin composites.

In our present setup, hybrid graphene-to-R2 composites (R2G) were produced in the form of cast films with a doping ratio of 0.15. Shown in **Supplementary Fig. S9**, R2G displayed strong mechanical performance and processability similar to R2 (p>0.05). Next, R2G performed comparable self-healing capacity, in comparison to its counterpart. These results, in turn, demonstrate that the intrinsic R2 properties were significantly preserved when graphene was incorporated.

Furthermore, the graphene-associated electrical conductivity of R2G was also evaluated. First, R2G strips were fabricated and pretreated at 37°C WVA (2h), followed by the cut-and-healed treatments (**Fig. 6A**). Subsequently, the repaired R2G samples were also placed into a simple electrical circuit, with the intention of evaluating the conductivity of both uncut R2G and the mended R2G. In **Fig. 6B**, the basic design featuring a DC battery, a blue LED, and a toggle switch, was constructed, and R2G (black strip) was connected to the circuit through two arrow electrodes. The completion of the circuity, signaled by the LED light output, is highly dependent on the conductivity of R2G, as the conductive wire. As a test result depicted in **Fig. 6B bottom**, the blue light successfully illuminated with the connection of uncut R2G, indicating that the graphene in R2G was electrically active. Furthermore, the healed R2G displayed retrieved conductivity by consistently lighting up the LED, but the unrepaired control unit (two physically overlapping portions of sliced R2G without healing) failed to restore conductivity.

**Figure 6.**
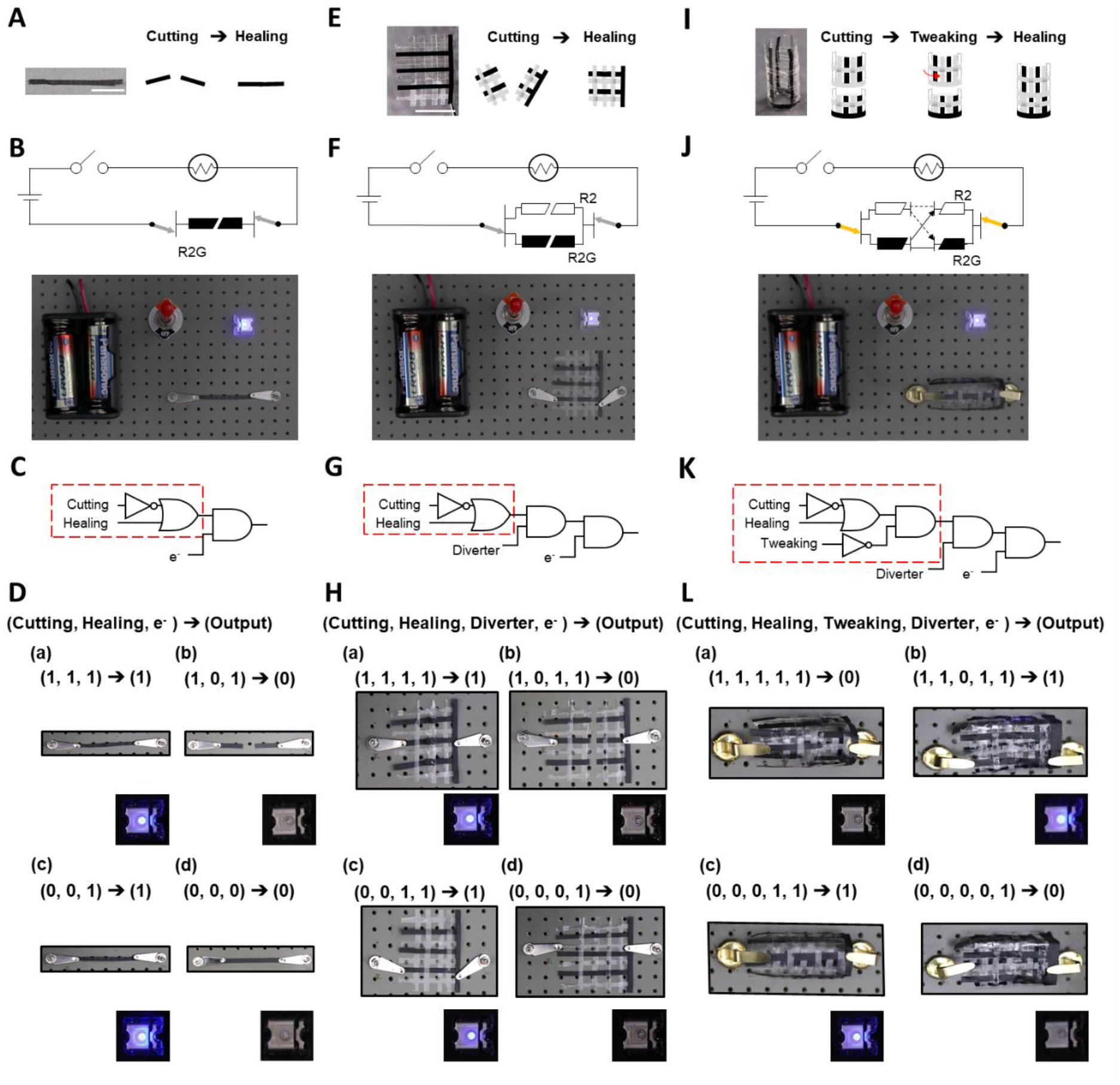
Biosynthetic R2 spidroin-based flexible logic gate circuits. (A) Graphene-doped R2 (R2G) and 1D fabric. The R2G pretreated with WVA 37°C (2h) was self-healable via the cut-and-healed process, generating the 1D patched fabric. (B) 1D fiber integration into a simple circuit device. The design of the open circuit included a battery, a switch, and a LED light. With the integration of 1D-R2G strip as the conducting wire to complete the circuit, the LED light was activated. (C) The 1D-R2G circuit’s logic gate construction using Boolean operators. The red-dashed square symbolized the processing of the 1D-R2G. “NOT (no cutting)” & “OR (with healing)”, are the major operators to achieve the success of LED lighting, as output “1”. (D) The outcomes of fabricated R2G in the simple circuit device and corresponding logic gate operation. (E) Self-healable woven 2D-R2/R2G fabric. (F) 2D R2/R2G fabric in the simple circuit. (G) 2D-fabric associated logic gate construction. (H) The experimental results of the 2D fabric logic gate circuit device. (I) 3D R2/R2G fabric construction. The “tweaking” can cause the misalignment of R2G/R2 in the structure. (J) & (K) indicated the integration of 3D fabric into the circuit and relevant logic gate design. “NOT (no tweaking)” is also essential for generating positive lighting output “1”. (L) The experimental results of 3D fabric in the simple circuit device. See **Supplementary Figs. 10-12** for detailed results.

Next, we used a Boolean operation strategy to build up a working model that describes the LED signaling output as a function of R2G physical layouts in the circuity. In other words, our goal throughout the materials processing and circuit design was to convert the R2G-encoded information into digital signals in the form of logic gate functions. As illustrated in **Fig. 6C**, this logic gate layout was designed with three levels of regulatory inputs (R2G cutting, R2G healing, and electricity), and one final output (LED lighting). The red-dashed square represented the combination of the following Boolean operations on R2G strips processing: “NOT (no R2G cutting)” and “OR (R2G healing)”.

Accordingly, a resultant truth table based on the Boolean setup was summarized in **Supplementary Table S1,** where a complete series of R2G fabrication and circuit assembly experiments were implemented and validated. Additionally, several representative results noted as cases **(a) ∼ (d)** were shown in **Fig. 6D**. In each case, we particularly highlighted the connection between the R2G input and LED output, as seen by the enlarged photos of the tests. As comprehended, the status of R2G inputs determined the final output, “1” for LED on while “0” for LED off. The remaining findings for truth table validation were reported in **Supplementary Fig. S10**, and those all matched the predicted outcomes in the truth table. Overall, the linear form of R2G strips (1D) exhibited both healability and electrical conductivity demonstrated in the simple logic gate test.

Building upon the initial success abovementioned, more sophisticated structural logic gate configurations, using 2D- or 3D- R2G fabrics, with repairability and moldability would be explored further. First, the 2D model were fabricated and demonstrated in **Fig. 6E-H**. As indicated in **Fig. 6E left**, R2 (transparent strips) and R2G (black strips) were manually woven into an interlaced 2D fabric structure as a primitive form of biosynthetic silk cloth, which was then treated with WVA at 37°C (2h). After drying, the inter-junctions of R2 and R2G were firmly bonded, resulting in a stable yet flexible spider silk-based braided fabric. The prototype fabric was implemented into the same electrical circuit design (**Fig. 6F**). Furthermore, post-processed fabric samples (with cut-and-healed treatments; see **Fig. 6E right**) were also put in the circuit individually for assessment.

**Fig. 6F** depicted a Boolean logic gate design and an example of the actual implementation utilizing 2D- R2/R2G fabric. In contrast to the 1D- R2G logic gate test (**Fig. 6A-D**), the 2D- R2/R2G configuration had an extra Boolean operation unit called diverter (**Fig. 6F-H**). As a spatially signal-diverting procedure in the circuit, connecting electrodes to the R2G (with electrical conductivity information) or R2 (without electrical conductivity information) inside the 2D fabric also determined circuit completion. In turn, the 2D truth table was shown in **Supplementary Table S2.** The overall signaling outcome was summarized in **Supplementary Fig. S11**, and several selected experimental results, cases (a) to (d), were demonstrated in **Fig. 6H**. In case (a), despite the fact that the 2D fabric was pre-treated under the cut-and-healed process, the final LED output remained positive “1”. In case (d), for instance, the LED light was switched off “0” because the “AND” operation of the diverter was set to “0”, with the electrode was directed to R2 (without electrical conductivity information).

Last, the 3D fabric-based logic gate model (**Fig. 6I-L**) was further developed and demonstrated by utilizing the flexibility and cohesiveness of the biosynthetic spider silk. The 2D fabric (identical to **Fig. 6E**) was bent and bonded with the application of water, resulting in the 3D tubular fabric structure seen in **Fig. 6I left**. Following that, the 3D tubular fabric structures were also subjected to the cutting-and-healing treatments, and we further employed a “Tweaking” operation to alter the order of alignment in rejoining both pieces of severed tubular fabric. As a result, two variant forms of patched products appeared, the original form (R2-to-R2 & R2G-to-R2G) and the tweaked form (R2-to-R2G or R2G-to-R2), as schemed in **Fig. 6I**.

Furthermore, the information content from the physical 3D fabric circuit (**Fig. 6J**) was extracted and transformed into the conceptual logic gate design in **Fig. 6K**. The integration of Boolean operations in the 3D logic-gate fabric layout, including cutting, healing, tweaking, and diverting, offered a dynamic operational layer in signal processing. The truth table was presented in **Supplementary Table S3.** Several exemplary outcomes, **cases (a)-(d)**, in **Fig. 6L**, highlighted the success of our 3D fabric design. For case (a), it was notable that the tweaked fabric patch resulted in an incomplete circuit (nonconductive R2 bonded to conductive R2G), resulting in no LED light as the final output. Several more experimental validations were carried out and summarized in **Supplementary Fig. S12**). Overall, R2 and R2G films treated with 37℃ WVA were woven in various intertwined architectures (1D to 3D) as a demonstration of flexible and repairable biosynthetic spidroin-based electrical logic gates application.

### Biosynthetic R2 based wearable device

Finally, we sought to explore the feasibility of reversible healing in biosynthetic spidroins, as well as its utility in bio-device applications. Encouraged by the preliminary results of transient cohesion of pretreated R2 at 60°C WVA, we emphasized further determining the cyclic healing-releasing profile of R2 films. Herein, a variety of R2 films prepared with 60°C (1 h) WVA were repeatedly self-healed and separated at the same cohesion location for 1 to 4 cycles, and the resultant films were mechanically tested. As seen in **Figs. 7A & B**, the films that went through multiple-healing cycles (1^st^-4^th^) had mechanical performance profiles that were equivalent to the control films (uncut), with no significant difference (p>0.05). As a result, 60°C (1 h) WVA-treated R2 exhibited reversible-healing capacities without sacrificing its mechanical performance.

**Figure 7.**
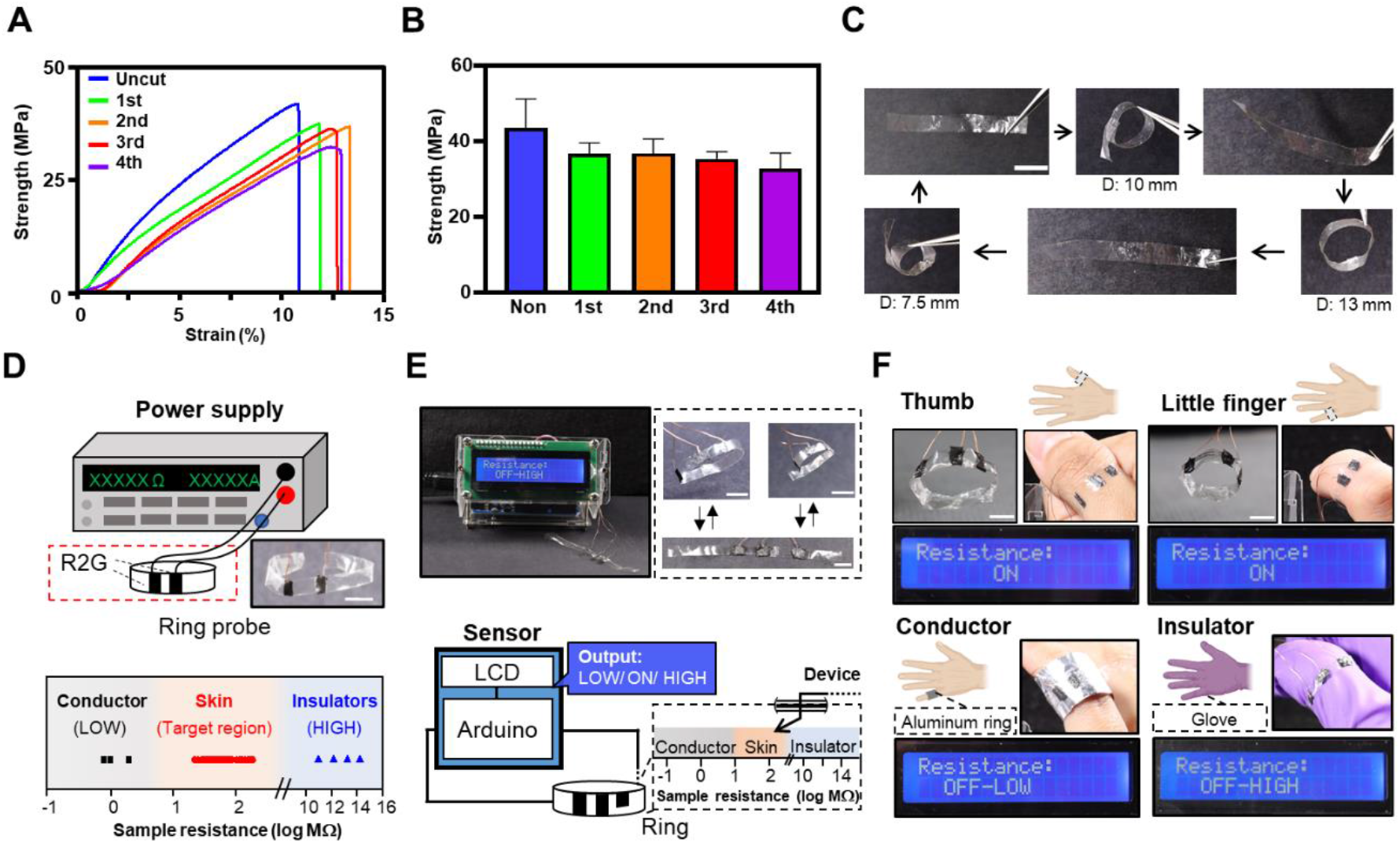
The re-healable R2 ring probe as wearable sensors. (A) The tensile performance of R2 sheets under reversible-healing conditions The R2 pretreated with WVA 60 °C (1h), was repeatedly self-healed and separated at the same cohesion location. The tensile performance of the R2 films were shown. (B) The maximum tensile strength of re-healable R2 film was summarized and plotted. No significant reduction in tensile strength was seen across cycling groups (p>0.05, ANOVA). (C) A reconfigurable R2 ring strip with variable sizes (7.5-13 mm in diameter) was accomplished by the repeatedly healing and re-opening of R2 strips. (D) R2G/R2 strip for fingertip resistance measurement. A complete resistance database was created using the R2G/R2 strip as the power source meter’s senor probe (bottom). (E) The portable and wearable finger recognition device was further created, including there-configurable ring probe (top right), the Arduino processing unit, and LCD panel for display (left). (F) The evaluation of the finger recognition device. The ring was adjustable for thumb (top left) or little finger (top right), both displaying the positive readout “ON” in the appropriate fingertip resistance range. The foil-wrapped finger was detected as “OFF-LOW” at the conductor range (bottom left); the glove-wearing finger was identified as an insulator with the signal readout “OFF-HIGH”.

Following that, the flexible and re-healable R2 was programmed into a tunable shape and structure. In **Fig. 7C**, the same piece of R2 film strip was able to repeatedly heal and re-open in cycles, generating customized circular rings with diameters of 10 mm, 13 mm, and 7.5 mm respectively.

Afterward, the reconfigurable R2 rings were further upgraded and used as finger ring probes for resistance measurement, or human skin sensors. Specifically, 3 small pieces of conductive R2G were integrated into the R2 ring via water-assisted adhesion, along with the embedment of enameled wires (**Fig. 7D top**). In this configuration, R2G patches were securely bonded to the inner surface of R2 rings, functioning as measuring electrodes. Furthermore, the enameled wires attached to the corresponding R2G electrodes were securely fixed between the R2G/R2 sandwiched layers before being connected to a power source meter. Consequently, the prototype ring sensor was utilized to measure the electrical resistance of physical objects.

To characterize the probe system, a collection of known conductive substances, including iron, copper, and aluminum, was used for baseline resistance measurement. In **Fig. 7D bottom**, the resultant reading of the conductors, as a reference control, was 1-2 M**Ω**. Apparently, such observed resistance level,1-2 M**Ω**, for the conductor materials was attributable to the intrinsic resistance of the R2G ring probe. Following that, a range of recognized non-conductive materials, including acrylic, plastic, and rubber, were used for resistance characterization of the non-conductive matters. After carefully defining the detectable ranges of our prototype system, we proceeded to take measurements of human subjects. As shown in **Fig. 7D bottom**, target resistance values between 20 and 200 MΩ were identified from our data source (7 people, over 70 fingertip measurements, spanning from thumbs to little fingers). All of the comprehensive data and measurements were provided in **Supplementary Tables S4-S6.** After confirming the utility of the resistance probe, a further fabrication of a portable and wearable finger recognition device was carried out. In brief, the data from the resistance library seen in **Fig. 7D** was used to program the Arduino unit, which acted as the signal processing and decision-making unit (see **Supplementary Fig. S14**). When the received signal from the silk ring probe was delivered for recognition to the Arduino interface, the decision result was then displayed on the LCD attached to the Arduino processor (see **Fig. 7E** top and **Supplementary Fig. S15**). In parallel with the Arduino setup (**Fig. 7E bottom left**), the silk-ring probe format was further refined and inspected in the studies. To boost the detection quality, one extra tiny piece of R2G was introduced in the ring probe as a ground electrode, in addition to the two original R2G patches as measuring electrodes (**Fig. 7E top & bottom left**). As a consequence, the fabricated wearable device with a re-configurable sensor probe ring was able to recognize the resistance of human fingertips.

In the final challenge of the wearable smart resistance device, an entirely new induvial (different from the participants who contributed to the resistance data library collection in **Supplementary Table S4**) was solicited. First, the single sensor probe was adjusted to fit either the test individual’s thumb or little finger, 13mm and 8mm, respectively as shown in **Fig. 7F top left & right**. As a result, both readouts displayed on the LCDs switched “ON”, indicating positive outcomes. When the test finger was wrapped in aluminum foil for detection, the signal readout shifted to “OFF-LOW” in **Fig. 7F bottom left**, indicating low electrical resistance. Also, in **Fig. 7F bottom right**, when the test finger was covered in a nitrile glove, the readout flipped to “OFF-HIGH,” suggesting an extra-high resistance. In light of such promising results, the silk-ring probe device might be deployed as part of a wearable biosensor or bioelectronic system.

## Conclusion

Spider silk is distinguished by its unique molecular hierarchy, with tandem repetitions of crystalline and amorphous domains, leading to intricate supramolecular interactions and remarkable physical characteristics. Many favorable capabilities of spider silk materials, like dynamic assemblability, pliability, and disturbance tolerance, can be further tweaked for self-adaptive material development. In this study, we revealed, for the first time, that spider dragline silk materials from *Nephila pilipes* exhibited remarkable intrinsic self-healing competence, as evidenced using thin film models for scratch repairing and cut healing. During a series of healing evaluations on two biosynthetic spidroins derived from *Nephila pilipes*, we identified R2 was substantially more prominent in healing capabilities in comparison to R1. Although further mechanistic research is needed, our pioneering work here has demonstrated spider dragline silk materials can be transformed into healable biomaterials and structures.

Next, the biosynthetic healable spidroin R2 was further integrated into prototype bioelectronics setups. An electrically conductive R2G with retained intrinsic cohesive characteristics and mechanical performance was successfully generated. Using woven R2G/R2 textiles as Boolean operators, unique logic gate circuits were created for the first demonstration. Flexible and repairable logic gates were constructed and tested over a variety of spatial hierarchies, from 1D to 3D, enabling the translation of digital electronic information through cascaded orthogonal logic circuits. Additionally, we also developed a wearable resistance probe in the form of a reconfigurable finger strip with R2G/R2. The re-shapable finger strip probe was reliably adjusted to fit user’s fingers of varying sizes. Such adaptable silk ring probe served as a conformable biometric recognition device.

Our proposed bioinspired self-healing spider silk model has the potential to readily expand versatility of silk-based materials. The bioengineered silk materials can be integrated into more sophisticated structures which can offer a spectrum of applications in wearable electronics, smart textiles, functional coatings, repairable devices, soft robotics, and protective gears. Indeed, there is a huge variety of untapped bio-resources that have intriguing chemical compositions and physical characteristics. By leveraging the power of synthetic biology and biofabrication, we can rapidly achieve biomaterials innovation and further transform the research landscape of biomimicry.

## Supporting Information

Supporting information is available from the Wiley Online Library.

## Author Contributions

W.-C.C., R.-C.W., S.-K.Y., T.-I.Y. and H.-C.W. were involved with conception and design of the work. W.-C.C., R.-C.W., J.-L.C. and Y.-H.K. were involved with strain cloning, fermentation, and purification. W.-C.C. and T.-Y.W. were involved with NGS experiments. S.-K.Y., P.-Y.C., H.-S.S. and S.-C.C. were involved with structural analyses. W.-C.C., S.-K.Y., W.-R.L. were involved with conductive silk experiments. W.-C.C, T.-I.Y. and H.-C.W. were involved with writing of the manuscript.

## Ethics approval

The experimental protocols for human subjects were reviewed and approved by Research Ethics Committee, National Taiwan University (NTU-REC No.: 202301HM017).

## Data Availability Statement

The data in this study are available from the corresponding author upon reasonable request.

## Acknowledgements

We acknowledge the support from National Taiwan University. We also thank R&D Center for Membrane Technology, Chung Yuan Christian University for technical assistance. This research is funded by National Synchrotron Radiation Research Center of Taiwan (2023-1-186-1), National Science and Technology Council of Taiwan (MOST 110-2224-E-038-001-; MOST 111-2221-E-033-003-; MOST 110-2314-B-002 -041 -MY3), Asian Office of Aerospace Research and Development (FA2386-20-1-4079; FA2386-22-1-4031), ITC-IPAC/DEVCOM (FA520922P0179).

## Supporting Information for Self-Healable Spider Dragline Silk Materials

### I. Identification of MA silk sequence from N. *pilipes*

Spider silk is one of the wonder materials in nature. MA spider silk, in particular, is a well-known high-performance biomaterial. To synthesize the biomimetic MA silk in our study, the first step is to identify the sequence of the native MA silk. The MA silk is composed of three distinct domains: N-terminal domain (NTD), C-terminal domain (CTD) and repetitive region. NTD and CTD are generally conserved among spider silk proteins, but repetitive regions typically contain distinct repetitive motifs in MaSp1 and MaSp2, two typical kinds of MA silk. MaSp1 silk protein contains poly-alanine and glycine-rich region whereas MaSp2 silk protein carries poly-alanine and GPGXX [1][2].

Here, we used two methodologies to identify the MA silk sequence from *N. pilipes*. First, the target fragments were amplified using polymerase chain reaction (PCR) and sequenced using Sanger sequencing. Afterwards, we used Illumina RNA-seq PE-150 reads (Genomics, Taiwan) to assemble the cDNA library from the MA silk gland.

The contigs sorting method is also described in Materials and Methods section in the main article. Briefly, the contigs were blasted against 594 Araneoidea spidroin related gene candidates and 134 *Nephila clavipes* spidroin proteins for open reading frames (ORFs) annotation. Next, the selected contigs were sorted according to the amino acid motif to determine the MA silk type (**Supplementary Fig. S1A**). The constructed cDNA library was uploaded to PubMed (Accession number: SRR21615954). The translation results of silk domains from Illumina sequencing and Sanger sequencing were shown in **Supplementary Fig. S1B-C**. Two representative transcripts that resemble the mostly typical MA silks, MaSp1 and MASp2, were identified as TRINITY_DN141_c0_g1_i17 and TRINITY_DN488_c0_g1_i9 respectively (**Supplementary Fig. S1B**). NTD (TRINITY_DN141_c0_g1_i17) and CTD (TRINITY_DN215_c0_g2_i1) sequences were isolated, identified, and confirmed accordingly in **Supplementary Fig. S1C**. Information was also described in the main article (Fig. 3A).

**Supplementary Figure S1.**
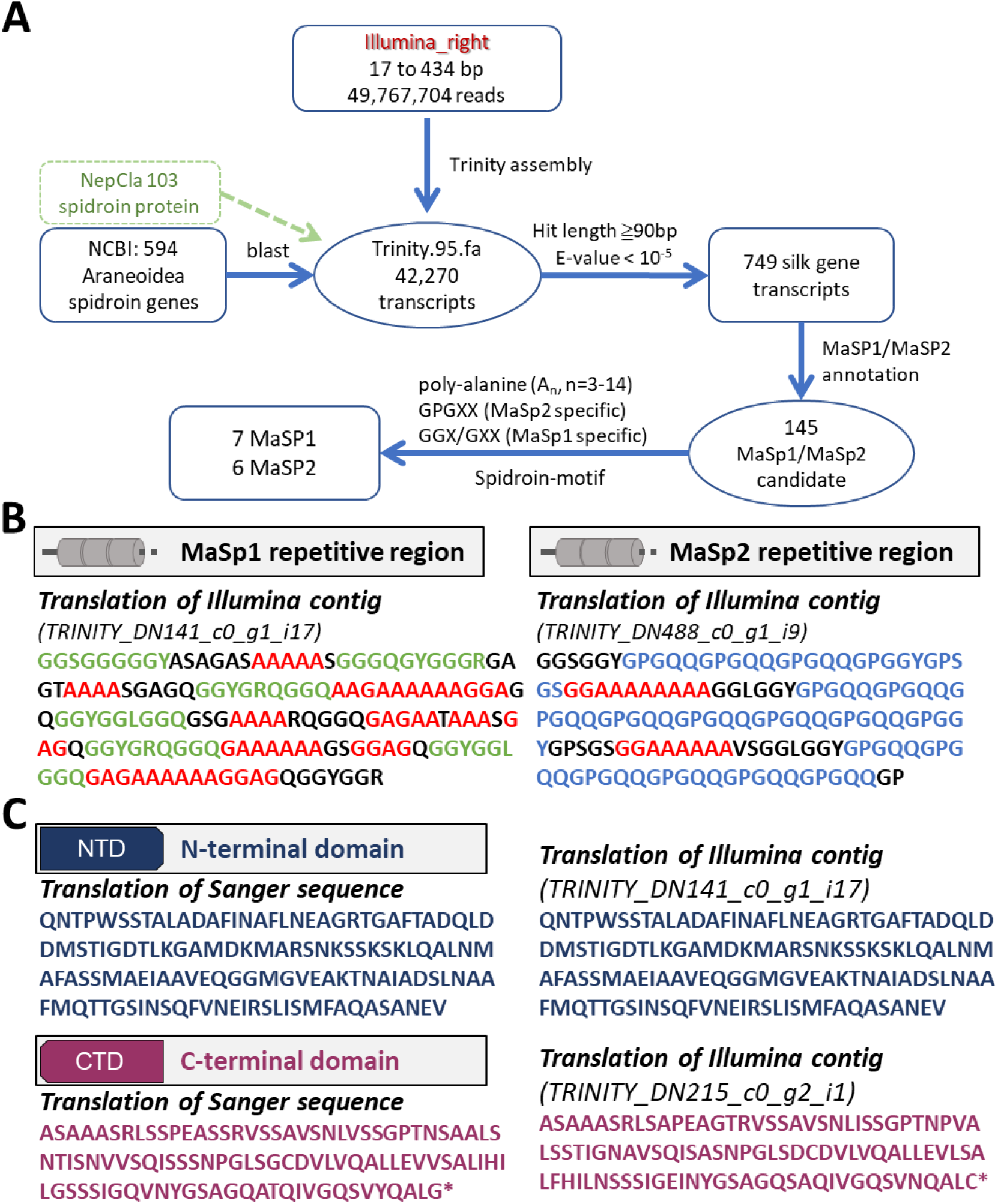
Sequence analysis of MA spider silk. (A) The flow chart of candidates sorting. (B) Partial sequencing results of MaSp1 and MaSp2 repetitive regions. Structural motifs were characterized as poly (A/GA) (red), GGX/GXX (Green), and GPGXX (Blue). (C) Sanger sequence and Illumina sequence results of NTD (dark blue) and CTD (dark red).

### II. Creation of codon-optimized biosynthetic MA silk gene for *E. coli* production

In order to construct the biosynthetic MA silk protein of our interest, the gene cassettes of NTD, CTD, MaSp1 repetitive unit (denoted as r1) and MaSp2 repetitive unit (denoted as r2) were developed using PCR. Each cassettes element was represented in Supplementary Fig. S2A.

Terminal domain cassettes, NTD and CTD, were amplified using overhang PCR from MA silk gland cDNA library. The r1 and r2 cassettes were identified using Illumina sequencing and reverse-translated into *E. coli* expression-optimized codons. The optimized r1 and r2 cassettes were created through overlap PCR. The gene cassettes were cloned onto the vector pET28a, individually.

Next, the gene cassettes were assembled followed by the outline indicated in Supplementary **Fig. S2B**. The first plasmid was cut using SpeI and HindIII, and the second was cut with NheI and HindIII. While HindIII tail were recombined to provide HindIII cutting sites again, the connection between SpeI and NheI was unable to be detected by restriction enzyme again[3]. Therefore, repetitive region was amplified with the cutting site flanked from previous fragment again. Following the procedure, plasmids pET28a-NTD-(r1)_32_-CTD and pET28a-NTD-(r2)_32_-CTD were generated, and the amino acid sequence of repetitive region were presented in **Supplementary Fig. S2C**.

For the protein expression, R1 spidroin was produced by pET28a-NTD-(r1)_32_-CTD in *E. coli* BLR(DE3) Δ*endA* (plasmid map in Supplementary Fig. S3B). For R2 expression, first, NTD-(r2)_32_-CTD were cloned into pENTH vector, pET28a without thrombin and His-tag expression to avoid unpredicted folding. Second, the expression of glycine tRNA and alanine tRNA was incorporated for enhanced the silk protein productivity[4]. Specifically, pSCLPP-glyT-alaT was introduced to enhance spidroin expression (plasmid map in Supplementary Fig. S3C). After selection, two strains were used to produce biosynthetic spidroins in 5L fermenters for further usage.

**Supplementary Figure S2.**
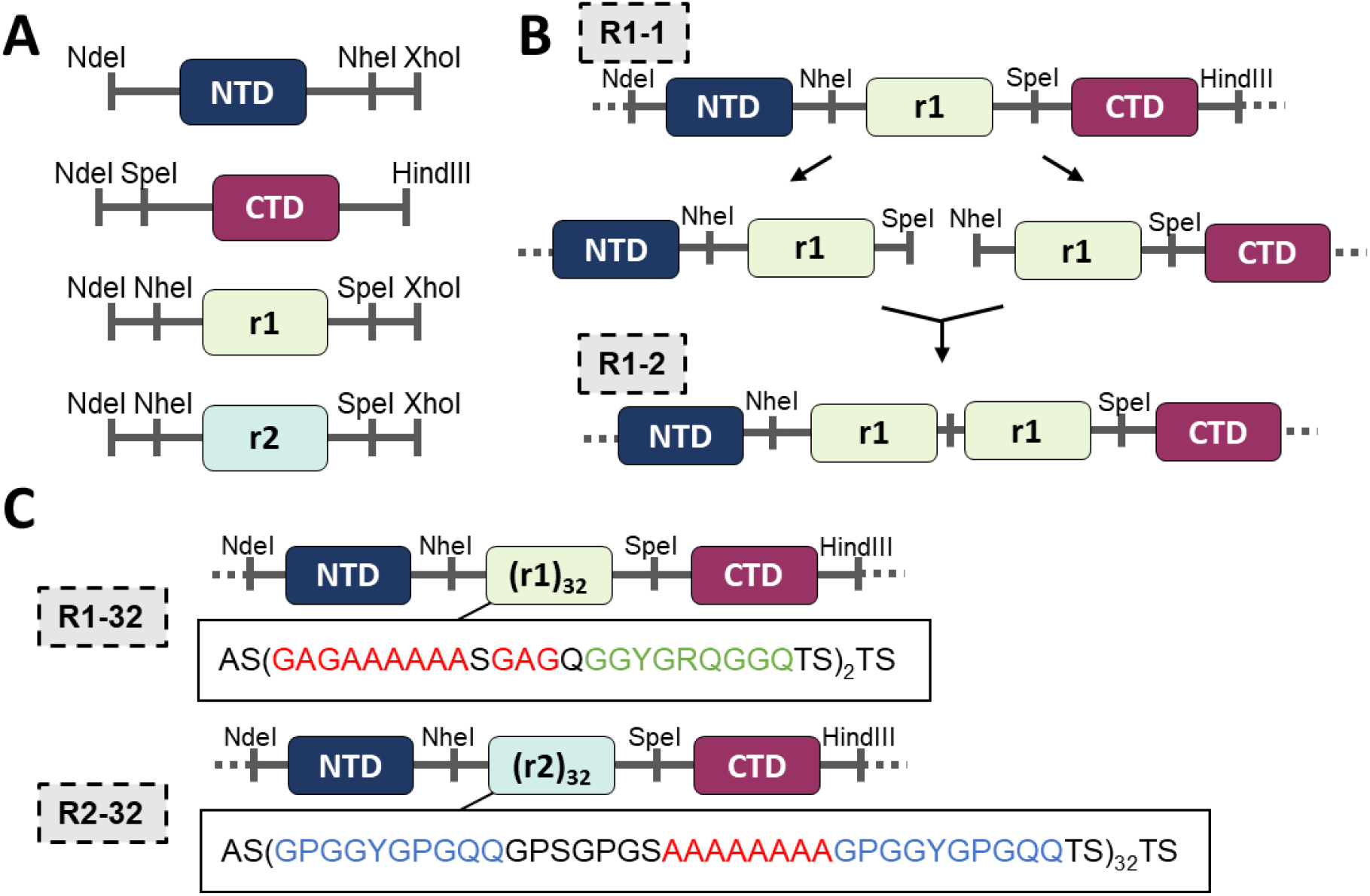
Creation of codon-optimized biosynthetic spidroins. (A) The terminal domain and repetitive unit cassettes construction by PCR amplification. (B) Amplification of repetitive units by BioBirck restriction enzyme cloning (SpeI and NheI pairs) (C) After cloning amplification, pET28a-NTD-(r1)32-CTD and pET28a-NTD-(r2)32-CTD were generated. The translation of repetitive regions was displayed and the motifs were characterized by poly (A/GA) (red), GGX/GXX (Green), and GPGXX (Blue).

**Supplementary Figure S3.**
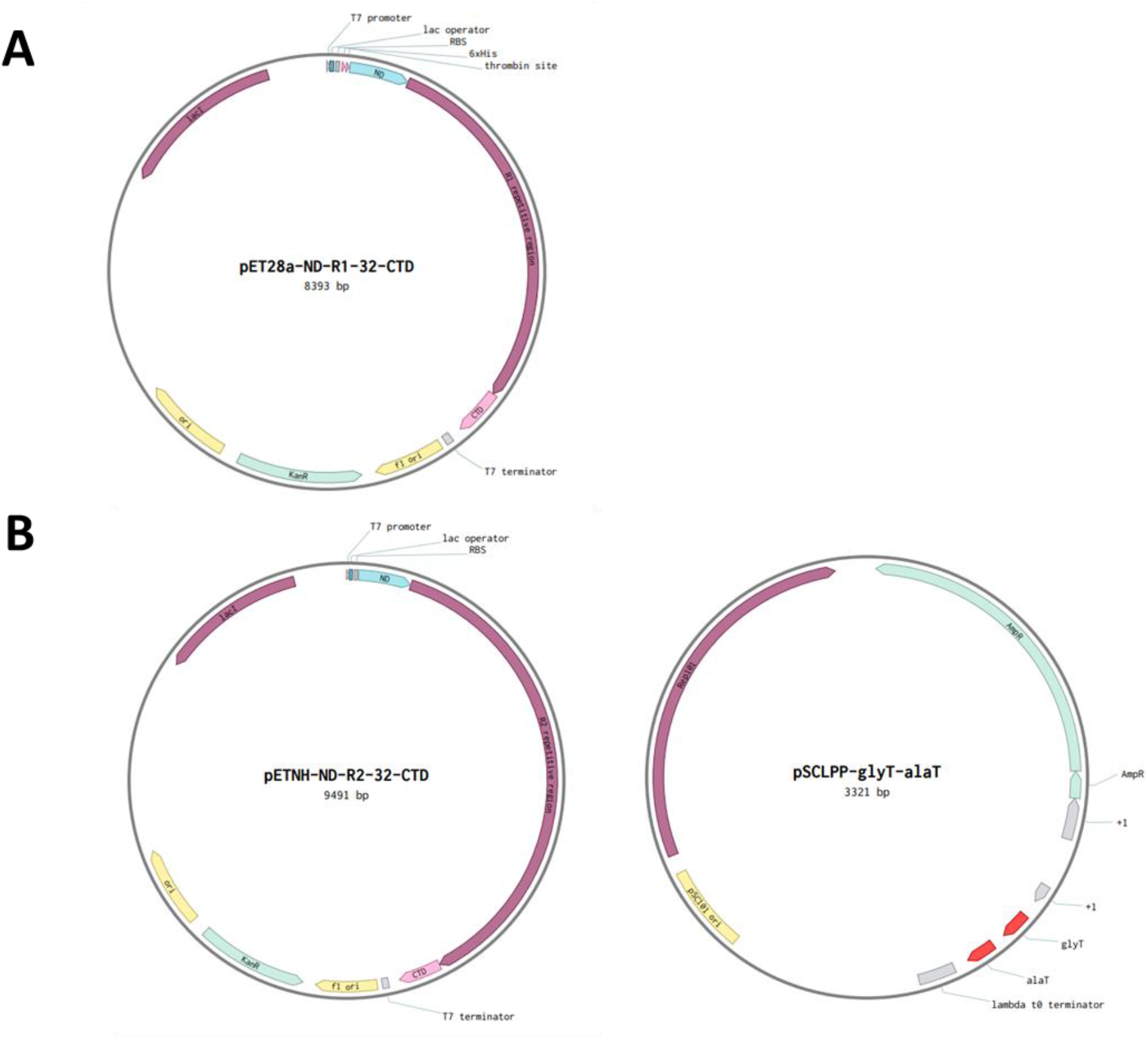
Engineered expression vectors for biosynthetic spidroins production. (A) pET28a-ND-R1-32-CTD contained R1 gene on the pET28a vector for R1 expression. (B) pENTH-ND-R2-32-CTD carried R2 gene on the pET28a vector without thrombin and his-tag expression. pSCLPP-glyT-alaT expressed glycine and alanine tRNA to enhance the silk protein expression.

### III. Preparation of mouse anti-NTD monoclonal antibody

To identify the biosynthetic silk protein for further detection, we prepared monoclonal anti-NTD antibody. Briefly, three major procedures were conducted: mouse immunization, cell hybridization, and cell selection.

In the beginning, the mouse was injected with antigen, NTD protein, for weeks. During the immunization, the mouse serum was collected every two weeks followed by Western blotting to determine the mouse antibody titer. In **Supplementary Fig. S4A**, NTD protein (15kDa) was recognized through serum and stronger signal indicated the mouse immunization against NTD was boosted.

After eight weeks immunization, the NTD antibody-producing spleen was homogenized into spleen cell suspension. The cells were fused with immortal myeloma cell line to generate hybridoma cell lines, followed by HAT (hypoxanthine-aminopterin-thymidine) medium selection. The normal myeloma cell was unable to synthesis DNA under HAT medium. Normal spleen cell was unable to divide cell immortally. Only the spleen-to-myeloma hybrid cell was able to survive in HAT medium.

After HAT medium selection, survived hybridoma were screened for the anti-NTD antibody productivity. We harvested the cultured medium from hybridoma cells and analyzed with ELISA for antibody efficacy evaluation. The ELISA results were shown in Supplementary Fig. S4B. The ELISA plates were coated with biosynthesis NTD protein or *N. pilipes* silk protein. When the medium contains anti-NTD antibody, the antigens are recognized and displayed the titer with TMB (3, 3’, 5, 5’-tetramethylbenzidine) at OD_450_. The B5, C1D1 colonies showed the high recognition ability against both NTD protein and spider silk protein. Specifically, the C6 colony developed the balance recognition ability between the two antigens. The C6 colony was selected for further single-cell isolation. Finally, the medium with monoclonal anti-NTD antibody was stored in 50% glycerol buffer at −20°C.

**Supplementary Figure S4.**
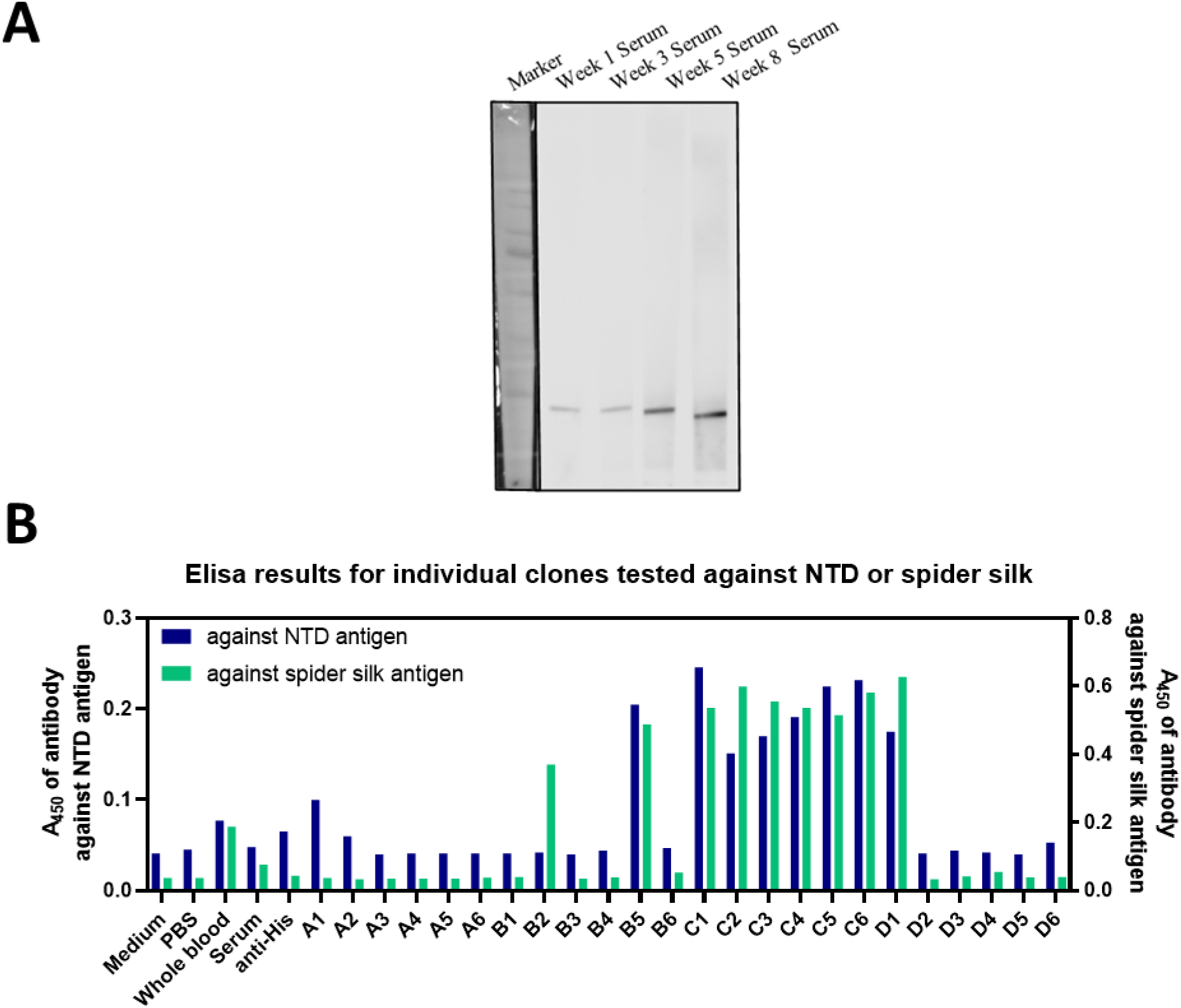
Construction of mouse anti-NTD monoclonal antibody. (A) Western blotting of serums from different weeks against biosynthetic NTD (15kDa). (B) ELISA titer tests of mouse anti-NTD antibody against two antigens, N-terminal domain and *N. pilipes* spider silk. The colony ‘C6’ was selected to isolated the monoclonal cell line.

### IV. Biosynthized spider silk-based material propelling systems

Along the R2materials fabrication process, an interesting phenomenon was observed: as-cast R2 films (untreated) was extremely vibrating/tumbling upon water contact. It’s unclear about the actual cause of such phenomena upon the R2’s interaction with water. It’s highly suspected to be associated with the supra-molecular reorganization or hydrogen bonding disruption. Harnessing such great potentiality resided in the silk materials, an autonomous silk-based motor system was created. Specifically, the swimming tadpole model was prototyped, where the head was made of an un-dissolvable color tape and the tail was made of the R2 films. That is the R2 membrane dissolution process propels the movement of color tape (Supplementary Fig. S5A).

To demonstrate the moving, each tail was attached to the color tape. The movement of the samples on the water’s surface was recorded. The sample that was placed on the water’s surface was recorded as the zero-second time point. The movies were recorded for 55 seconds, whereas photographs were captured every 5 seconds. The red arrows labeled the moving of color tape. The color tape with R2 tail was repelled while color tapes in R1 and the control tape group remained stationary (Supplementary Fig. S5B-S5D).

The moving distance of color tapes were demonstrated in the Supplementary Fig. S6. The labeling tape with R2 tail traveled about 60 mm, whereas R1 and tape tail traveled about 10mm. Combinded with the crystallnility results(Fig. 4E and 4F), it is suggested that the H-bond was iteracting with random coil in R2, triggering the vibration and even the kinetics during the decomposition process. R1 with high crystallinity suggested the intact structures that water molecule are neither to vibrate the film nor disrupt the structure.

**Supplementary Figure S5.**
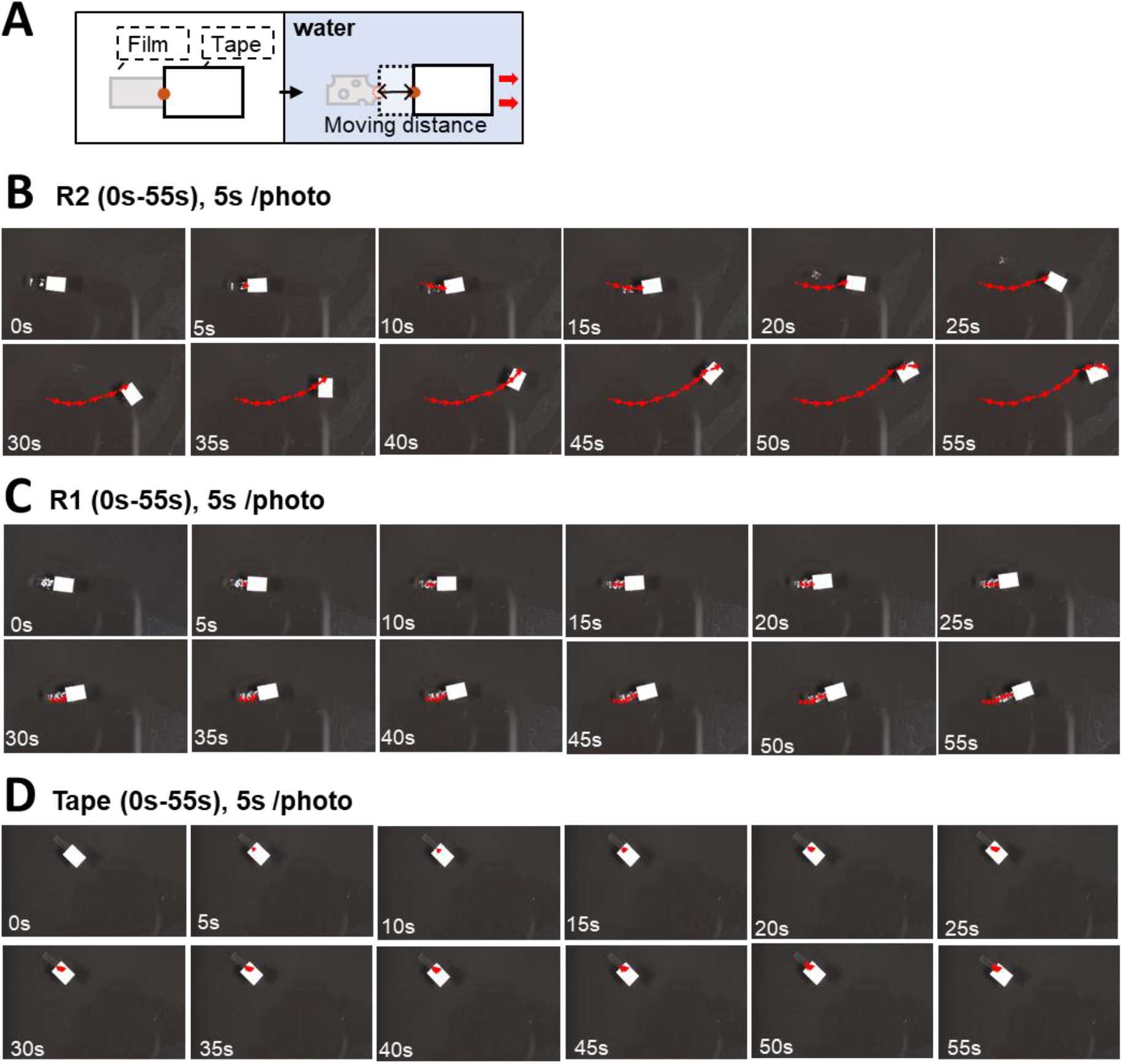
Biosynthetic spidroin-based tape propelling in water. (A) After the film was attached on the color tape, the tape was placed on the water surface to record the tape movement. (B-D) The time lapse of R2 (B), R1 (C), tape (D) on the water’s surface. Moving directions and distances were depicted by red arrows.

**Supplementary Figure S6.**
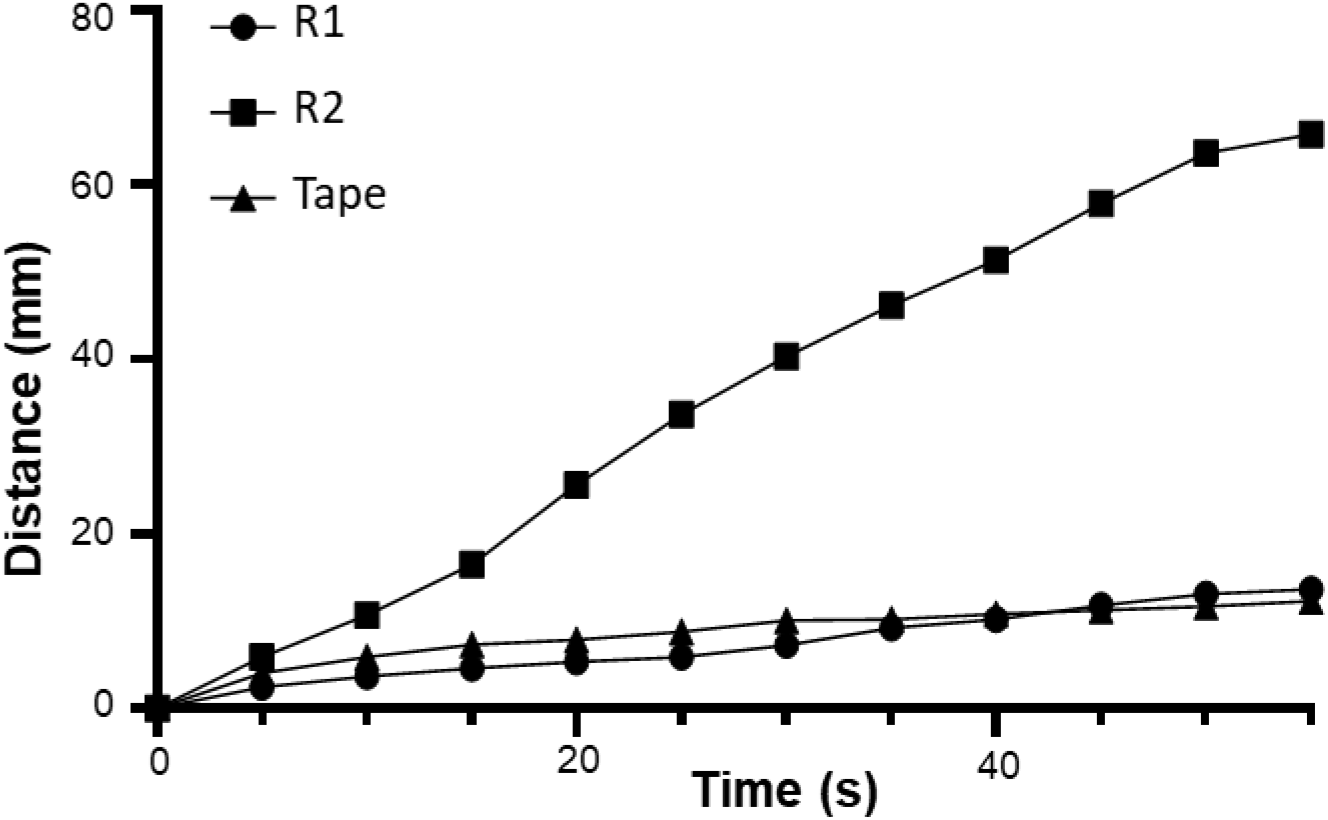
Migration distance of biosynthetic spidroin films in water. The color tape traveling distance was recorded and calculated into overall distance in every five seconds.

### V. Analysis of protein structure of biosynthetic spidroin

Along the water propelling system, the film solubility was increased after alcohol treatment. The detail was depicted in main text. It is unclear whether how the protein structures of biosynthetic spidroin films varied within MA spidroins after the post-treatments. The stability was highly suspected to be attributed to the crystallinity. Herein, the X-ray diffraction (XRD) and Fourier-transform infrared spectroscopy (FT-IR) was introduced to recognize the crystallinity. The data collection was described in Materials and Methods.

After data collection, each pattern was analyzed by gaussian deconvolution in OriginPro 9.0. The deconvolution patterns of XRD and FTIR were illustrated in Supplementary Figs. S7A-S7J and S8A-S8J.

In XRD (**Supplementary Fig. S7A-S7J**), peaks represented β-sheet were shown at ∼13.5° (Green dots) and ∼16°(Orange dots), whereas random coil peaks was demonstrated at ∼7°(light blue dots),10-12°(red dots) and 13°(dark blue dots). The dark yellow dot was represented the integration of all peaks. The crystallinity was the ratio of β-sheet area to total area; the percentages were displayed in the top right of the plots. R1 crystallinity remained about 40% in as-cast and after EtOH100, whereas R2 crystallinity increased from 20% to 33%. It is highly suspected that the crystallinity of R2 was programmable under post-treatment while R1 was almost fixed. Next, the typical post-treatments, alcohol and water vapor annealing, were introduced to R2. In alcohol treatments, the crystallinity of R2 remained over 30%. Surprisingly, the crystallinity was controlled by vapor temperature and treatment duration. For further investigation, FT-IR studies was conducted.

In FTIR (Supplementary Fig. S8A-S8J), peaks represented β-sheet were shown in wavenumber ∼1622 cm^-1^ (Green dots) and ∼1700 cm^-1^ (Orange dots); side chain was demonstrated in wavenumber 1610 cm^-1^ (Red dots); random coil was displayed in wavenumber 1649 cm^-1^ (dark blue dots); b-turn was shown in wavenumber 1661 cm^-1^ (light blue dots) and 1683 cm^-1^ (pink dots). The dark yellow dot represented the integration of all peaks. The crystallinity ratio was calculated by normalizing the β-sheet area to total area; the percentages were displayed in the top right of plots. Similar to the trends show in XRD, the measurement of crystallinity in R1 before and after EtOH treatment remined while R2 was tunable. In ethanol treatments, crystallinity levels remained above 30%. Moreover, the crystallinity was also determined by the WVA temperature and treatment duration.

Overall, the crystallinity of R2 films were programmable while R1 films were stable. The programmable R2 films were suspected to be associated with the β-sheet formation processes, which are dependent on the dynamics of environmental factors, time and energy input. Such programmability of R2 has inspired us to investigate further. The following experiment was depicted in Fig. 5.

**Supplementary Figure S7.**
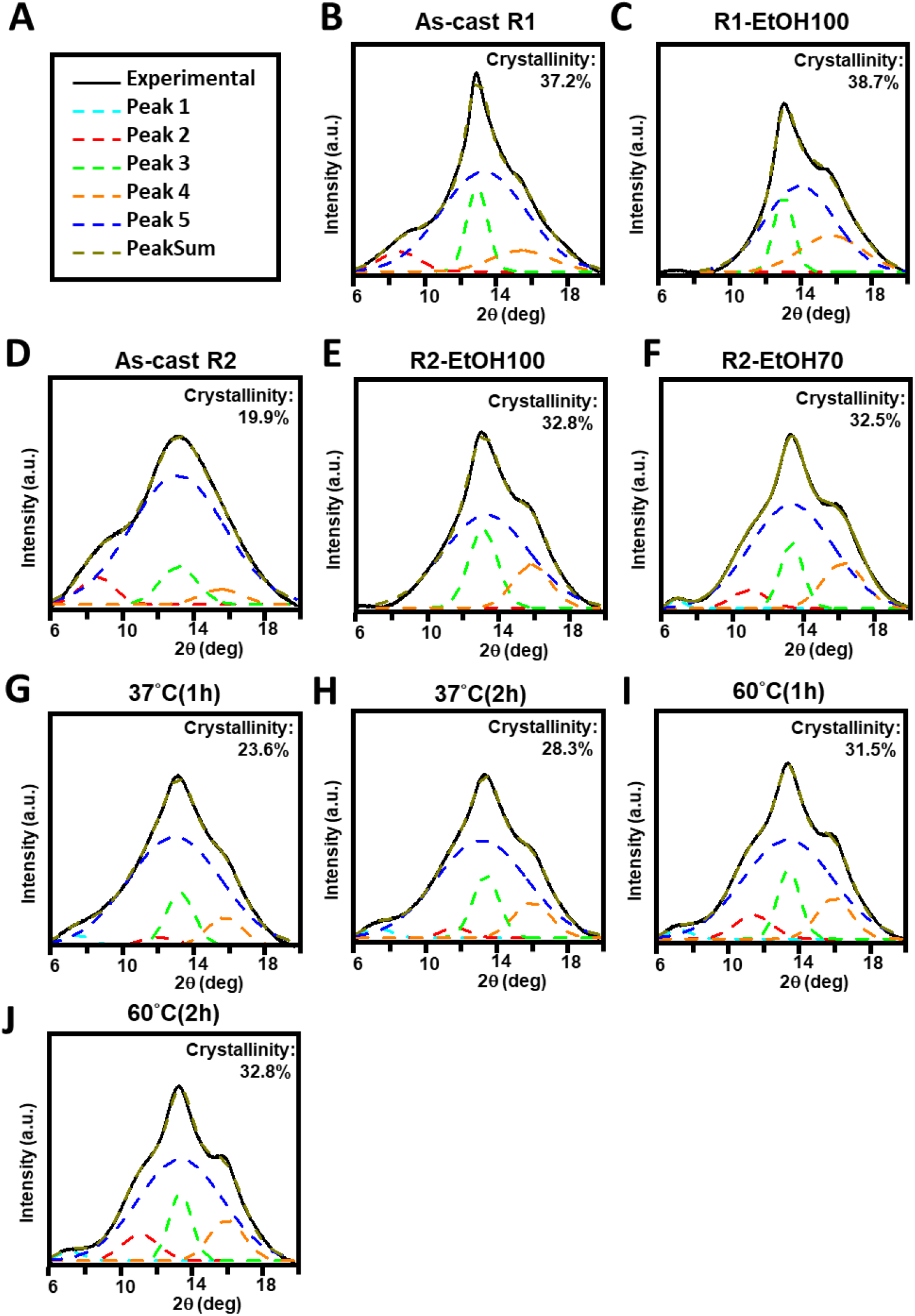
XRD structural characterization of biosynthetic spidroin films. (A) The symbol of the line. Black line: Collected data; Light blue, red, and dark blue: Random coil; Green and orange: β-sheet; dark yellow: The integration of peaks. (B-J) Different treatment of spidroin. Each pattern was represented As-cast R1 (B), ETOH100 treated R1 (C), As-cast R2 (D), ETOH100 treated R2 (E), ETOH70 treated R2 (F), WVA 37°C (1h) for (G) and WVA 37°C (2h) for (H), 60°C WVA (1 h) for (I) and 60°C WVA (2 h) for (J).

**Supplementary Figure S8.**
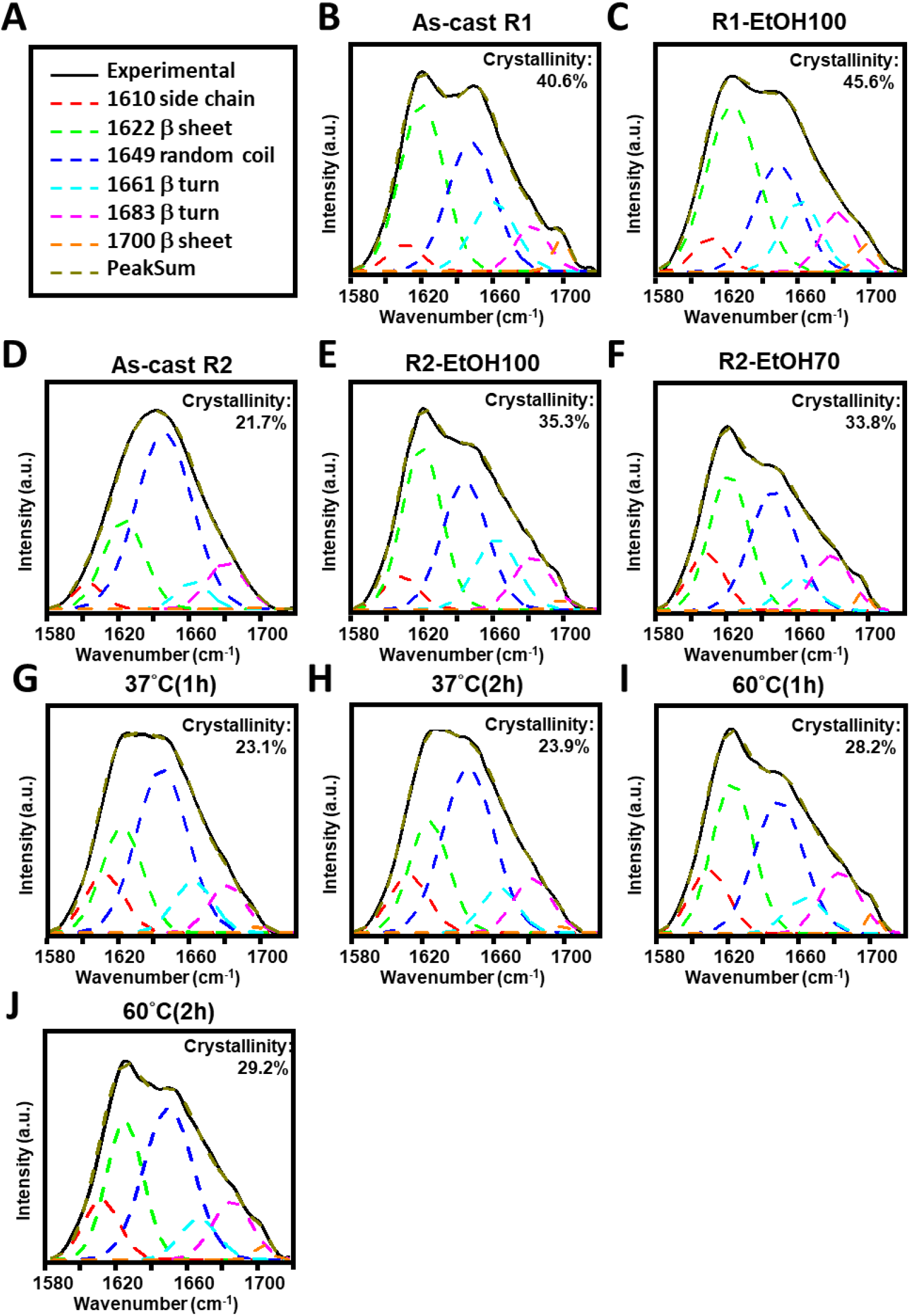
FTIR structural characterization of biosynthetic spidroin films. (A) The symbol of the line. Black line: Collected data; Red dots: Side chain; Dark blue dots: Random coli; Light blue and pink dots: β-turn; Green and orange: β-sheet; dark yellow: The integration of peaks. (B-J) Different treatment of spidroin. Each pattern was represented As-cast R1 (B), ETOH100 treated R1 (C), As-cast R2 (D), ETOH100 treated R2 (E), ETOH70 treated R2 (F), WVA 37°C (1hr) for (G) and WVA 37°C (2h) for (H), 60°C WVA (1 h) for (I) and 60°C WVA (2 h) for (J).

### VI. Fabrication and mechanical characterization of novel graphene-doped spider silk materials

Following the graphene-doped R2 film was created, the effects of graphene doping on the performance of R2 materials was examined. Both mechanical strength and cohesive repairing were evaluated. To characterize the strength variation in graphene-doped R2 (denoted R2G), mechanical properties of R2G films were conducted. Additionally, we also investigated the healability of the R2G as compared to R2 alone.

In the beginning, we compared non-treated conditions, as-cast (denoted as As) and cut-and-healed as-cast film (denoted as As-H) groups. The strength of graphene-contained R2 film was comparably similar (p>0.05). Next, we studied the 37°C WVA (2hr) film. The similar effect was observed in WVA groups, with no statistical difference in comparison to the control group. Also, the recovered strength of cut-and-healed R2G film was similar to the uncut one. It showed that graphene-doped R2 film had a good ability to self-repair, as comparable to the standard R2 films.

**Supplementary Figure S9.**
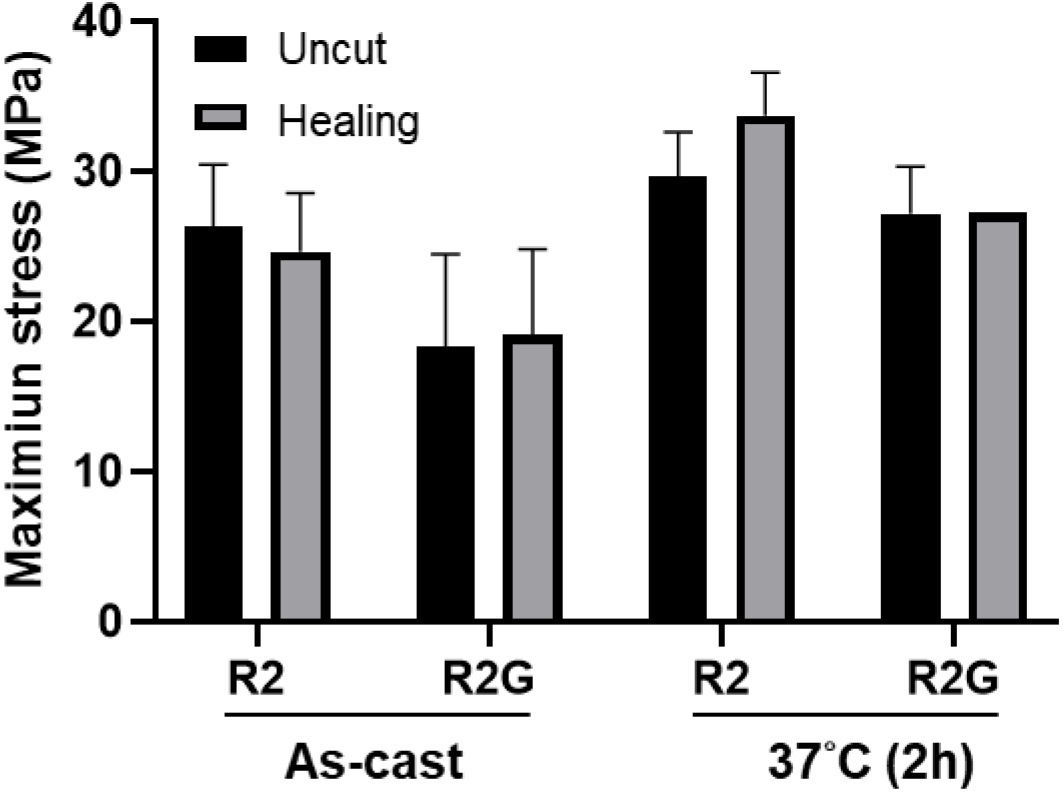
Tensile strength evaluation of graphene-doped biosynthetic spidroin films. The maximum strength values of the as-cast R2 film and graphene-doped R2 film prior to and after the cut-and-healed processes were determined.

### VII. Repairable biosynthetic spidroin-based flexible logic gates

Based on R2G mechanical properties, the soft-fabrics were generated. Harnessing the spidroin-based materials, the complex fabrication process was also establised.

1Dfabric demonstrated the healability of R2G and the logic gate table was shown in Supplementary Fig. S10 and Supplementary Table S1. For 2D fabric, combing R2G with R2 did significantly increase the complexity and utility of the system (Supplementary Fig. S11A). The 2D fabric was installed in the device for the circuit evaluation (Supplementary Fig. S11B). The collected outputs were shown in Supplementary Fig. S11C. The output was determined by: Cutting, healing, graphene-dope R2 (Diverter), and electricity (e^-^). Each panel was corresponding to specific operation in Supplementary Fig. S11C and the collective logic gate table was shown in Supplementary Table S2.

Next, 2Dfabric was self-healed to create 3D fabric (Supplementary Fig. S12A). The whole setup and the circuit diagram illustration were shown in Supplementary Fig. S12B. Besides the original 2D fabric properties, the variable for orientational alignment was introduced in 3D fabric. Specifically, tweaking, followed by the healing process can cause the misalignment of 3D structures, rendering a disconnected circuit. That is, the output was determined by: cutting, healing, tweaking, graphene-dope R2 (Diverter), and electricity (e^-^). Each panel was corresponding to specific operation in Supplementary Fig. S12C and the final logic gate table was shown in Supplementary Table S3.

**Supplementary Figure S10.**
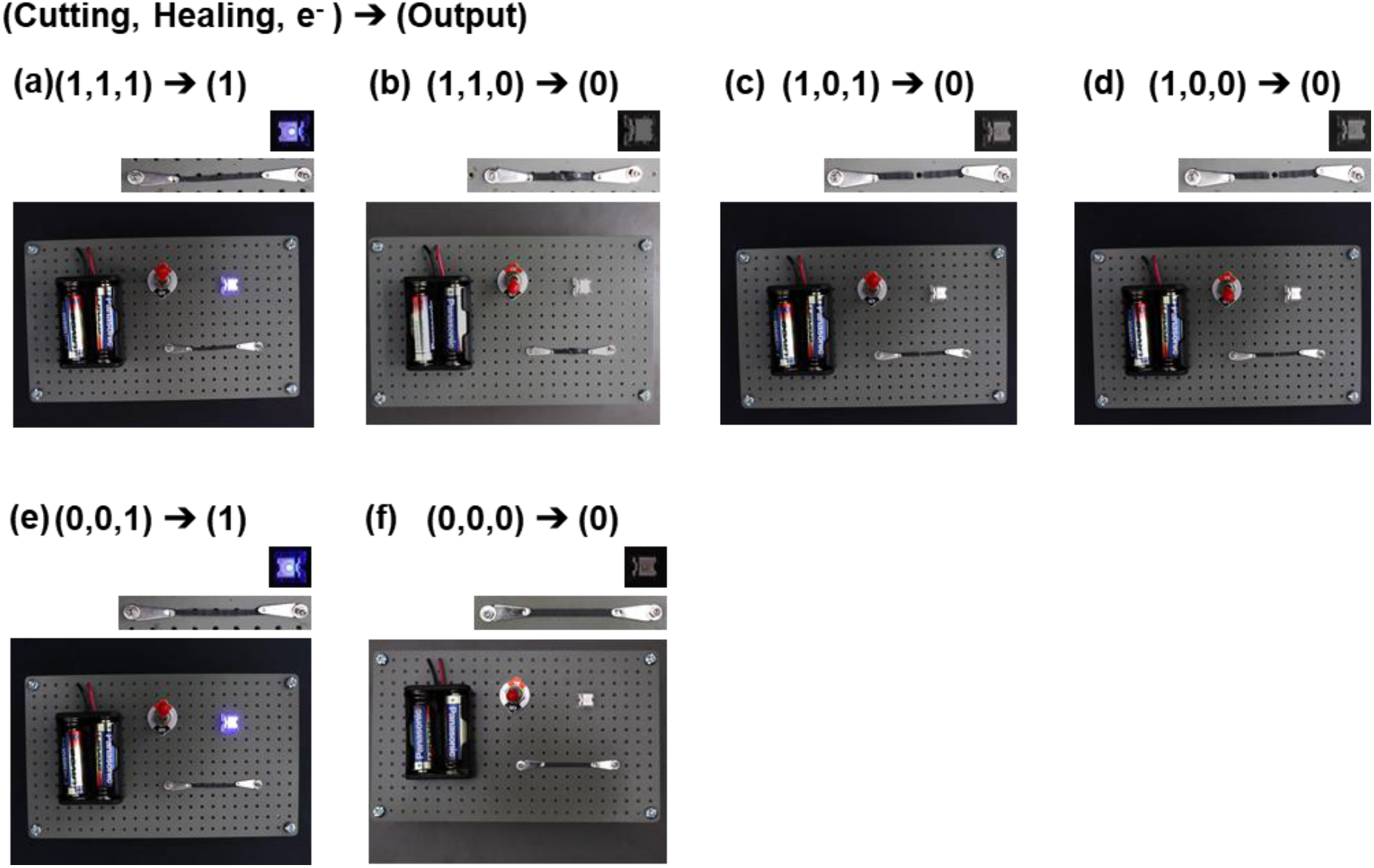
Repairable biosynthetic spidroin-based 1D logic gates.

**Supplementary Table S1.**
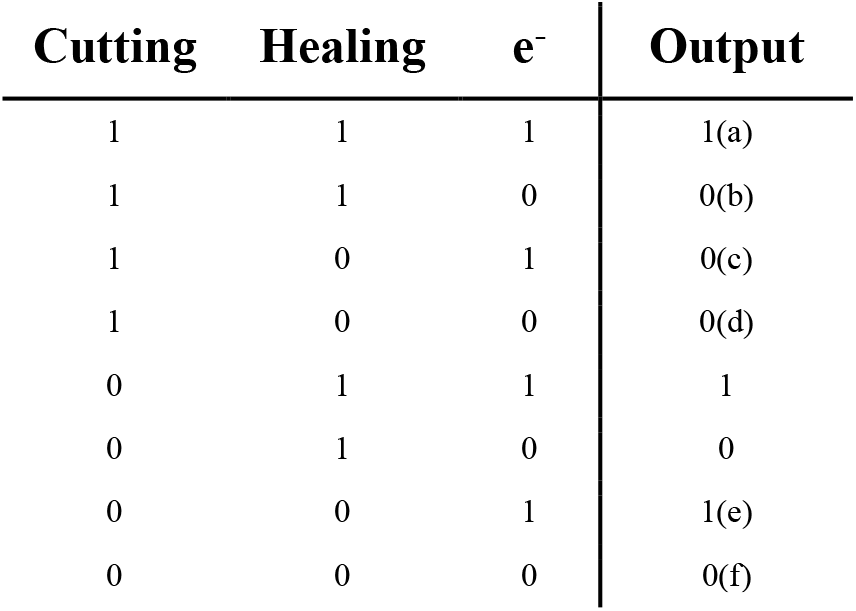
Truth table of 1D biosynthetic spidroin-based logic gate circuitry.

**Supplementary Figure S11.**
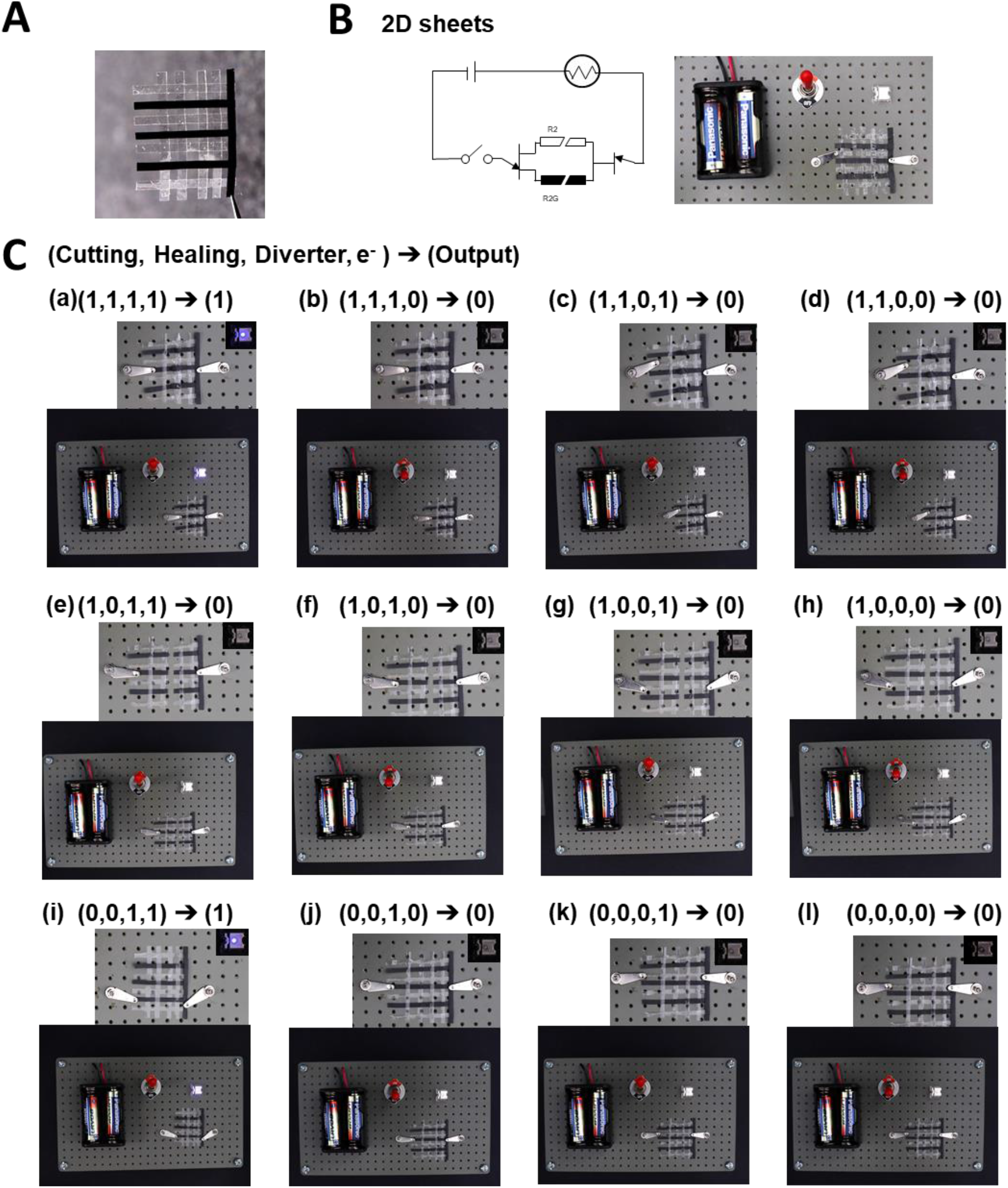
Repairable biosynthetic spidroin-based 2D logic gates. (A) Fabrication of spider spidroin-based 2D sheet. (B) Setup and circuit diagram of 2D fabric system. (C) The operations and output of 2D fabric.

**Supplementary Table S2.**
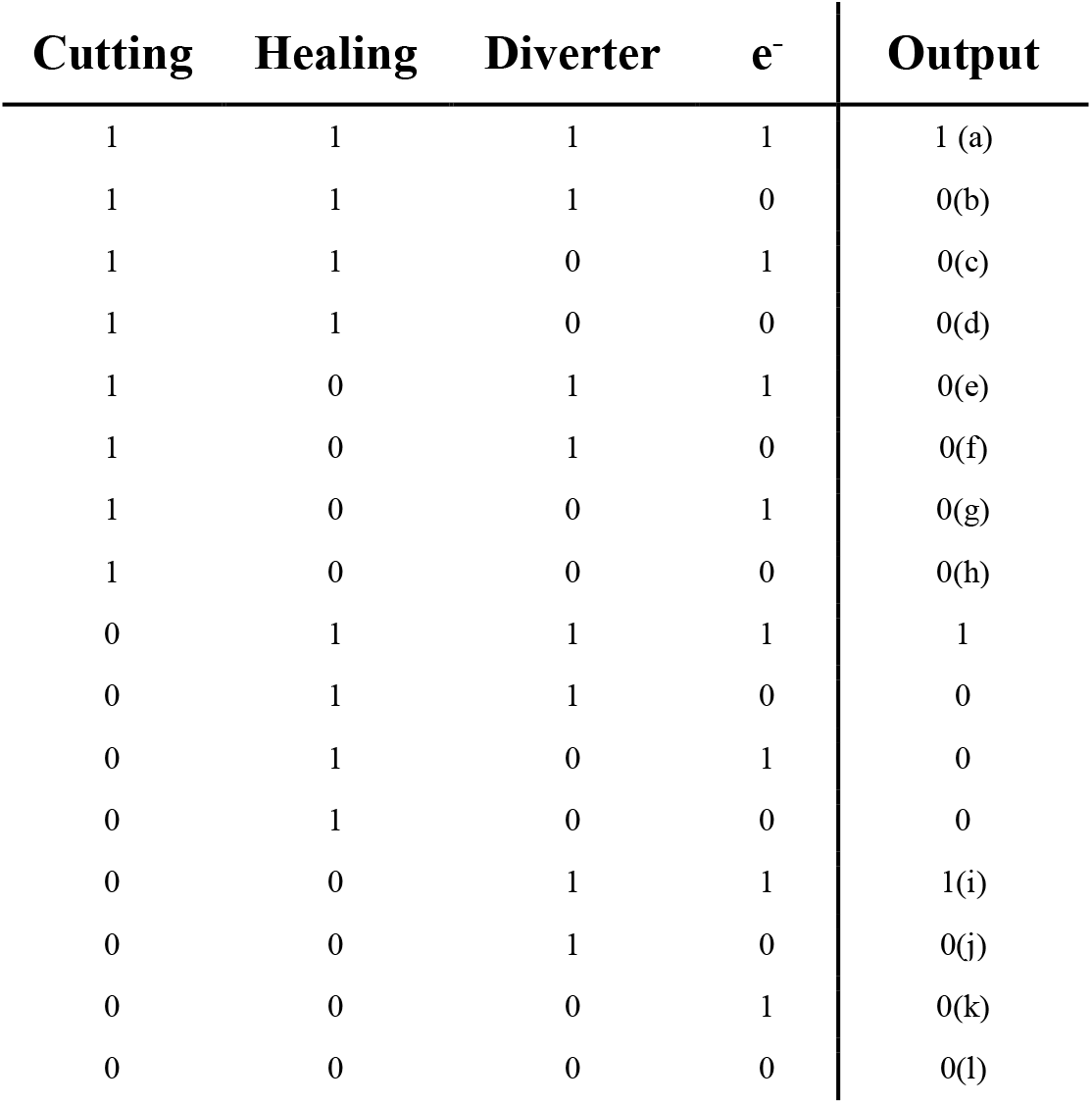
Truth table of 2D biosynthetic spidroin-based logic gate circuitry.

**Supplementary Figure S12.**
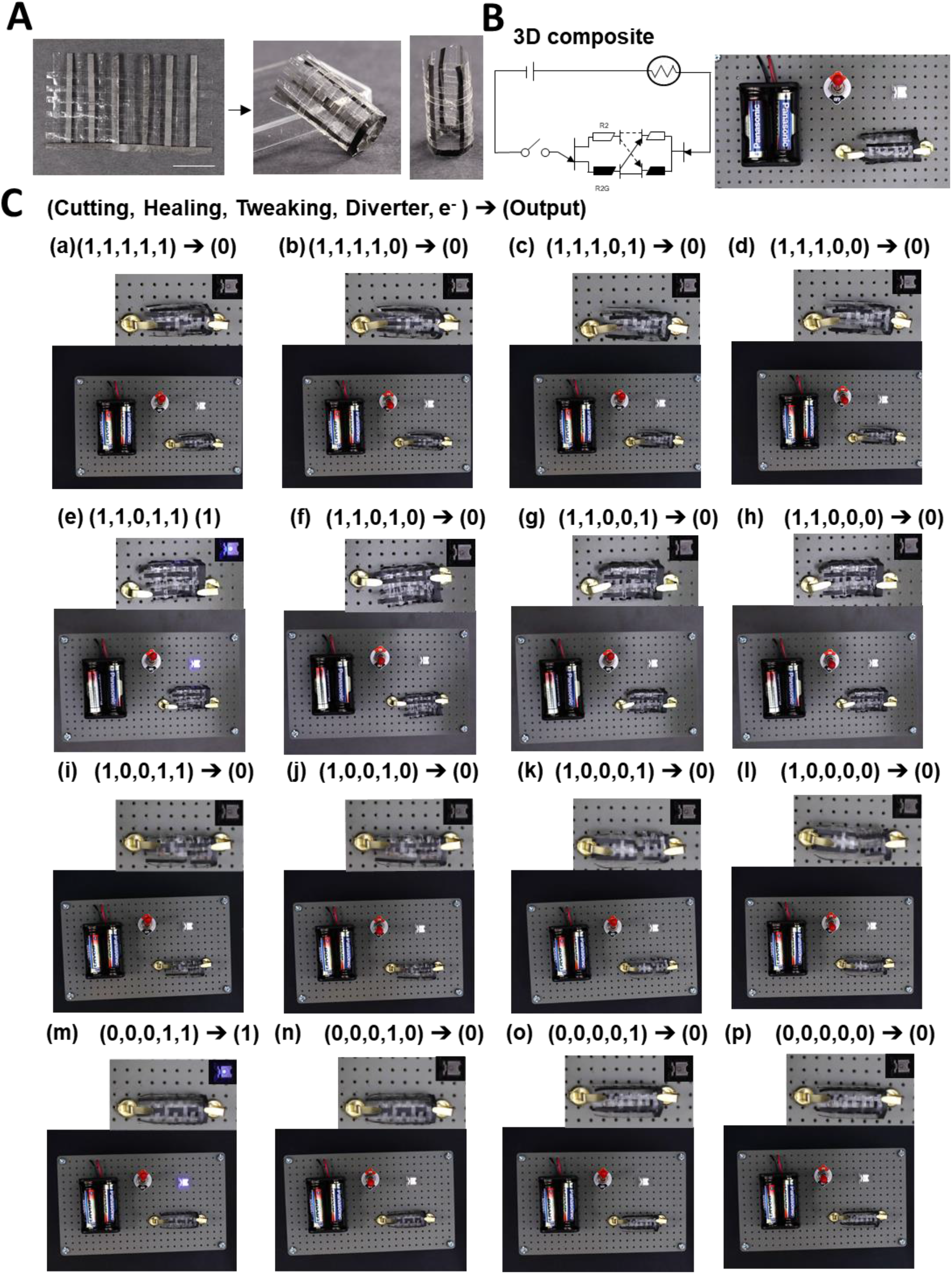
Repairable biosynthetic spidroin-based 3D logic gates. (A) Fabrication of spider spidroin-based 3D fabric. (B) Setup and circuit diagram of 3D fabric. (C) The operations and output of 3D fabric.

**Supplementary Table S3.**
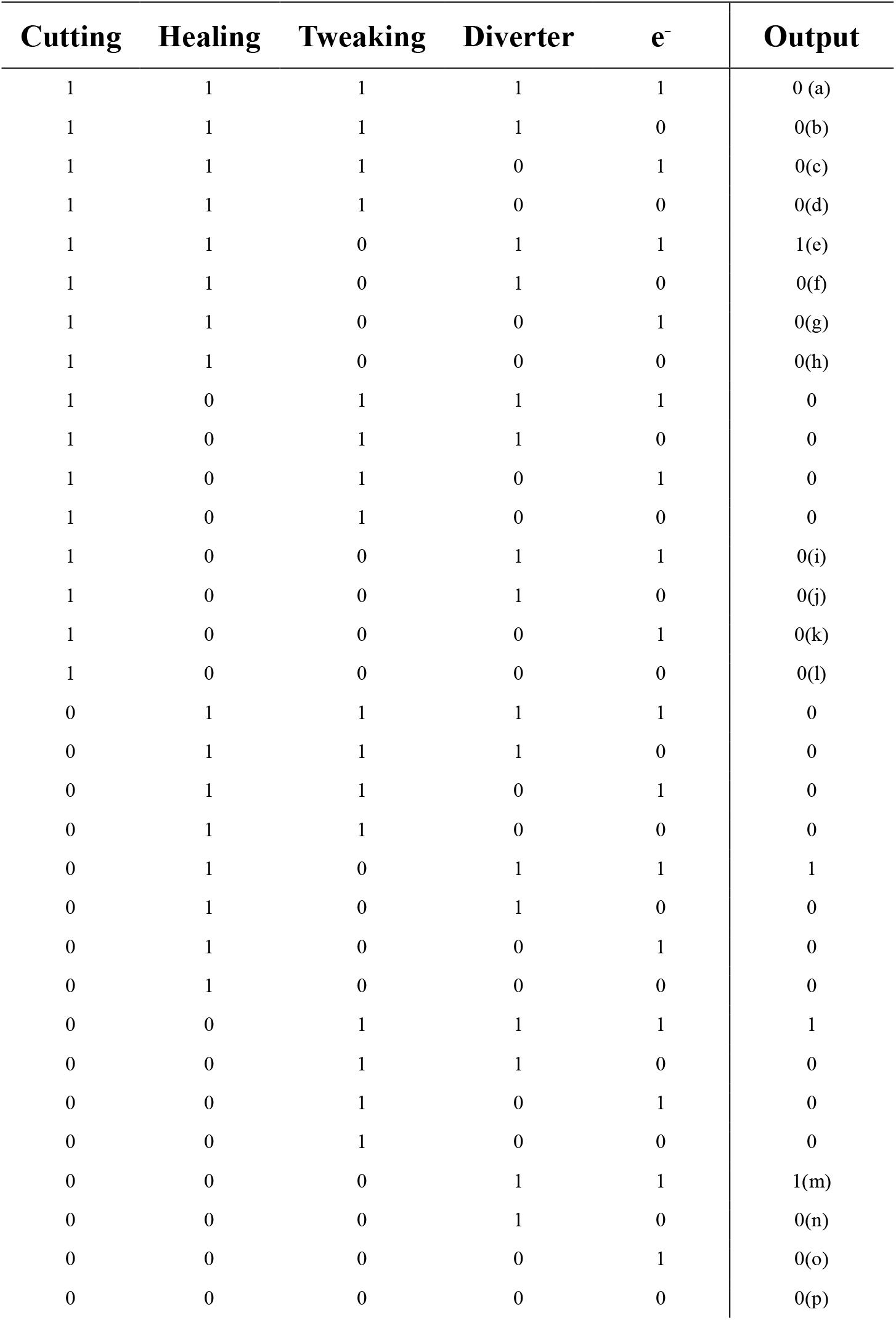
Truth table of 3D biosynthetic spidroin-based logic gate circuitry.

### VIII. Device setup for the fabricated spider silk-based biorecognition sensor

Harnessing reversible healability, WVA 60°C (1h) R2, the materials were exploited to be integrated into a smart finger/skin recognition biodevice. Specifically, the R2G film was cut into small pieces and warped the wire on the R2-based ring (**Supplementary Fig. 7C**). The R2G film served as probes to detect the surface resistance of objects. Fingers’ resistances were shown in **Supplementary Table S4** while electrical conductors’ resistances were displayed in **Supplementary Table S5**. Insulator resistances were beyond the detection range of the Keithley 2400 source/meter. Consequently, the insulators’ resistances were referred to Arc resistance [5] and was demonstrated in **Supplementary Table S6**.

Along resistance database, the spider silk-based biorecognition sensor was created. Biorecognition sensor was designed to identify the object and display the readout on the LCD. The Arduino sketch code was displayed in **Supplementary Fig. S13**. In the hardware design, interface of Arduino was represented in **Supplementary Fig. S14A** and complete coding was shown in **Supplementary Fig. S14B**. Finally, the biorecognition sensor was created in **Supplementary Fig. S14C**. The further test was depicted in **Supplementary Fig. 7E**.

**Supplementary Table S4.**
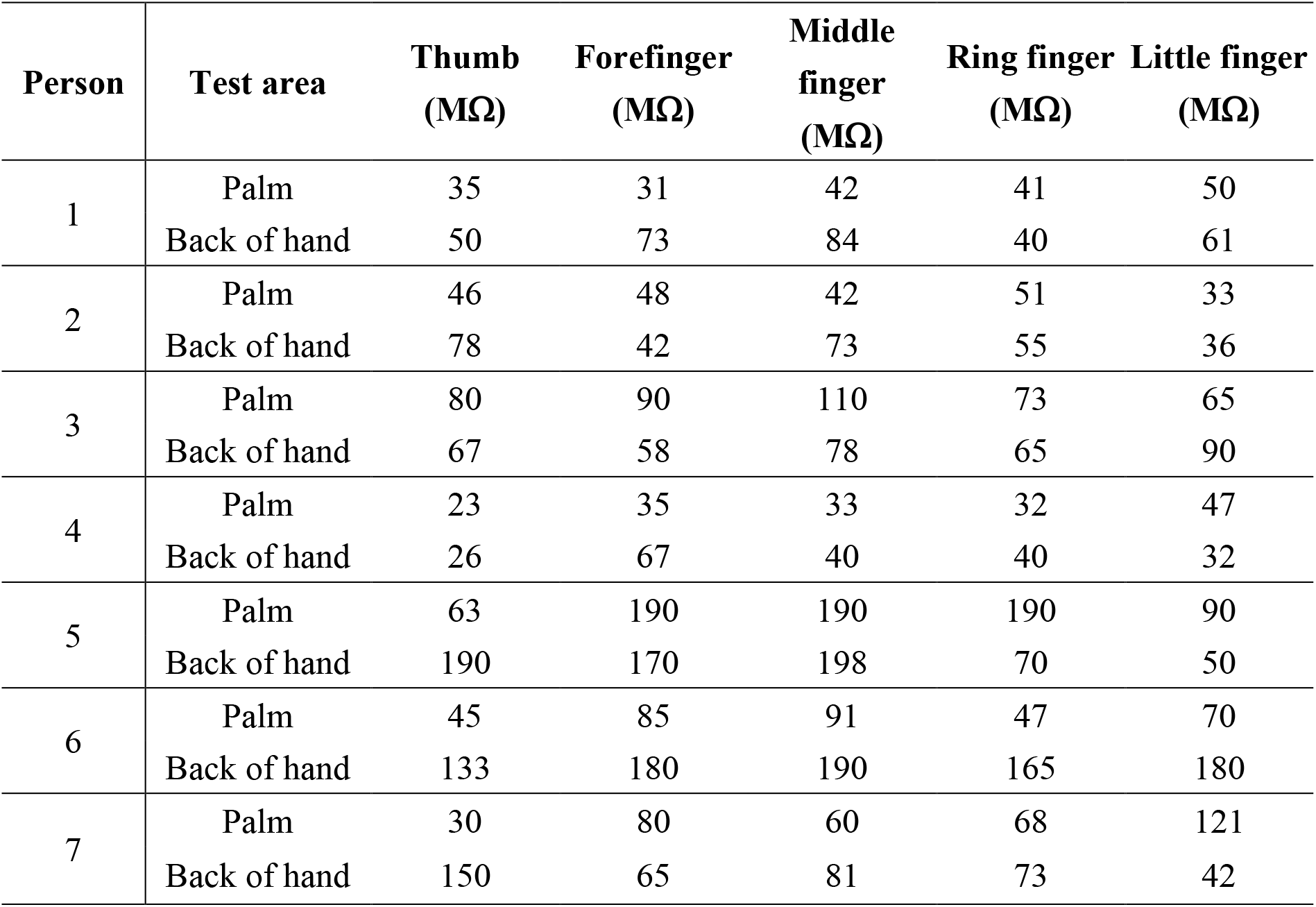
Resistance measurements of human fingers via the biosynthetic spidroin ring detector.

**Supplementary Table S5.**
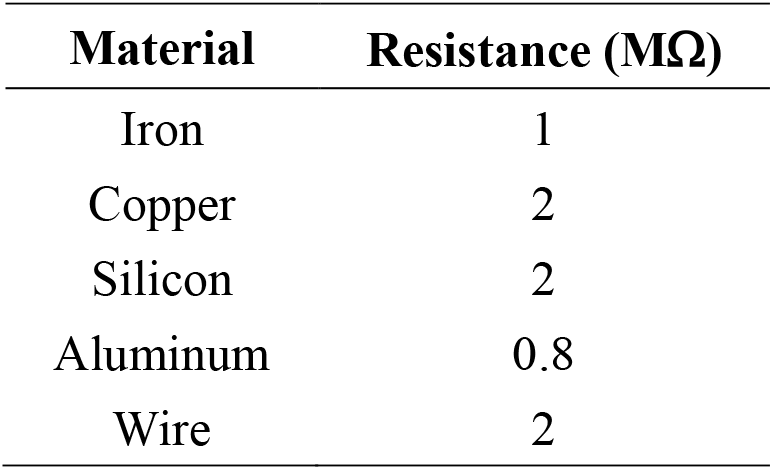
Resistance measurements of conductors s via the biosynthetic spidroin ring detector.

**Supplementary Table S6.**
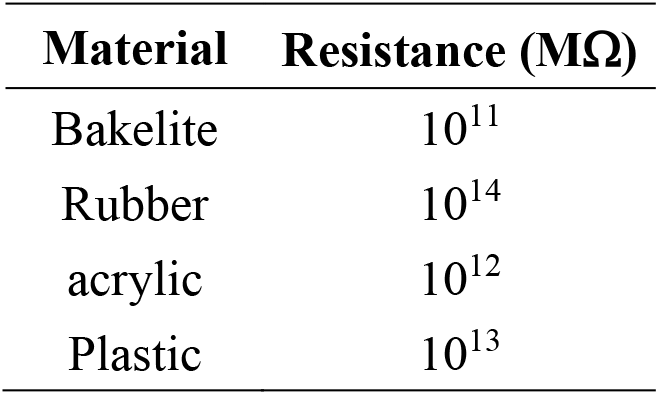
Reference Resistances of insulator materials [5].

**Supplementary Figure S13.**
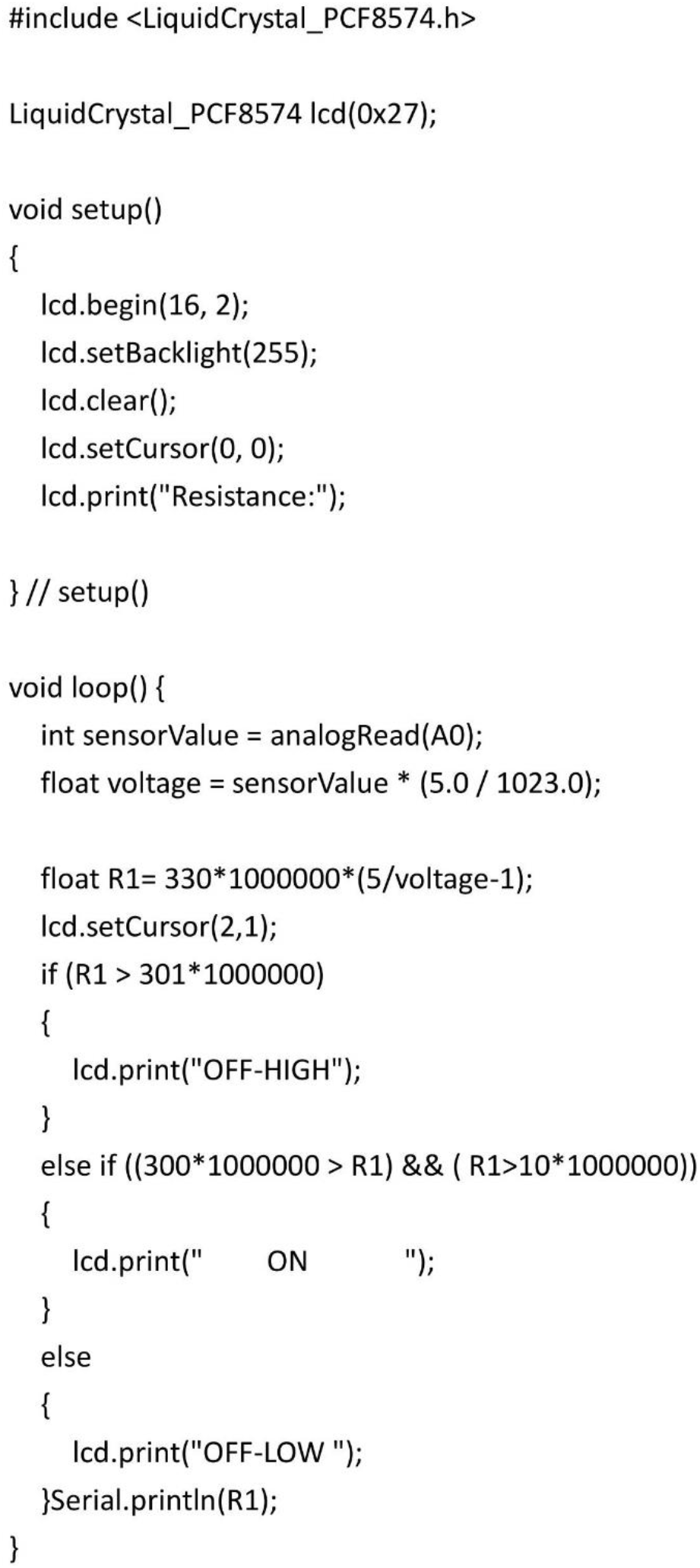
Arduino sketch code of biosynthetic spidroin-based biorecognition sensor.

**Supplementary Figure S14.**
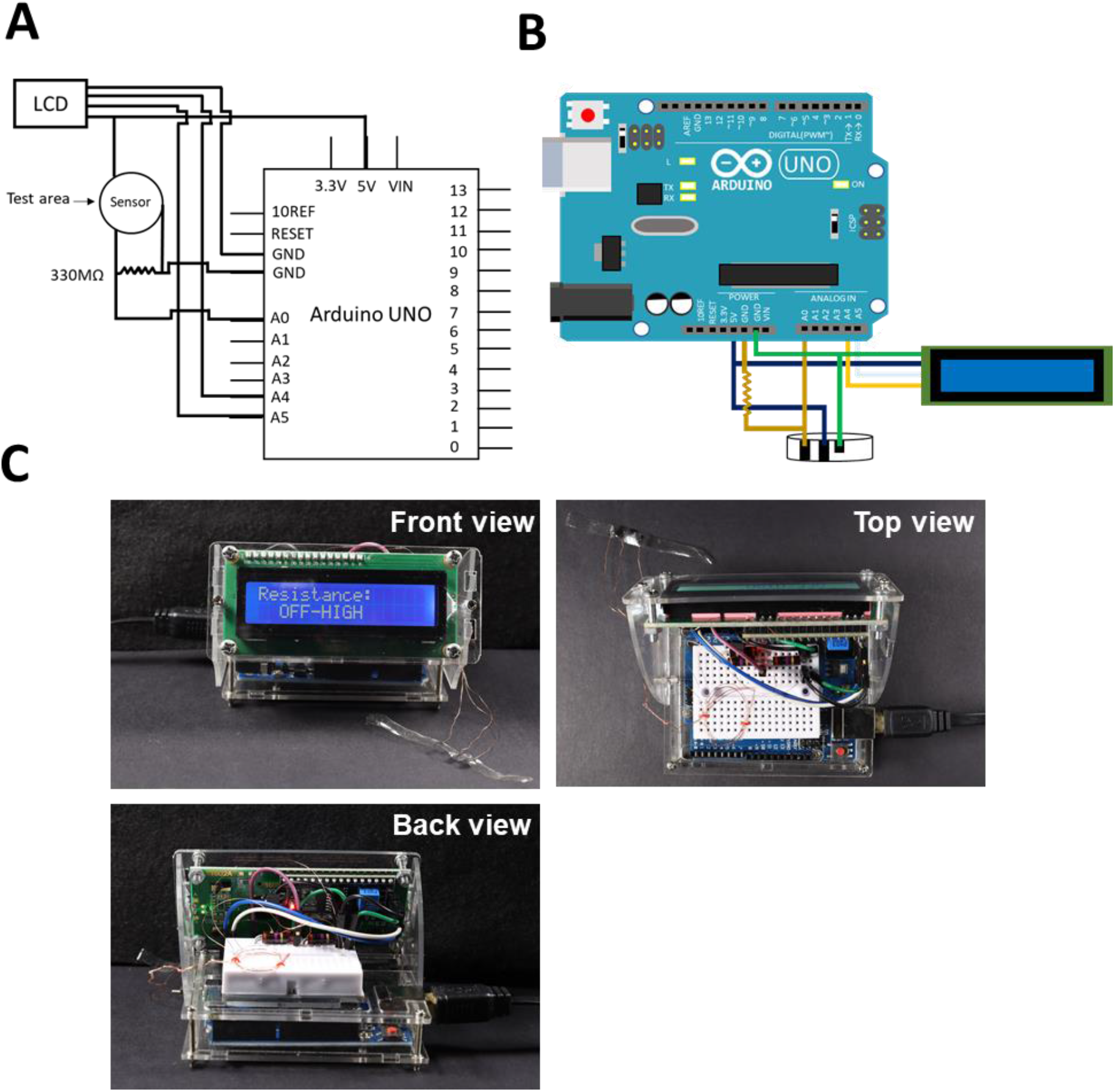
Device configuration for biosynthetic spidroin-based biorecognition sensor. (A) Interface of the Arduino (B) Wiring diagram for spider silk-based biorecognition sensor (C) Device front view (top left), top view (top right), and back view (bottom)

