## Supplementary data for "Self-Healable Spider Dragline Silk Materials"

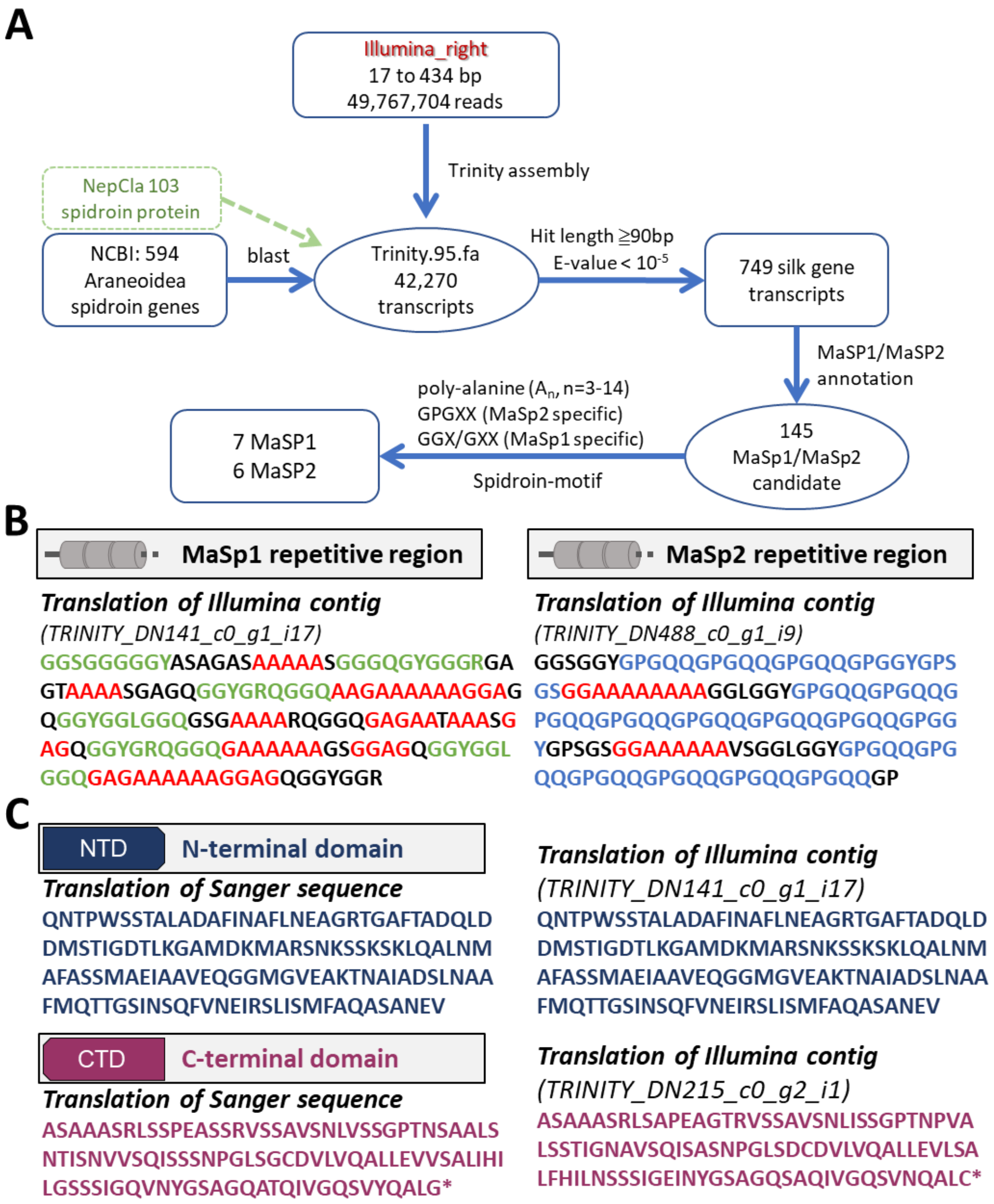

**Supplementary Figure S1. Sequence analysis of MA spider silk.** (A) The flow chart of candidates sorting. (B) Partial sequencing results of MaSp1 and MaSp2 repetitive regions. Structural motifs were characterized as poly (A/GA) (red), GGX/GXX (Green), and GPGXX (Blue). (C) Sanger sequence and Illumina sequence results of NTD (dark blue) and CTD (dark red).

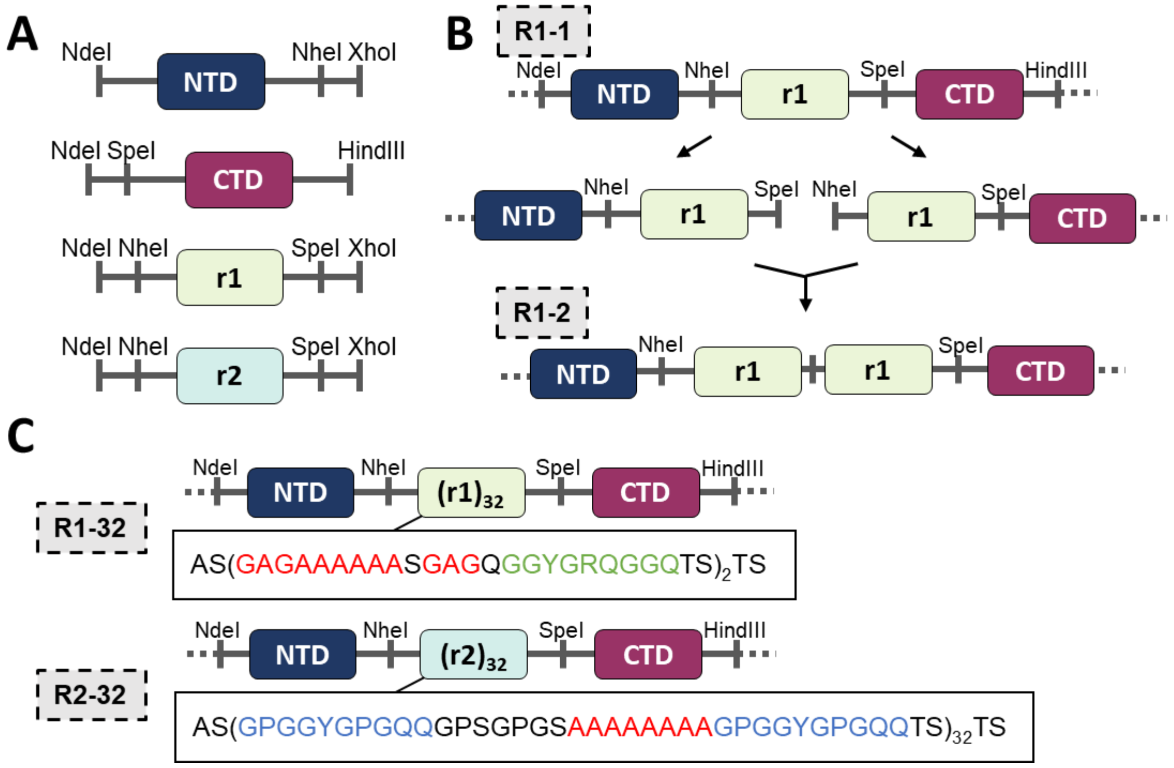

**Supplementary Figure S2. Creation of codon-optimized biosynthetic spidroins.** (A) The terminal domain and repetitive unit cassettes construction by PCR amplification. (B) Amplification of repetitive units by BioBirck restriction enzyme cloning (SpeI and NheI pairs) (C) After cloning amplification, pET28a-NTD-$(r1)_{32}$-CTD and pET28a-NTD-$(r2)_{32}$-CTD were generated. The translation of repetitive regions was displayed and the motifs were characterized by poly (A/GA) (red), GGX/GXX (Green), and GPGXX (Blue).

A

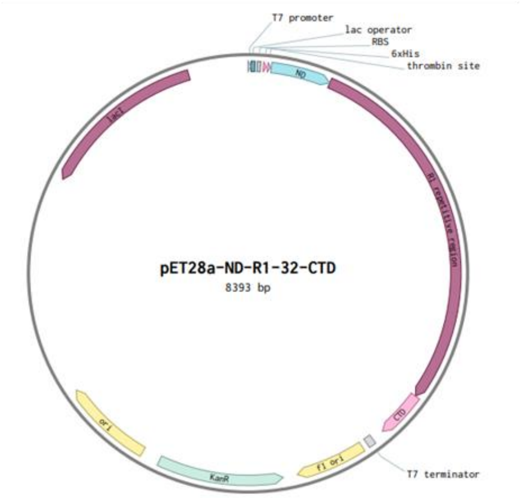

B

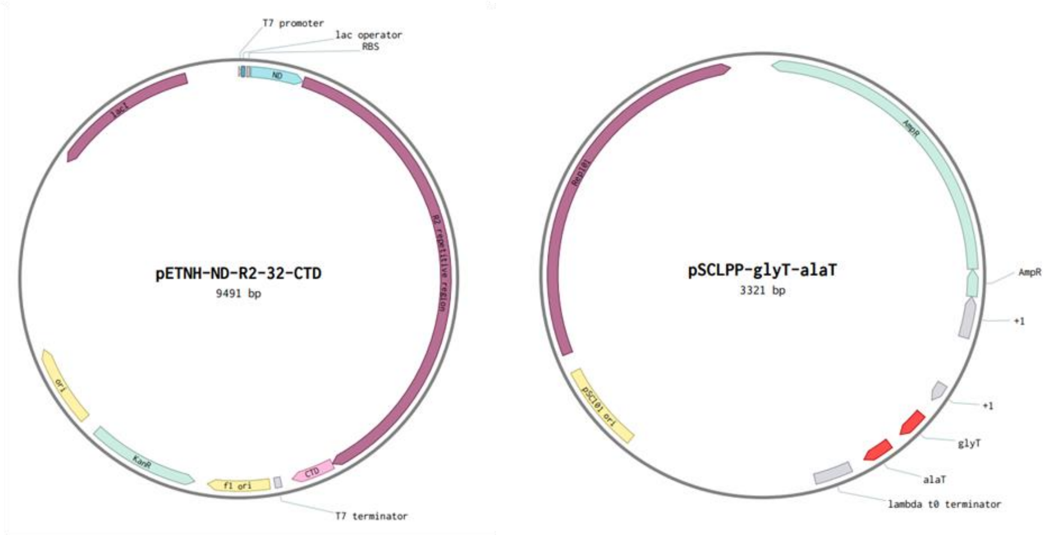

**Supplementary Figure S3. Engineered expression vectors for biosynthetic spidroins production.** (A) pET28a-ND-R1-32-CTD contained R1 gene on the pET28a vector for R1 expression. (B) pENTH-ND-R2-32-CTD carried R2 gene on the pET28a vector without thrombin and his-tag expression. pSCLPP-glyT-alaT expressed glycine and alanine tRNA to enhance the silk protein expression.

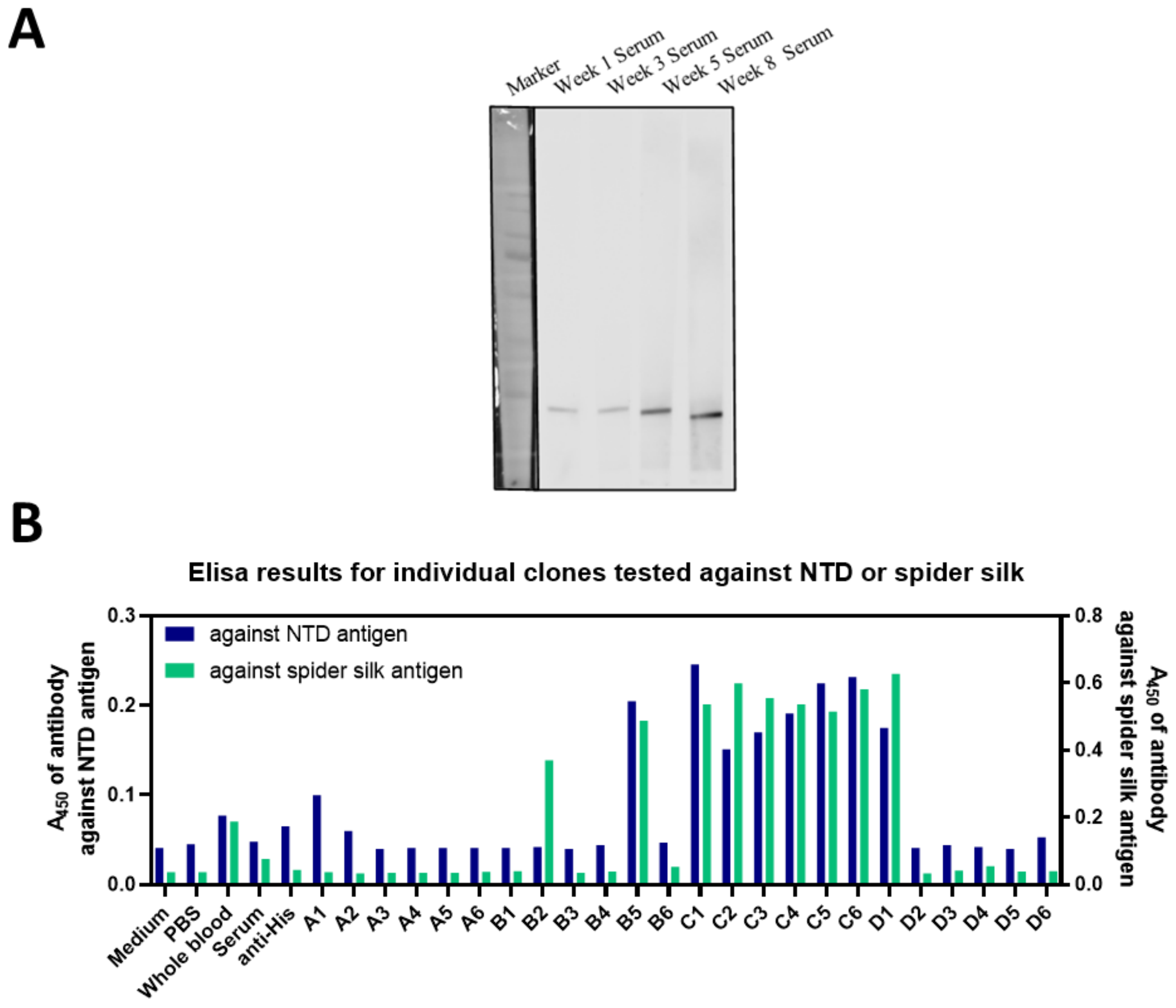

**Supplementary Figure S4. Construction of mouse anti-NTD monoclonal antibody.** (A) Western blotting of serums from different weeks against biosynthetic NTD (15kDa). (B) ELISA titer tests of mouse anti-NTD antibody against two antigens, N-terminal domain and *N. pilipes* spider silk. The colony 'C6' was selected to isolated the monoclonal cell line.

### IV. Biosynthized spider silk-based material propelling systems

Along the R2 materials fabrication process, an interesting phenomenon was observed: as-cast R2 films (untreated) was extremely vibrating/tumbling upon water contact. It's unclear about the actual cause of such phenomena upon the R2's interaction with water. It's highly suspected to be associated with the supra-molecular reorganization or hydrogen bonding disruption. Harnessing such great potentiality resided in the silk materials, an autonomous silk-based motor system was created. Specifically, the swimming tadpole model was prototyped, where the head was made of an un-dissolvable color tape and the tail was made of the R2 films. That is the R2 membrane dissolution process propels the movement of color tape (**Supplementary Fig. S5A**).

To demonstrate the moving, each tail was attached to the color tape. The movement of the samples on the water's surface was recorded. The sample that was placed on the water's surface was recorded as the zero-second time point. The movies were recorded for 55 seconds, whereas photographs were captured every 5 seconds. The red arrows labeled the moving of color tape. The color tape with R2 tail was repelled while color tapes in R1 and the control tape group remained stationary (**Supplementary Fig. S5B-S5D**).

The moving distance of color tapes were demonstrated in the **Supplementary Fig. S6**. The labeling tape with R2 tail traveled about 60 mm, whereas R1 and tape tail traveled about 10mm. Combinded with the crystallnility results(**Fig. 4E and 4F)**, it is suggested that the H-bond was iteracting with random coil in R2, triggering the vibration and even the kinetics during the decomposition process. R1 with high crystallinity suggested the intact structures that water molecule are neither to vibrate the film nor disrupt the structure.

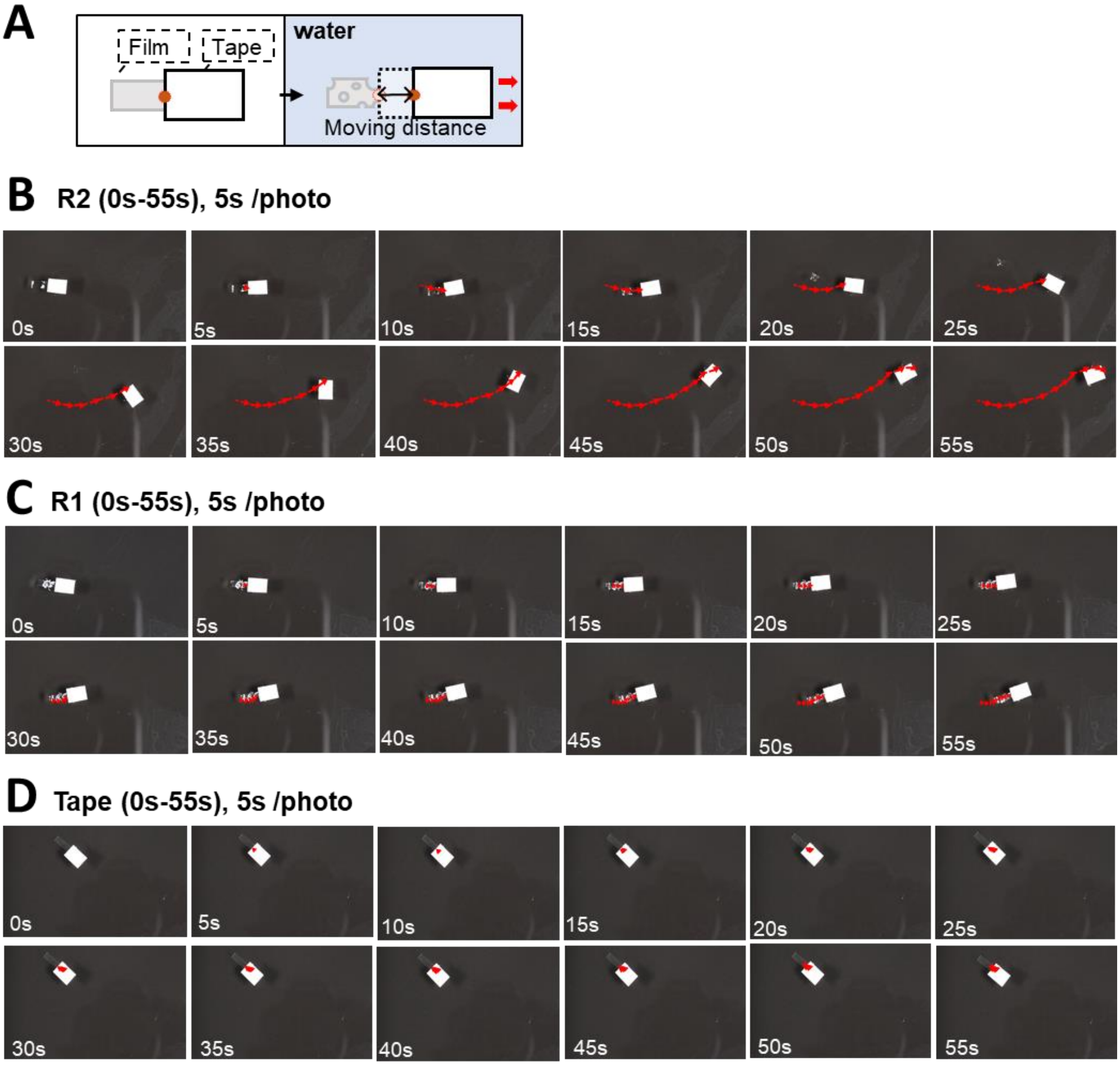

**Supplementary Figure S5. Biosynthetic spidroin-based tape propelling in water.** (A) After the film was attached on the color tape, the tape was placed on the water surface to record the tape movement. (B-D) The time lapse of R2 (B), R1 (C), tape (D) on the water's surface. Moving directions and distances were depicted by red arrows.

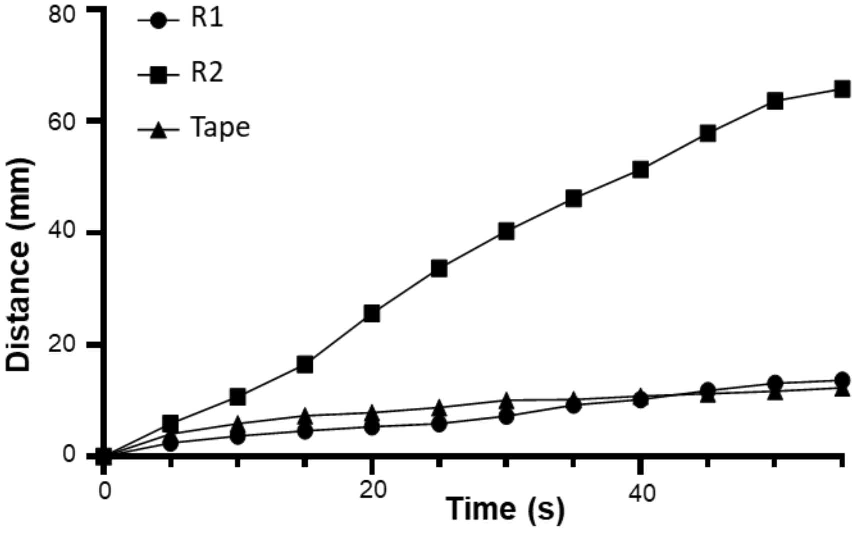

**Supplementary Figure S6. Migration distance of biosynthetic spidroin films in water.** The color tape traveling distance was recorded and calculated into overall distance in every five seconds.

**V. Analysis of protein structure of biosynthetic spidroin.**

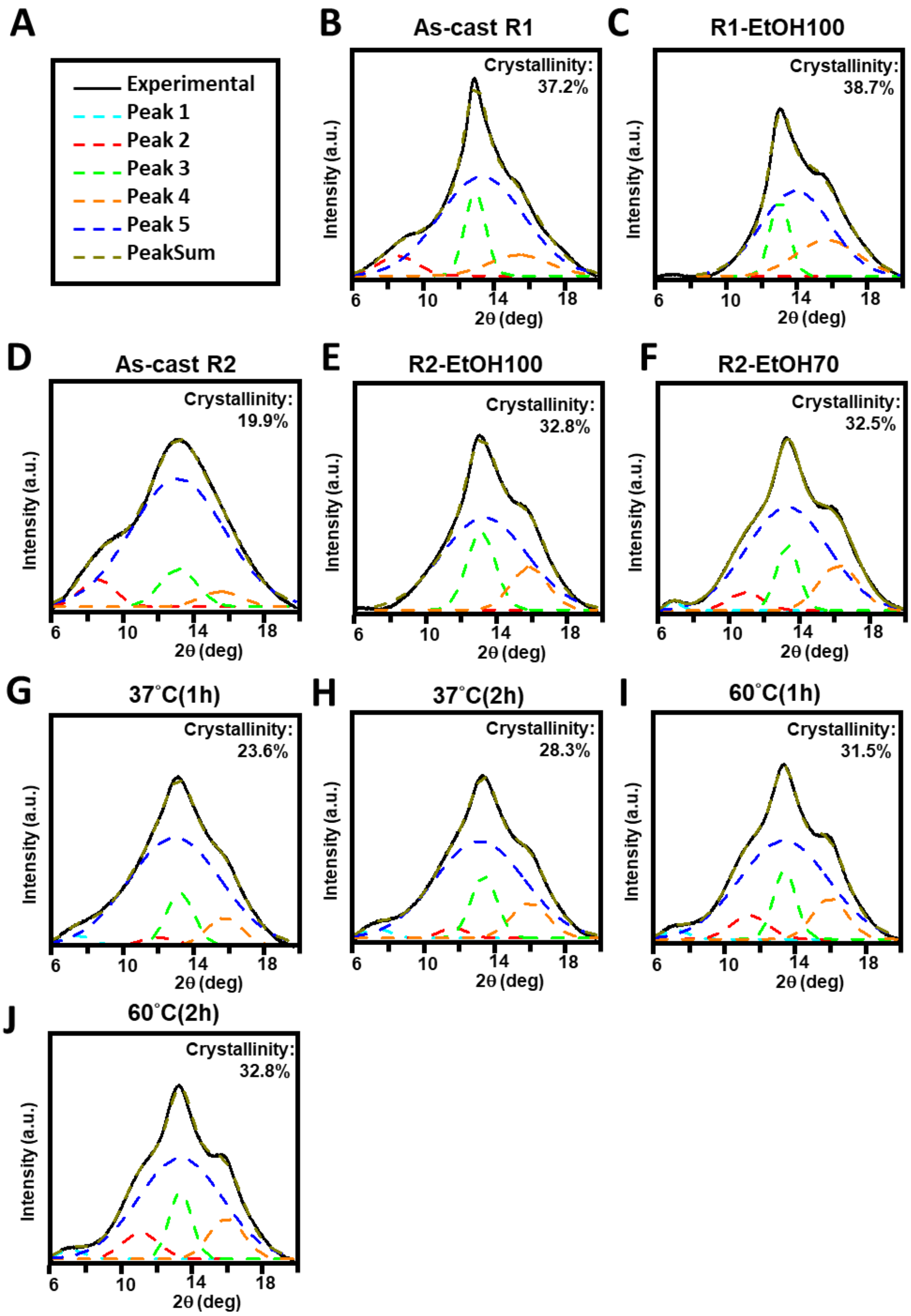

**Supplementary Figure S7. XRD structural characterization of biosynthetic spidroin films.** (A) The symbol of the line. Black line: Collected data; Light blue, red, and dark blue: Random coil; Green and orange: β-sheet; dark yellow: The integration of peaks. (B-J) Different treatment of spidroin. Each pattern was represented As-cast R1 (B), ETOH100 treated R1 (C), As-cast R2 (D), ETOH100 treated R2 (E), ETOH70 treated R2 (F), WVA 37°C (1h) for (G) and WVA 37°C (2h) for (H), 60°C WVA (1 h) for (I) and 60°C WVA (2 h) for (J).

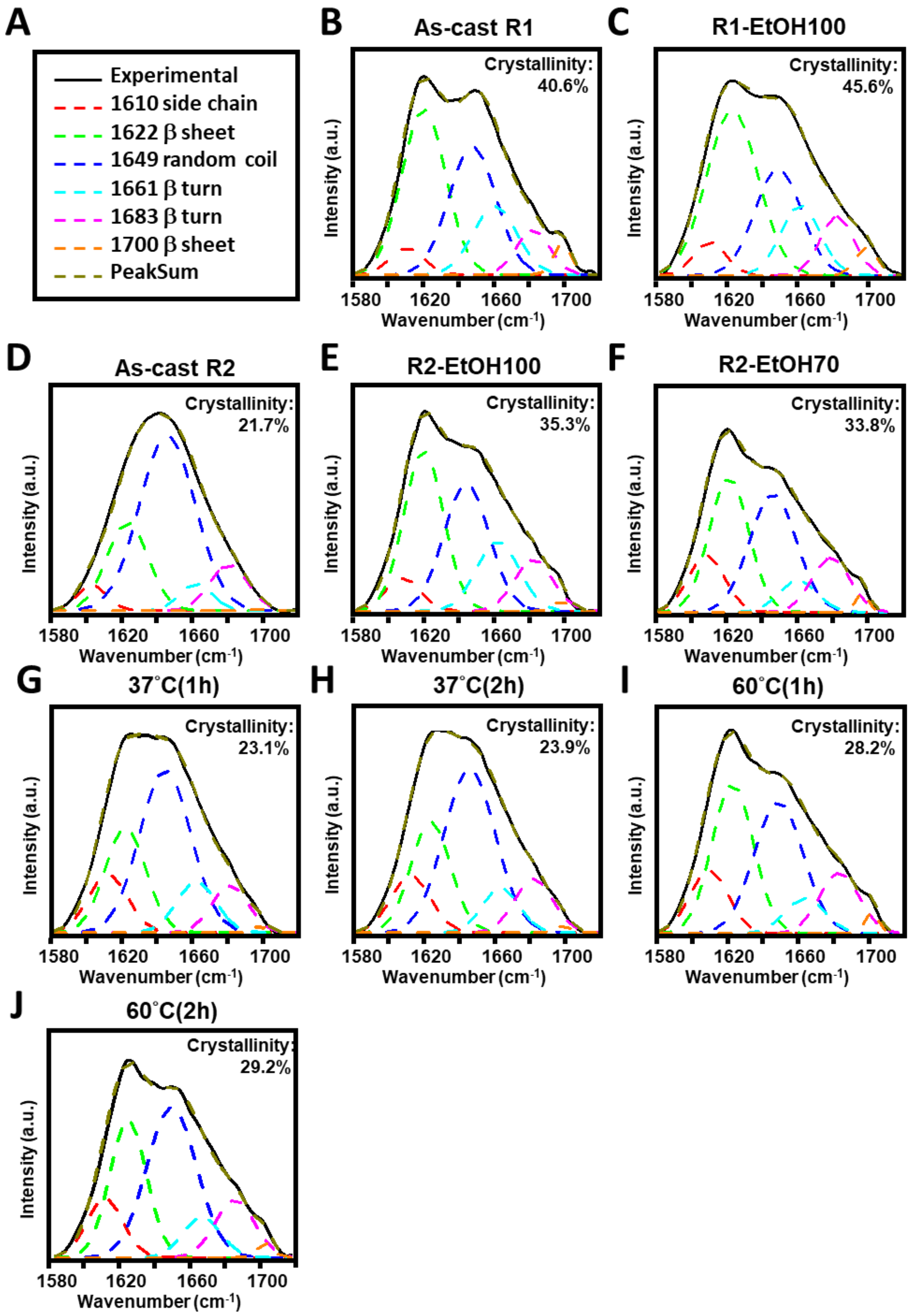

**Supplementary Figure S8. FTIR structural characterization of biosynthetic spidroin films.** (A) The symbol of the line. Black line: Collected data; Red dots: Side chain; Dark blue dots: Random coli; Light blue and pink dots: β-turn; Green and orange: β-sheet; dark yellow: The integration of peaks. (B-J) Different treatment of spidroin. Each pattern was represented As-cast R1 (B), ETOH100 treated R1 (C), As-cast R2 (D), ETOH100 treated R2 (E), ETOH70 treated R2 (F), WVA 37°C (1hr) for (G) and WVA 37°C (2h) for (H), 60°C WVA (1 h) for (I) and 60°C WVA (2 h) for (J).

**VI. Fabrication and mechanical characterization of novel graphene-doped spider silk materials**

Following the graphene-doped R2 film was created, the effects of graphene doping on the performance of R2 materials was examined. Both mechanical strength and cohesive repairing were evaluated. To characterize the strength variation in graphene-doped R2 (denoted R2G), mechanical properties of R2G films were conducted. Additionally, we also investigated the healability of the R2G as compared to R2 alone.

In the beginning, we compared non-treated conditions, as-cast (denoted as As) and cut-and-healed as-cast film (denoted as As-H) groups. The strength of graphene-contained R2 film was comparably similar ($p>0.05$). Next, we studied the 37 ° C WVA (2hr) film. The similar effect was observed in WVA groups, with no statistical difference in comparison to the control group. Also, the recovered strength of cut-and-healed R2G film was similar to the uncut one. It showed that graphene-doped R2 film had a good ability to self-repair, as comparable to the standard R2 films.

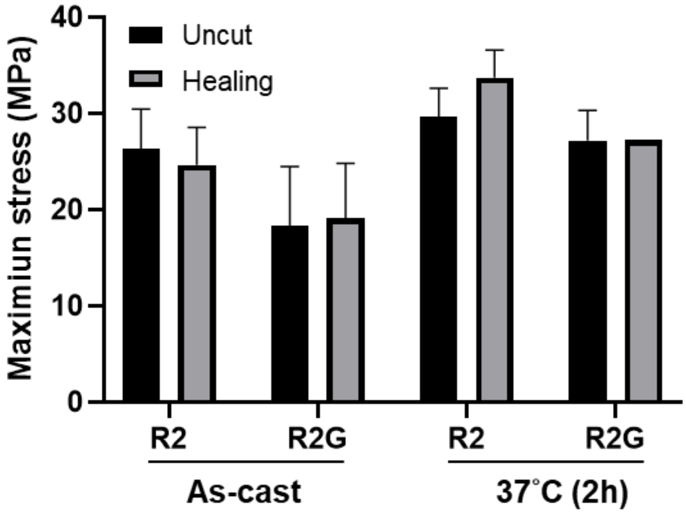

**Supplementary Figure S9. Tensile strength evaluation of graphene-doped biosynthetic spidroin films.** The maximum strength values of the as-cast R2 film and graphene-doped R2 film prior to and after the cut-and-healed processes were determined.

### VII. Repairable biosynthetic spidroin-based flexible logic gates

Based on R2G mechanical properties, the soft-fabrics were generated. Harnessing the spidroin-based materials, the complex fabrication process was also establised.

1D fabric demonstrated the healability of R2G and the logic gate table was shown in **Supplementary Fig. S10 and Supplementary Table S1.** For 2D fabric, combing R2G with R2 did significantly increase the complexity and utility of the system (**Supplementary Fig. S11A)**. The 2D fabric was installed in the device for the circuit evaluation (**Supplementary Fig. S11B**). The collected outputs were shown in **Supplementary Fig. S11C**. The output was determined by: Cutting, healing, graphene-dope R2 (Diverter), and electricity ($e^-$). Each panel was corresponding to specific operation in **Supplementary Fig. S11C** and the collective logic gate table was shown in **Supplementary Table S2**.

Next, 2D fabric was self-healed to create 3D fabric (**Supplementary Fig. S12A)**. The whole setup and the circuit diagram illustration were shown in **Supplementary Fig. S12B.** Besides the original 2D fabric properties, the variable for orientational alignment was introduced in 3D fabric. Specifically, tweaking, followed by the healing process can cause the misalignment of 3D structures, rendering a disconnected circuit. That is, the output was determined by: cutting, healing, tweaking, graphene-dope R2 (Diverter), and electricity ($e^-$). Each panel was corresponding to specific operation in **Supplementary Fig. S12C** and the final logic gate table was shown in **Supplementary Table S3**.

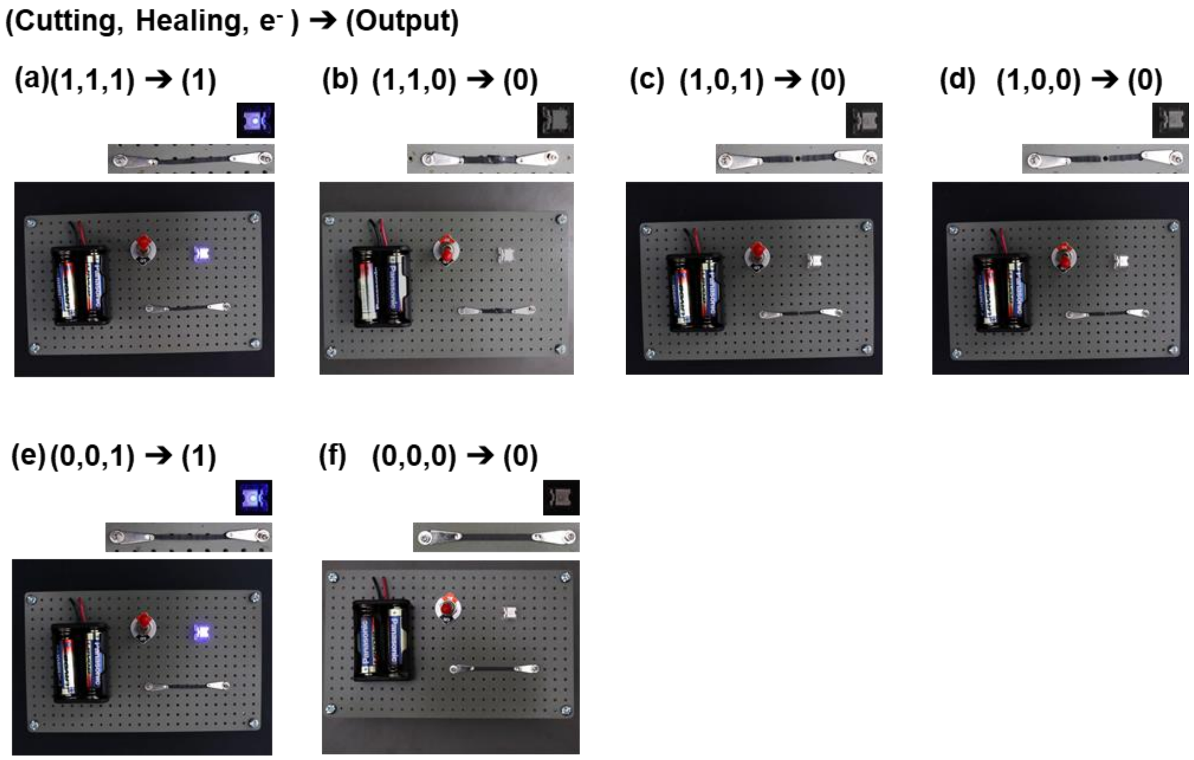

**Supplementary Figure S10. Repairable biosynthetic spidroin-based 1D logic gates.**

**Supplementary Table S1. Truth table of 1D biosynthetic spidroin-based logic gate circuitry.**

| Cutting | Healing | $e^-$ | Output |
|---|---|---|---|
| 1 | 1 | 1 | 1(a) |
| 1 | 1 | 0 | 0(b) |
| 1 | 0 | 1 | 0(c) |
| 1 | 0 | 0 | 0(d) |
| 0 | 1 | 1 | 1 |
| 0 | 1 | 0 | 0 |
| 0 | 0 | 1 | 1(e) |
| 0 | 0 | 0 | 0(f) |

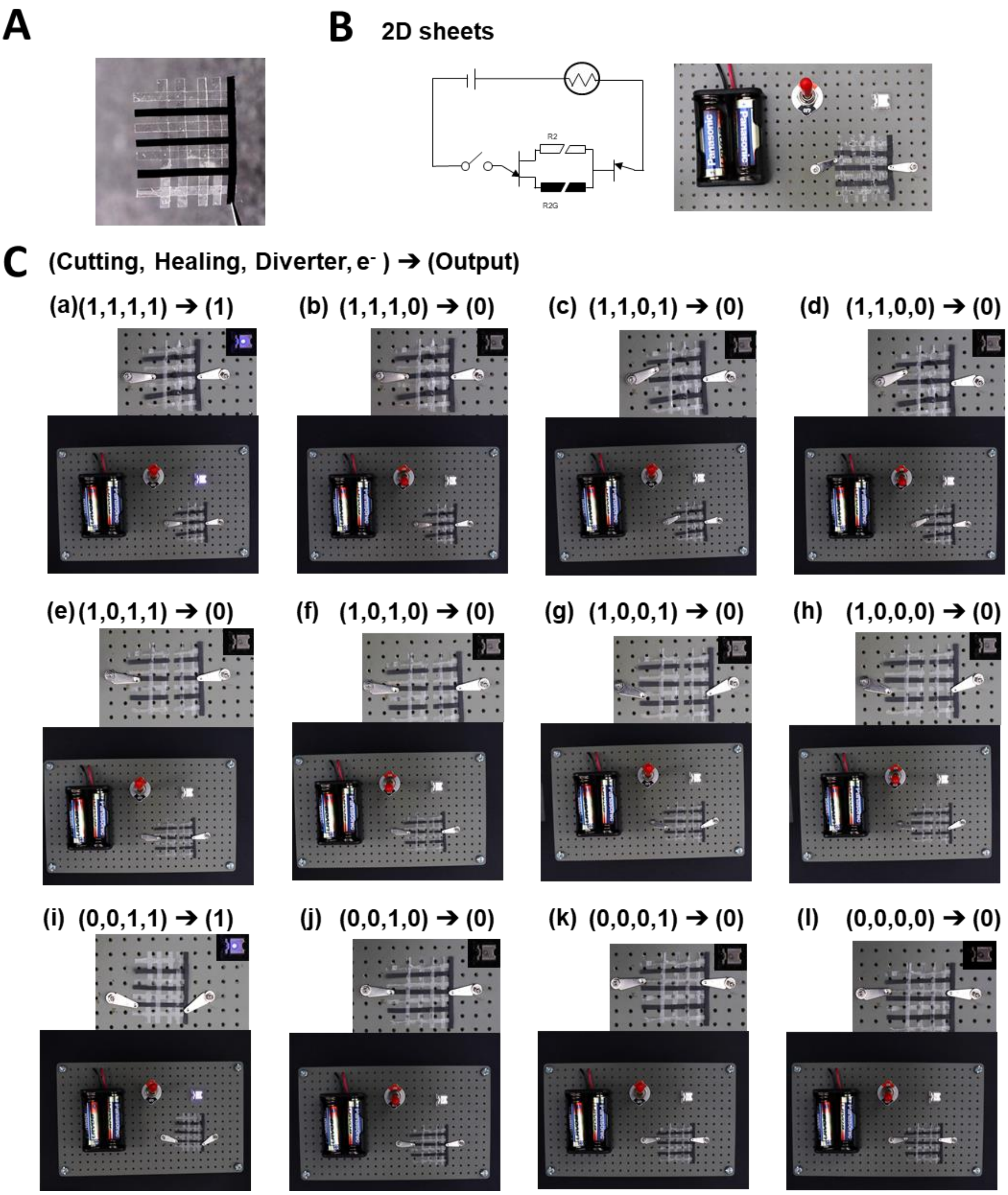

**Supplementary Figure S11. Repairable biosynthetic spidroin-based 2D logic gates.** (A) Fabrication of spider spidroin-based 2D sheet. (B) Setup and circuit diagram of 2D fabric system. (C) The operations and output of 2D fabric.

**Supplementary Table S2. Truth table of 2D biosynthetic spidroin-based logic gate circuitry.**

| Cutting | Healing | Diverter | $e^-$ | Output |
|---|---|---|---|---|
| 1 | 1 | 1 | 1 | 1 (a) |
| 1 | 1 | 1 | 0 | 0(b) |
| 1 | 1 | 0 | 1 | 0(c) |
| 1 | 1 | 0 | 0 | 0(d) |
| 1 | 0 | 1 | 1 | 0(e) |
| 1 | 0 | 1 | 0 | 0(f) |
| 1 | 0 | 0 | 1 | 0(g) |
| 1 | 0 | 0 | 0 | 0(h) |
| 0 | 1 | 1 | 1 | 1 |
| 0 | 1 | 1 | 0 | 0 |
| 0 | 1 | 0 | 1 | 0 |
| 0 | 1 | 0 | 0 | 0 |
| 0 | 0 | 1 | 1 | 1(i) |
| 0 | 0 | 1 | 0 | 0(j) |
| 0 | 0 | 0 | 1 | 0(k) |
| 0 | 0 | 0 | 0 | 0(l) |

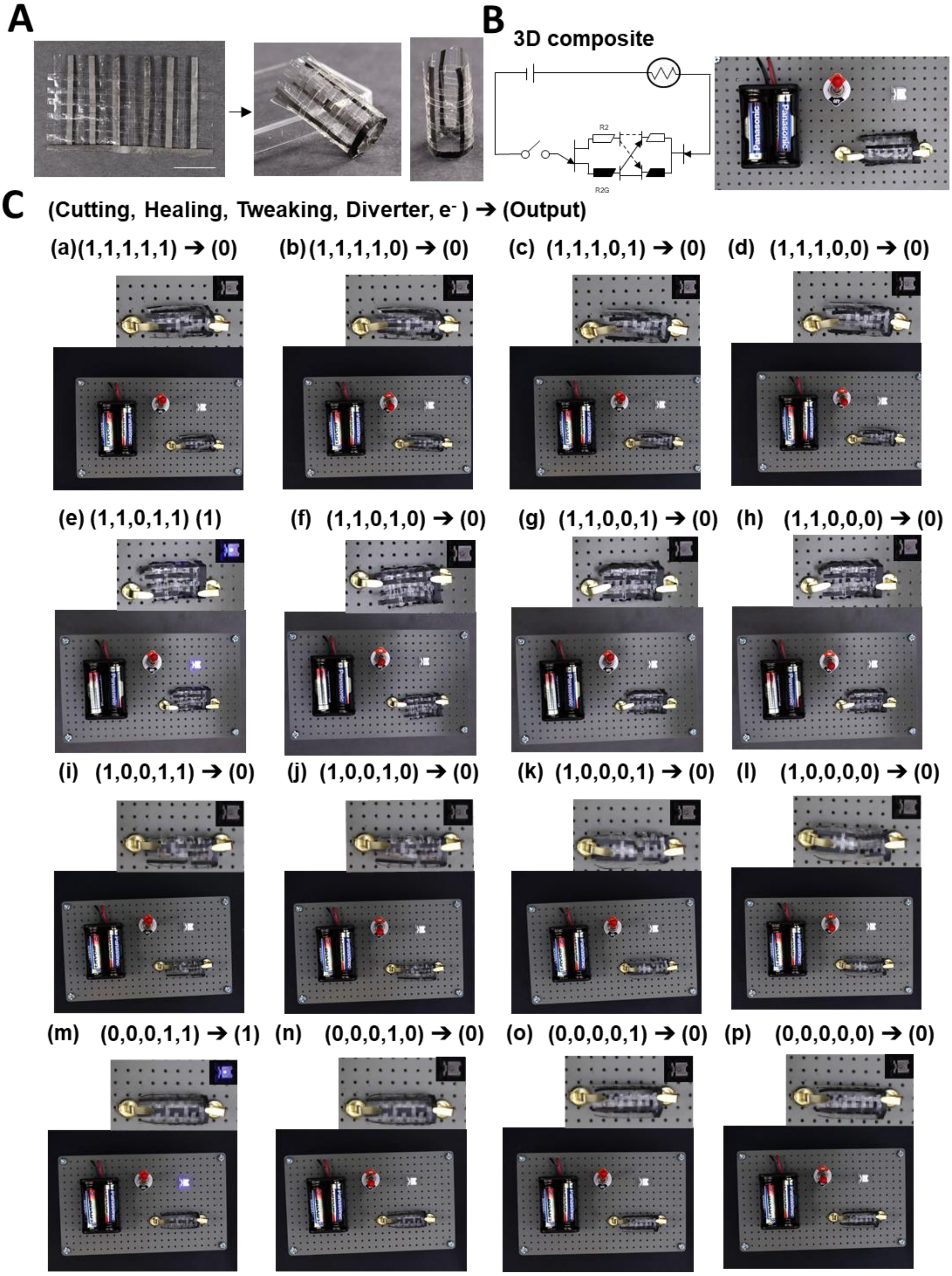

**Supplementary Figure S12. Repairable biosynthetic spidroin-based 3D logic gates.** (A) Fabrication of spider spidroin-based 3D fabric. (B) Setup and circuit diagram of 3D fabric. (C) The operations and output of 3D fabric.

**Supplementary Table S3. Truth table of 3D biosynthetic spidroin-based logic gate circuitry**

| Cutting | Healing | Tweaking | Diverter | $e^-$ | Output |
|---|---|---|---|---|---|
| 1 | 1 | 1 | 1 | 1 | 0 (a) |
| 1 | 1 | 1 | 1 | 0 | 0(b) |
| 1 | 1 | 1 | 0 | 1 | 0(c) |
| 1 | 1 | 1 | 0 | 0 | 0(d) |
| 1 | 1 | 0 | 1 | 1 | 1(e) |
| 1 | 1 | 0 | 1 | 0 | 0(f) |
| 1 | 1 | 0 | 0 | 1 | 0(g) |
| 1 | 1 | 0 | 0 | 0 | 0(h) |
| 1 | 0 | 1 | 1 | 1 | 0 |
| 1 | 0 | 1 | 1 | 0 | 0 |
| 1 | 0 | 1 | 0 | 1 | 0 |
| 1 | 0 | 1 | 0 | 0 | 0 |
| 1 | 0 | 0 | 1 | 1 | 0(i) |
| 1 | 0 | 0 | 1 | 0 | 0(j) |
| 1 | 0 | 0 | 0 | 1 | 0(k) |
| 1 | 0 | 0 | 0 | 0 | 0(l) |
| 0 | 1 | 1 | 1 | 1 | 0 |
| 0 | 1 | 1 | 1 | 0 | 0 |
| 0 | 1 | 1 | 0 | 1 | 0 |
| 0 | 1 | 1 | 0 | 0 | 0 |
| 0 | 1 | 0 | 1 | 1 | 1 |
| 0 | 1 | 0 | 1 | 0 | 0 |
| 0 | 1 | 0 | 0 | 1 | 0 |
| 0 | 1 | 0 | 0 | 0 | 0 |
| 0 | 0 | 1 | 1 | 1 | 1 |
| 0 | 0 | 1 | 1 | 0 | 0 |
| 0 | 0 | 1 | 0 | 1 | 0 |
| 0 | 0 | 1 | 0 | 0 | 0 |
| 0 | 0 | 0 | 1 | 1 | 1(m) |
| 0 | 0 | 0 | 1 | 0 | 0(n) |
| 0 | 0 | 0 | 0 | 1 | 0(o) |
| 0 | 0 | 0 | 0 | 0 | 0(p) |

**VIII. Device setup for the fabricated spider silk-based biorecognition sensor**

Harnessing reversible healability, WVA 60ºC (1h) R2, the materials were exploited to be integrated into a smart finger/skin recognition biodevice. Specifically, the R2G film was cut into small pieces and warped the wire on the R2-based ring (**Supplementary Fig. 7C**). The R2G film served as probes to detect the surface resistance of objects. Fingers' resistances were shown in **Supplementary Table S4** while electrical conductors' resistances were displayed in **Supplementary Table S5**. Insulator resistances were beyond the detection range of the Keithley 2400 source/meter. Consequently, the insulators' resistances were referred to Arc resistance [5] and was demonstrated in **Supplementary Table S6**.

Along resistance database, the spider silk-based biorecognition sensor was created. Biorecognition sensor was designed to identify the object and display the readout on the LCD. The Arduino sketch code was displayed in **Supplementary Fig. S13**. In the hardware design, interface of Arduino was represented in **Supplementary Fig. S14A** and complete coding was shown in **Supplementary Fig. S14B**. Finally, the biorecognition sensor was created in **Supplementary Fig. S14C**. The further test was depicted in **Supplementary Fig. 7E**.

**Supplementary Table S4. Resistance measurements of human fingers via the biosynthetic spidroin ring detector.**

| Person | Test area | Thumb (MΩ) | Forefinger (MΩ) | Middle finger (MΩ) | Ring finger (MΩ) | Little finger (MΩ) |
|---|---|---|---|---|---|---|
| 1 | Palm | 35 | 31 | 42 | 41 | 50 |
| | Back of hand | 50 | 73 | 84 | 40 | 61 |
| 2 | Palm | 46 | 48 | 42 | 51 | 33 |
| | Back of hand | 78 | 42 | 73 | 55 | 36 |
| 3 | Palm | 80 | 90 | 110 | 73 | 65 |
| | Back of hand | 67 | 58 | 78 | 65 | 90 |
| 4 | Palm | 23 | 35 | 33 | 32 | 47 |
| | Back of hand | 26 | 67 | 40 | 40 | 32 |
| 5 | Palm | 63 | 190 | 190 | 190 | 90 |
| | Back of hand | 190 | 170 | 198 | 70 | 50 |
| 6 | Palm | 45 | 85 | 91 | 47 | 70 |
| | Back of hand | 133 | 180 | 190 | 165 | 180 |
| 7 | Palm | 30 | 80 | 60 | 68 | 121 |
| | Back of hand | 150 | 65 | 81 | 73 | 42 |

**Supplementary Table S5. Resistance measurements of conductors s via the biosynthetic spidroin ring detector.**

| Material | Resistance (MΩ) |
|---|---|
| Iron | 1 |
| Copper | 2 |
| Silicon | 2 |
| Aluminum | 0.8 |
| Wire | 2 |

**Supplementary Table S6. Reference Resistances of insulator materials [5].**

| Material | Resistance (MΩ) |
|---|---|
| Bakelite | $10^{11}$ |
| Rubber | $10^{14}$ |
| acrylic | $10^{12}$ |
| Plastic | $10^{13}$ |

```
#include <LiquidCrystal_PCF8574.h>

LiquidCrystal_PCF8574 lcd(0x27);

void setup()
{
  lcd.begin(16, 2);
  lcd.setBacklight(255);
  lcd.clear();
  lcd.setCursor(0, 0);
  lcd.print("Resistance:");

} // setup()

void loop() {
  int sensorValue = analogRead(A0);
  float voltage = sensorValue * (5.0 / 1023.0);

  float R1= 330*1000000*(5/voltage-1);
  lcd.setCursor(2,1);
  if (R1 > 301*1000000)
  {
    lcd.print("OFF-HIGH");
  }
  else if ((300*1000000 > R1) && ( R1>10*1000000))
  {
    lcd.print("     ON       ");
  }
  else
  {
    lcd.print("OFF-LOW ");
  }Serial.println(R1);
}
```

**Supplementary Figure S13. Arduino sketch code of biosynthetic spidroin-based biorecognition sensor.**

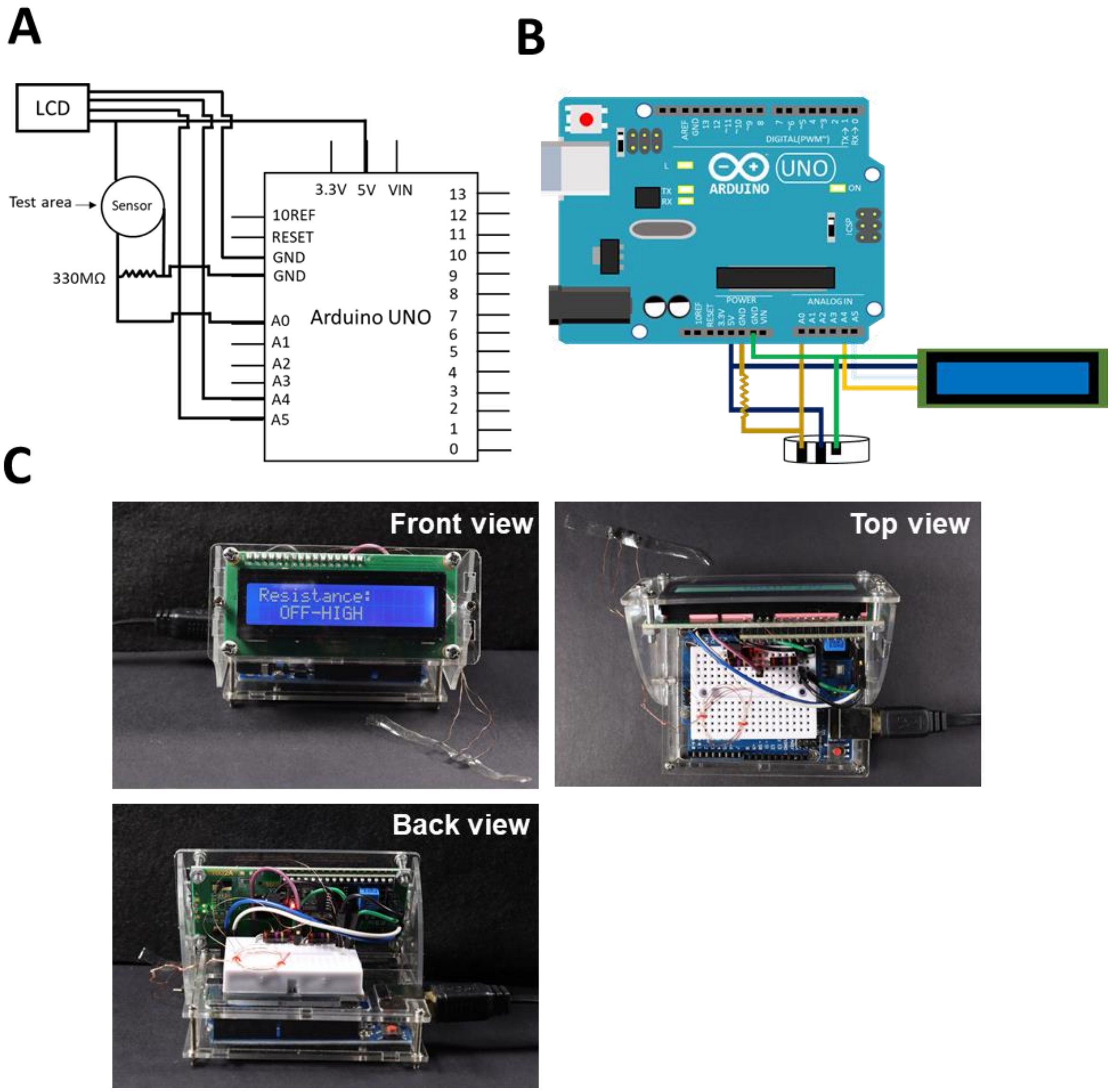

**Supplementary Figure S14. Device configuration for biosynthetic spidroin-based biorecognition sensor.** (A) Interface of the Arduino (B) Wiring diagram for spider silk-based biorecognition sensor (C) Device front view (top left), top view (top right), and back view (bottom)
